# Differential efficacies of Cas nucleases on microsatellites involved in human disorders and associated off-target mutations

**DOI:** 10.1101/857714

**Authors:** Lucie Poggi, Lisa Emmenegger, Stéphane Descorps-Declère, Bruno Dumas, Guy-Franck Richard

## Abstract

Microsatellite expansions are the cause of more than 20 neurological or developmental human disorders. Shortening expanded repeats using specific DNA endonucleases may be envisioned as a gene editing approach. Here, a new assay was developed to test several CRISPR-Cas nucleases on microsatellites involved in human diseases, by measuring at the same time double-strand break rates, DNA end resection and homologous recombination efficacy. Broad variations in nuclease performances were detected on all repeat tracts. *Streptococcus pyogenes* Cas9 was the most efficient of all. All repeat tracts did inhibit double-strand break resection. We demonstrate that secondary structure formation on the guide RNA was a major determinant of nuclease efficacy. Using deep sequencing, off-target mutations were assessed genomewide. Out of 221 CAG/CTG or GAA/TTC trinucleotide repeats of the yeast genome, three were identified as carrying statistically significant low frequency mutations, corresponding to off-target effects.

## Introduction

A growing number of neurological disorders were identified to be linked to microsatellite expansions (Orr and Zoghbi, 2007). Each disease is associated to a repeat expansion at a specific locus (Table 1). No cure exists for any of these dramatic disorders. Shortening the expanded array to non-pathological length could suppress symptoms of the pathology and could be used as a new gene therapy approach (Richard, 2015). Indeed, when a trinucleotide repeat contraction occurred during transmission from father to daughter of an expanded myotonic dystrophy type 1 allele, clinical examination of the daughter showed no sign of the disease (O’Hoy et al., 1993) (Shelbourne et al., 1992).

**Table 1.**
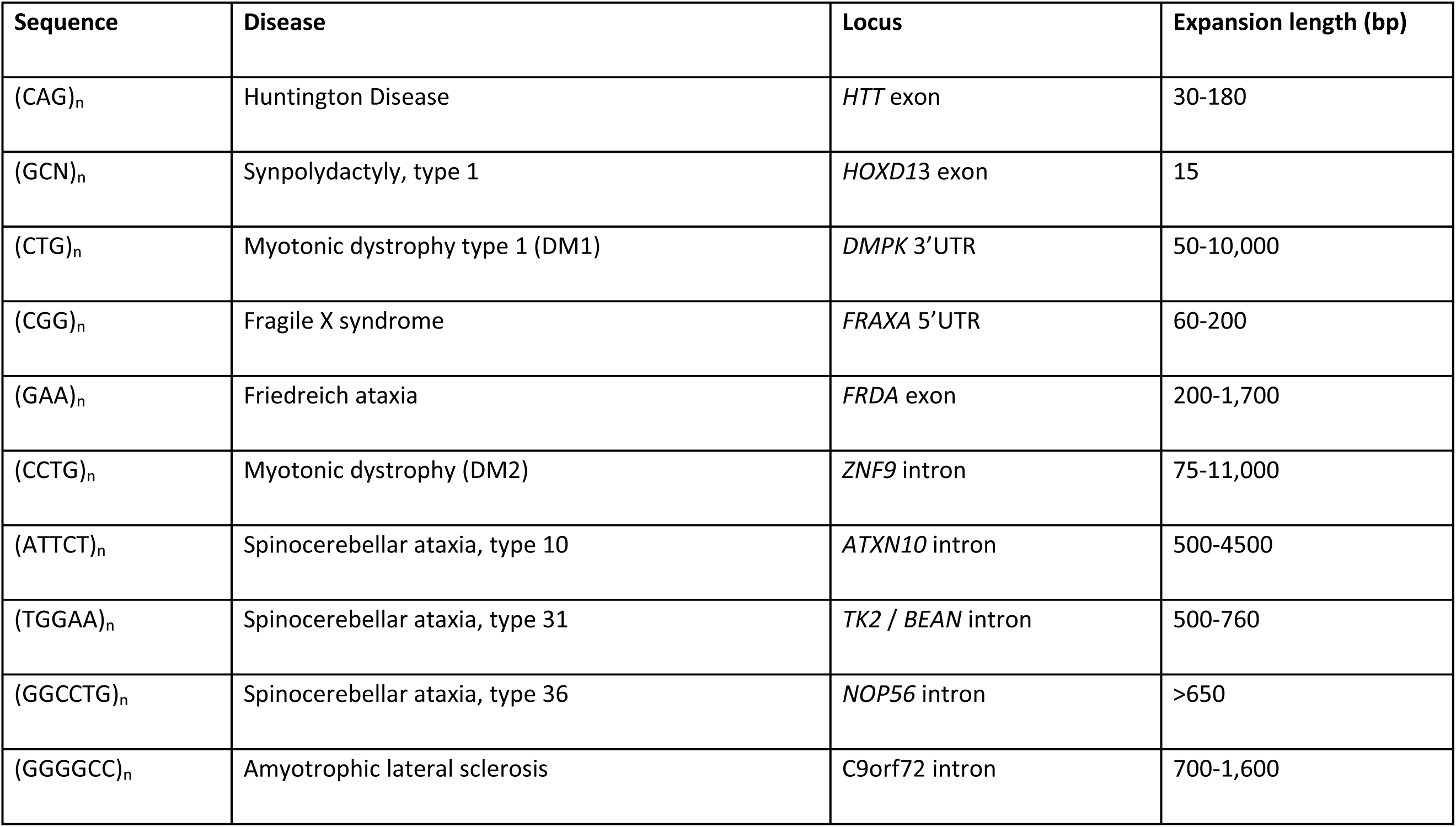
Summary of the main microsatellite disorders and associated repeat expansions.

In order to induce a double-strand break (DSB) into a microsatellite, different types of nucleases can be used: meganucleases, Zinc Finger Nucleases (ZFN), Transcription activator-like effector nucleases (TALEN) and CRISPR-Cas9. Previous experiments using the I-*Sce*I meganuclease to induce a DSB into a CTG repeat tract showed that repair occurred by annealing between the flanking CTG repeats (Richard et al., 1999). Later on, ZFNs were used to induce DSBs into CAG or CTG repeats, which mostly led to contractions in CHO cells (Mittelman et al., 2009) and in a HEK293 cell GFP reporter assay (Santillan et al., 2014). As only one arm was enough to induce a DSB into the repeat tract and since CAG zinc fingers can recognize CTG triplets and *vice versa*, the authors concluded that the specificity was too low for further medical applications. As a proof of concept of the approach, a myotonic dystrophy type 1 CTG repeat expansion was integrated into a yeast strain. A TALEN was designed to recognize and cut the CTG triplet repeat and was very efficient at shortening it in yeast cells (>99% cells showed contraction) and highly specific as no other mutation was detected (Richard et al., 2014). The TALEN was shown to induce specific repeat contractions through single-strand annealing (SSA) by a *RAD52*, *RAD50* and *SAE2* dependent mechanism (Mosbach et al., 2018). As a proof of concept of the approach, a myotonic dystrophy type 1 CTG repeat expansion was integrated into a yeast strain. A TALEN was designed to recognize and cut the CTG triplet repeat and was very efficient at shortening it in yeast cells (>99% cells showed contraction) and highly specific as no other mutation was detected (Richard et al., 2014).

The CRISPR-Cas system is the easiest to manipulate and to target any locus, as sequence recognition is based on the complementarity to a guide RNA (gRNA). To recognize its sequence, Cas9 requires a specific protospacer adjacent motif (PAM) that varies depending on the bacterial species of the Cas9 gene. The most widely used Cas9 is wild-type *Streptococcus pyogenes* Cas9 (SpCas9) (Cong et al., 2013). Its Protospacer Adjacent Motif (PAM) is NGG and induces a blunt cut 3-4 nucleotides away from it, through concerted activation of two catalytical domains, RuvC and HNH, each catalyzing one single-strand break (SSB). Issues were recently raised about the specificity of SpCas9, leading to the engineering of more specific variants. In eSpCas9, three positively charged residues interacting with the phosphate backbone of the non-target strand were neutralized, conferring an increased specificity (Kleinstiver et al., 2016). Similarly, Cas9-HF1 was mutated on 4 residues interacting through hydrogen bonds with the target strand (Slaymaker et al., 2016). *Staphylococcus aureus* is a smaller Cas9, its PAM is NNGRRT, having a similar structure to SpCas9 with two catalytic sites. Finally, type V CRISPR-Cas, Cpf1 nucleases, exhibit very different features including a T-rich PAM located 3’ of target DNA and making staggered cuts leaving five-nucleotide overhangs by iterative activation of a single RuvC catalytic site (Zetsche et al., 2015). Cutting repeated sequences like microsatellites may be difficult due to stable secondary structures that may form either on target DNA or on the guide RNA, making some repeats more or less permissive to nuclease recognition and cleavage. In addition, secondary structure formation could impede DSB resection or later repair steps. Eukaryotic genomes contain thousands of identical microsatellites, therefore the specificity issue may become a real problem when targeting one single locus. Here we developed an *in vivo* assay in the yeast *Saccharomyces cerevisiae* in order to test different nucleases belonging to the CRISPR-Cas family on synthetic microsatellites associated to human disorders. Our experiments revealed that these sequences may be cut, with surprisingly different efficacies between nucleases and between microsatellites. SpCas9 was the most efficient and nuclease efficacy relied mainly on gRNA stability, strongly suggesting that secondary structures are the limiting factor in inducing a DSB *in vivo.* DSB resection was decreased to different levels in all repeated tracts. In addition, we analyzed off-target mutations genomewide and found that three microsatellites with similar sequences were also edited by the nuclease. The mutation pattern was different depending on the microsatellite targeted.

## Materials and Methods

### Yeast plasmids

A synthetic cassette (synYEGFP) was ordered from ThermoFisher (GeneArt). It is a pUC57 vector containing upstream and downstream *CAN1* homology sequences flanking a bipartite eGFP gene interrupted by the I-*Sce* I recognition sequence (18 bp) under the control of the *TEF1* promoter and followed by the *CYC1* terminator. The *TRP1* selection marker along with its own promoter and terminator regions was added downstream the eGFP sequences (Figure 1A). The I-*Sce* I site was flanked by *Sap* I recognition sequences, in order to clone the different repeat tracts. Nine out the 10 repeat tracts were ordered from ThermoFisher (GeneArt) as 151 bp DNA fragments containing 100 bp of repeated sequence flanked by *Sap* I sites. The last repeat (GGGGCC) was ordered from Proteogenix. All these repeat tracts were cloned at the *Sap* I site of synYEGFP by standard procedures, to give plasmids pLPX101 to pLPX110 (Supplemental Table S1). All nucleases were cloned in a centromeric yeast plasmid derived from pRS415 (Sikorski and Hieter, 1989), carrying a *LEU2* selection marker. Each open reading frame was placed under the control of the GalL promoter, derived from *GAL10*, followed by the *CYC1* terminator (DiCarlo et al., 2013).These plasmids were cloned directly into yeast cells by homology-driven recombination (Muller et al., 2012) using 34-bp homology on one side and 40-bp homology on the other and were called pLPX10 to pLPX16. Primers used to amplify each nuclease are indicated in Supplemental Table S2. Nucleases were amplified from Addgene plasmids indicated in Supplemental Table S1. The I-*Sce* I gene was amplified from pTRi103 (Richard et al., 2003).Guide RNAs for SpCas9 (and variants) were ordered from ThermoFisher (GeneArt), flanked by *Eco* RI sites for subsequent cloning into pRS416 (Sikorski and Hieter, 1989). SaCas9 and FnCpf1 guide RNAs were ordered at Twist Biosciences, directly cloned into pRS416 (see Supplemental Table S1 for plasmid names). Each guide RNA was synthetized under the control of the *SNR52* promoter.

**Figure 1:**
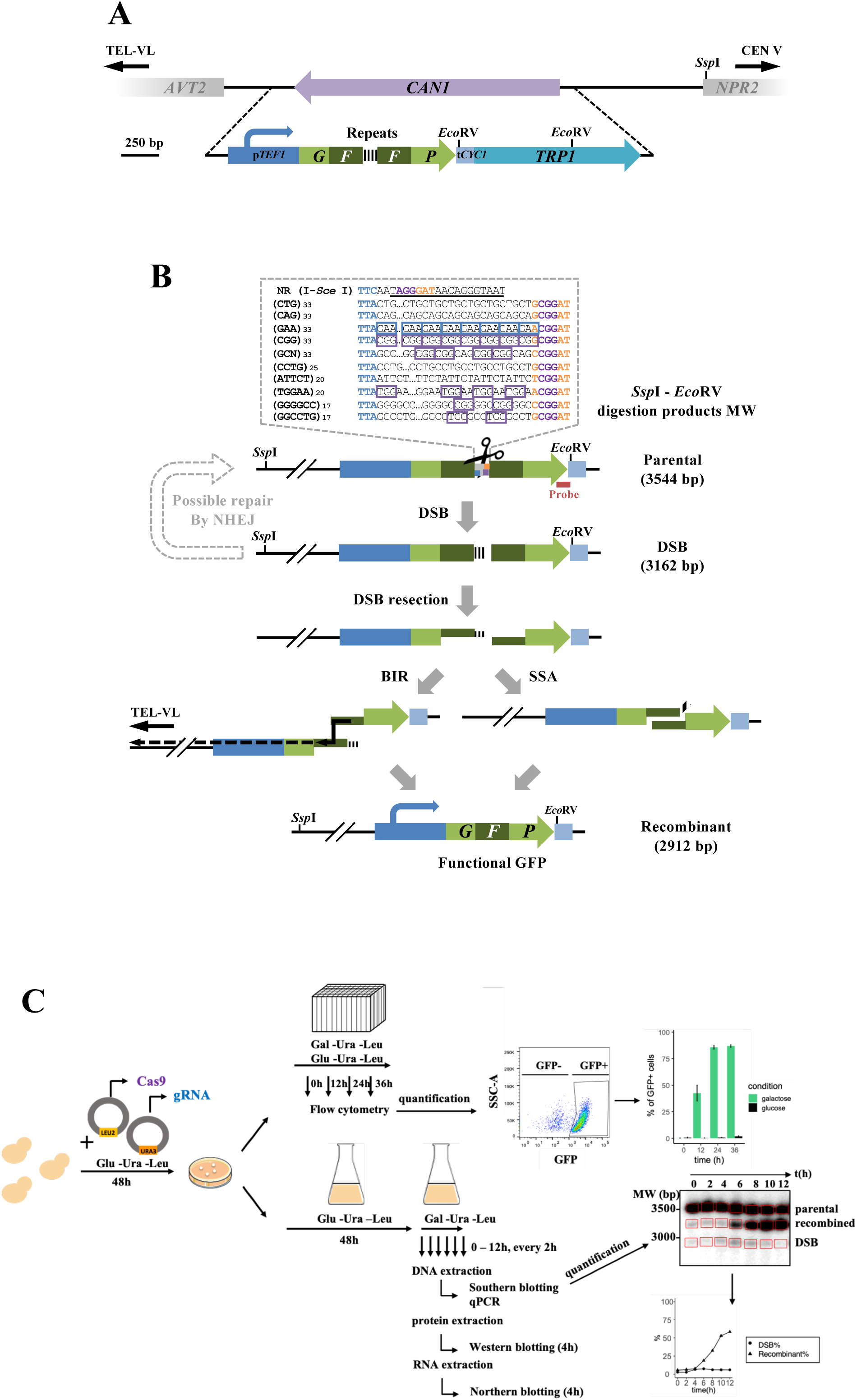
GFR reporter assay. **A:** The *CAN1* locus was replaced by recombinant GFP cassettes. Each synthetic GFP contains the constitutive *TEF1* promoter, followed by the bipartite eGFP interrupted by one of the ten microsatellites or by the I-*Sce* I recognition sequence and the *CYC1* terminator. The *TRP1* gene was used as an auxotrophic marker. **B**: Upon double-strand break induction, cells may repair by single-strand annealing, break-induced replication to reconstitute a functional GFP. Alternatively, repair by NHEJ may occur but will never lead to functional GFP. PAM sequences common to all constructs are colored (orange: SaCas9, purple: SpCas9, blue: FnCpf1) and additional PAM are boxed (same color code). *Ssp* I and *Eco* RV restriction site positions are indicated. Predicted molecular weights of each molecular species are indicated on the right. **C:** Cartoon depicting the experimental protocol (see text). Dot-plot axes are FSC-A/SSC-A.

### Yeast strains

Each synYEGFP cassette containing repeat tracts was digested by *Bam* HI in order to linearize it and transformed into the FYBL1-4D strain (Gietz et al., 1995). Correct integrations at the *CAN1* locus were first screened as [CanR, Trp+] transformants, on SC - ARG -TRP +Canavanine (60 μ/ml) plates. Repeats were amplified by PCR using LP30b-LP33b primers and sequenced (Eurofins/GATC). As a final confirmation, all transformants were also analyzed by Southern blot and all the [CanR, Trp+] clones showed the expected profile at the *CAN1* locus. Derived strains were called LPY101 to LPY111 (Supplemental Table S3).

### Flow cytometry assay

Cells were transformed using standard lithium-acetate protocol (Gietz et al., 1995) with both guide and nuclease and selected on 2% glucose SC -URA -LEU plates and grown for 36 hours. Each colony was then picked and seeded into a 96-well plate containing 300 μL of either 2% glucose SC -URA -LEU or 2% galactose SC -URA -LEU. At each time point (0h, 12h, 24h, 36h) cells were diluted in PBS and quantified by flow cytometry after gating on homogenous population, single cells and GFP-positive cells. The complete protocol was extensively described in (Poggi et al., 2020).

### Time courses of DSB inductions

Cells were transformed using standard lithium-acetate protocol (Gietz et al., 1995) with both guide and nuclease and selected on 2% glucose SC -URA -LEU plates and grown for 36 hours. Each colony was seeded into 2 mL of 2% glucose SC -URA -LEU for 24 hours and then diluted into 10 mL of 2% glucose SC -URA -LEU for 24 hours as a pre-culture step. Cells were washed twice in water and diluted at ca. 7 x 10^6^ cells/mL in 2% galactose SC-URA -LEU, before being harvested at each time point (0h, 2h, 4h, 6h, 8h, 10h, 12h) for subsequent DNA extractions.

### Southern blot analyses

For each Southern blot, 3-5 μg of genomic DNA digested with *Eco* RV and *Ssp* I were loaded on a 1% agarose gel and electrophoresis was performed overnight at 1V/cm. The gel was manually transferred overnight in 20X SSC, on a Hybond-XL nylon membrane (GE Healthcare), according to manufacturer recommendations. Hybridization was performed with a ^32^P-randomly labeled *CAN1* probe amplified from primers CAN133 and CAN135 (Supplemental Table S2) (Viterbo et al., 2018). The membrane was exposed 3 days on a phosphor screen and quantifications were performed on a FujiFilm FLA-9000 phosphorimager, using the Multi Gauge (v. 3.0) software. Percentages of DSB and recombinant molecules were calculated as the amount of each corresponding band divided by the total amount of signal in the lane.

### Agarose plug DNA preparation

During time courses of DSB induction (see above), 2 x 10^9^ cells were collected at each time point and centrifuged. Each pellet was resuspended in 330 μL 50 mM EDTA (pH 9.0), taking into account the pellet volume. Under a chemical hood, 110 μL of Solution I (1 M sorbitol, 10 mM EDTA (pH 9.0), 100 mM sodium citrate (pH 5.8), 2.5% β-mercaptoethanol and 10 μL of 100 mg/mL Zymolyase 100T-Seikagaku) were added to the cells, before 560 mL of 1% InCert agarose (Lonza) were delicately added and mixed. This mix was rapidly poured into plug molds and left in the cold room for at least 10 minutes. When solidified, agarose plugs were removed from the molds and incubated overnight at 37°C in Solution II (450 mM EDTA (pH 9.0), 10 mM Tris-HCl (pH 8.0), 7.5% β-mercaptoethanol). In the morning, tubes were cooled down on ice before Solution II was delicately removed with a pipette and replaced by Solution III (450 mM EDTA (pH 9.0), 10 mM Tris-HCl (pH 8.0), 1% N-lauryl sarcosyl, 1 mg/mL Proteinase K). Tubes were incubated overnight at 65°C, before being cooled down on ice in the morning. Solution III was removed and replaced by TE (10 mM Tris pH 8.0, 1 mM EDTA). Blocks were incubated in 1 mL TE in 2 mL eppendorf tubes for one hour at 4°C, repeated four times. TE was replaced by 1 mL restriction enzyme buffer (Invitrogen REACT 2) for one hour, then replaced by 100 μl buffer containing 100 units of each enzyme (*Eco* RV and *Ssp* I) and left overnight at 37°C. Agarose was melted at 70°C for 10 minutes without removing the buffer, 100 units of each enzyme *Eco* RV and *Ssp* I was added and left at 37°C for one hour. Then, 2 μL of β-agarase (NEB M0392S) and 2 μL of RNAse A (Roche 1 119 915) were added and left for one hour at 37°C. Microtubes were centrifuged at maximum speed in a tabletop centrifuge for one minute to pellet undigested agarose. The liquid phase was collected with wide bore 200 μL filter tips (Fisher Scientific #2069G) and loaded on a 1% agarose gel, subsequently processed as for a regular Southern blot (see above).

### Northern blot analyses

Each repeat-containing strain transformed with its cognate gRNA and nucleases was grown for 4 hours in 2% galactose SC-URA-LEU. Total RNAs were extracted using standard phenol-chloroform procedure (Richard et al., 1997) or the miRVANA kit, used to extract very low levels of small RNAs with high efficacy (ThermoFisher). Total RNA samples were loaded on 50% urea 10% polyacrylamide gels and run at 20 W for one hour. Gels were electroblotted on N^+^ nylon membranes (GE Healthcare), hybridized at 42°C using a SpCas9, SaCas9, FnCpf1 or *SNR44* oligonucleotidic probe. Each probe was terminally labeled with γ-^32^P ATP in the presence of polynucleotide kinase.

### Western blot analyses

Total proteins were extracted in 2X Laemmli buffer and denatured at 95°C before being loaded on a 12% polyacrylamide gel. After migration, the gel was electroblotted (0.22 A, constant voltage) on a Nytran membrane (Whatman), blocked for one hour in 3% NFDM/TBS-T and hybridized using either anti-SpCas9 (ab202580, dilution 1/1000), anti-SaCas9 (ab203936, dilution 1/1000), anti-HA (ab9110, dilution 1/1000) or anti-ZWF1 (A9521, dilution 1/100 000) overnight. Membranes were washed in TBS-T for 10 minutes twice. Following anti-SaCas9 and anti-HA hybridization, a secondary hybridization using secondary antibody Goat anti-Rabbit 31460 (dilution 1/5000). Membranes were read and quantified on a Bio-Rad ChemiDoc apparatus.

### Analysis of DSB end resection

A real-time PCR assay using primer pairs flanking *Sty* I sites 282 bp away from 5’ end of the repeat sequence and 478 bp away from the 3′ end of the repeat tract (LP001/LP002 and LP003/LP004, respectively) was used to quantify end resection. Another pair of primers was used to amplify a region of chromosome X to serve as an internal control of the DNA amount (JEM1f-JEM1r). Genomic DNA of cells collected at t = 12h was split in two fractions; one was used for *Sty* I digestion and the other one for a mock digestion in a final volume of 15 μL. Samples were incubated for 5h at 37°C and then the enzyme was inactivated for 20 min at 65°C. DNA was subsequently diluted by adding 55 μL of ice-cold water, and 4 μL was used for each real-time PCR reaction in a final volume of 25 μL. PCRs were performed with EurobioProbe qPCR Mix Lo-ROX in a CFX96 Real time machine (Bio-Rad) using the following program: 95°C for 15 min, 95°C for 15 s, 55°C for 30 s, and 72°C for 30 s repeated 40 times, followed by a 20-min melting curve. Reactions were performed in triplicate, and the mean value was used to determine the amount of resected DNA using the following formula: raw resection = 2/(1+2ΔCt) with ΔC_t_ = C_t,StyI_−C_t,mock_. Relative resection values were calculated by dividing raw resection values by the percentage of DSB quantified at the corresponding time point (Chen et al., 2013). Ratios of relative resection rates from both sides of the repeated sequence were calculated and compared to a non-repeated control sequence.

### Determination of off-target mutations

Cell were grown overnight in YPGal medium and diluted for 2 more hours. Cells were incubated in 20ml 0.1M of Lithium Acetate/TE buffer for 45 minutes at 30°C. 500 µl of 1M DTT was added and cells were incubated for a further 15 minutes at the same temperature. Cells were washed in water, then in 1M Sorbitol and resuspended in 120 µl ice-cold 1M Sorbitol. Then, 40 µl of competent cells were mixed with 150 ng of guide RNA-expressing plasmid, 300 ng of nuclease-expressing plasmid and 100 µM of dsODN (5′-PG*T*TTAATTGAGTTGTCATATGTTAATAACGGT*A*T-3’; where P represents a 5′ phosphorylation and * indicates a phosphorothioate linkage). Cells were electroporated at 1.5 kV, 25 μF, 200 Ω. Right after electroporation (BioRad Micropulser), 1 ml 1M Sorbitol was added to the mixture. Cells were centrifuged and supernatant was removed to plate a volume of 200 µl on 2% galactose SC -URA -LEU plates and grown for 72 hours. Negative control consisted of the same procedure without dsODN. Genomic DNA was extracted and approximately 10 μg of total genomic DNA was extracted and sonicated to an average size of 500 bp, on a Covaris S220 (LGC Genomics) in microtubes AFA (6×16 mm) using the following setup: Peak Incident Power: 105 Watts, Duty Factor: 5%, 200 cycles, 80 seconds. DNA ends were subsequently repaired with T4 DNA polymerase (15 units, NEBiolabs) and Klenow DNA polymerase (5 units, NEBiolabs) and phosphorylated with T4 DNA kinase (50 units, NEBiolabs). Repaired DNA was purified on two MinElute columns (Qiagen) and eluted in 16 μl (32 μl final for each library). Addition of a 3’ dATP was performed with Klenow DNA polymerase (exo-) (15 units, NEBiolabs). Home-made adapters containing a 4-bp unique tag used for multiplexing, were ligated with 2 μl T4 DNA ligase (NEBiolabs, 400,000 units/ml). DNA was size fractionated on 1% agarose gels and 500-750 bp DNA fragments were gel extracted with the Qiaquick gel extraction kit (Qiagen). A first round of PCR was performed using primers GSP1 and P1 (First denaturation step: 98°C for 30s; 98°C for 30s, 50°C for 30s, 72°C for 30s repeated 30 times; followed by 72°C for 7min). A second round of PCR was performed using primers PE1 and GSP2-PE2 (First denaturation step: 98°C for 30s; 98°C for 30s, 65°C for 30s, 72°C for 30s repeated 30 times; followed by 72°C for 7min) A final round of PCR was performed using primers PE1 and PE2 (First denaturation step: 98°C for 30s; 98°C for 30s, 65°C for 30s, 72°C for 30s repeated 15 times; followed by 72°C for 7min). Libraries were purified on agarose gel and quantified on a Bioanalyzer. Equimolar amounts of each library were loaded on a Next-Seq Mid output flow cell cartridge (Illumina NextSeq 500/550 #20022409).

### Computer analysis of off-target mutations

In a first step, all fastq originating from the different libraries were scanned in order to identify reads coming from the dsODN specific amplification. The test was carried out with the standard unix command grep and the result was used to split each former fastq file in two: with or without the dsODN tag. In a second step, all the resulting fastq files were mapped against the S288C reference genome obtained from the SGD database (release R64-2-1_20150113, https://www.yeastgenome.org/). Mapping was carried out by minimap2 (Li, 2018) using “-ax sr --secondary=no” parameters. Sam files resulting from mappings were then all sorted and indexed by the samtools software suite (Li et al., 2009). Subsequently, for dsODN-containing sequences, double strand break positions were identified by searching coverage peaks. Peaks were defined as region showing a coverage at least equal to twice the median coverage. Regarding reads that did not contain the dsODN tag, mutations within predicted off-target sites were detected by the mean of samtools pileup applied to all regions of interest identified by crispor (Haeussler et al., 2016). Each of the 56 positions exhibiting mutations was manually examined using the IGV visualization software and validated or not, as explained in the text.

### Analysis of Gibbs free energy for gRNA guide sequences

The Gibbs free energy formation for gRNA secondary structures were determined using the MFOLD RNA 2.3 (http://unafold.rna.albany.edu/?q=mfold/RNA-Folding-Form2.3) with temperature parameter set to 30°C.

### Statistical Analysis

All statistical tests were performed with R3.5.1. Linear regression model was performed to test the correlation between DSB value and the percentage of GFP-positive cells at different time points. Linear regression was performed to determine statistical significance of proteins levels and gRNA levels over the percentage of GFP-positive cells. For each linear regression, R^2^ and p-value were calculated. One-way analysis of variance (ANOVA) was used to determine the impact of gRNA free energy over the percentage of GFP-positive cells at 36 h. P-values less than 0.05 were considered significant. Figures were plotted using the package ggplot2.

## Results

### A GFP reporter assay integrated in the *Saccharomyces cerevisiae* genome enables the quantification of nuclease activity

The goal of the present experiments was to design and build a reporter system in the yeast *S. cerevisiae* to determine efficacy and specificity of different Cas nucleases on various microsatellites. In order to accurately compare experiments, we decided to use synthetic microsatellites integrated at the same position in the yeast genome. The advantage of this approach -as compared to using the human repeat tract sequences- was that all nucleases could be tested on the same genomic and chromatinian environment. In addition, we made the synthetic constructs in such a way that PAM sequences were available to each nuclease, which was not possible with human sequences. We therefore built a set of 11 isogenic yeast strains, differing only by the repeat sequence cloned in a cassette containing two synthetic GFP halves flanking 100 bp-repeats, integrated at the same genomic locus and replacing the *CAN1* gene on yeast chromosome V (Figure 1A). Note that given the repeated nature of the target DNA, some of them also harbor internal PAM sequences (Figure 1B).

Upon DSB induction, haploid yeast cells may fix the break by three different pathways. Homology regions flanking the DSB site may be used to repair the DSB either by single-strand annealing (SSA) between the two GFP halves, or by break-induced replication (BIR) to the end of the chromosome. In both cases, a fully functional GFP gene will be reconstituted. Note that in our experimental system, we cannot distinguish between BIR and SSA events. Alternatively, the DSB may be repaired by end-joining (NHEJ) between the two DNA ends. However, this is very unlikely, since NHEJ is downregulated in haploid cells (Valencia et al., 2001). In any case, perfectly religated DSB ends could be recut by the nuclease, until a functional GFP could be reconstituted by homologous recombination.

All experiments were performed as follows: independent yeast colonies expressing each Cas nuclease and its cognate gRNA were picked from glucose plates and seeded either in 96-deep well plates for flow cytometry measurements over a 36-h time period. Simultaneously, a colony from the same strain was expanded in a 500 mL flask to recover sufficient cells for further molecular analyses (Figure 1C). As a control in all experiments, we used a non-repeated sequence containing the I-*Sce*I recognition site.

By flow cytometry, two distinct populations separated by one or two fluorescence intensity logarithms, corresponding to GFP-negative and GFP-positive cells, were observed upon nuclease induction (Figure 2A). Tested nucleases showed very different efficacies, SpCas9, FnCpf1 and SaCas9 were all more efficient than I-*Sce*I itself, as indicated by a higher number of GFP-positive cells. In order to know whether GFP-positive cells were a good readout of DSB efficacy, Southern blots were performed to detect and quantify parental and recombinant products as well as the DSB. A time course was run over a 12-hour period of time for each strain and each nuclease (except for the N863A Cas9 nickase). Parental, recombinant and DSB signals were quantified using phosphorimaging technology. In all cases, the DSB and recombinant products were detected, although in variable amounts (Figure 2B). The only exceptions were Cas9-HF1 in which no DSB nor recombinant band were detected, and Cas9-D10A in which a faint DSB signal was recorded but no recombinant molecules could be seen (see later). In subsequent experiments, the I-*Sce* I sequence will be used as our reference and called ‘NR’ (for Non Repeated).

**Figure 2:**
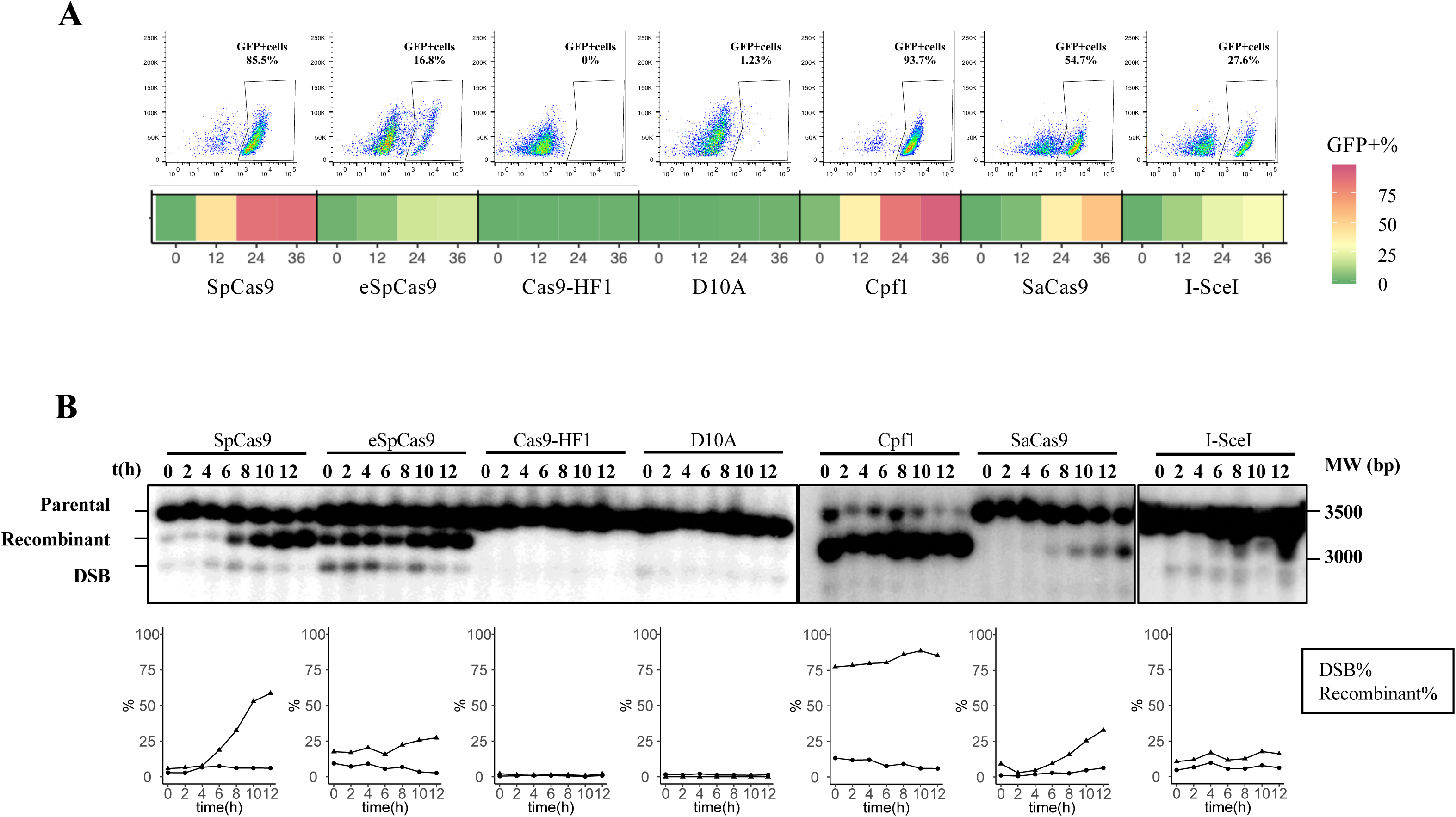
CRISPR-Cas nuclease induction on non-repeated sequence containing an I-*Sce* I recognition site. **A:** Top: Percentage of GFP+ cells was measured throughout a time course of 36 hours. Dot plots indicate final populations at 36 hours. X-axis: FITC, Y-axis: SSC. Bottom: GFP+ cells are represented by a color code: from low recombination rates in dark green to high recombination in dark red. **B:** Top: Repair time courses were carried out during 12 hours. Parental (3500 bp), recombinant (3100 bp) and DSB (2900 bp) products were quantified (see Materials & Methods). Bottom: DSB and recombinant products are represented as a percentage of the total signal in each lane.

### *Streptococcus pyogenes* Cas9 variants exhibit a wide range of efficacies

Once the experimental setup was optimized with I-*Sce* I (NR), the same exact assay was performed using seven different nucleases on ten microsatellites, including tri-, tetra-, penta- and hexanucleotide repeats (Supplemental Figures S1 & S2 and Supplemental Table S4). SpCas9 was able to cut every repeated sequence although (GGCCTG)_15_, (GAA)_33_, (CAG)_33_ and (CTG)_33_ were less efficiently cut (Figure 3A). Surprisingly, the G-quadruplex forming sequence GGGGCC was the most efficiently cut, although it is supposed to form stable secondary structures *in vitro*

**Figure 3:**
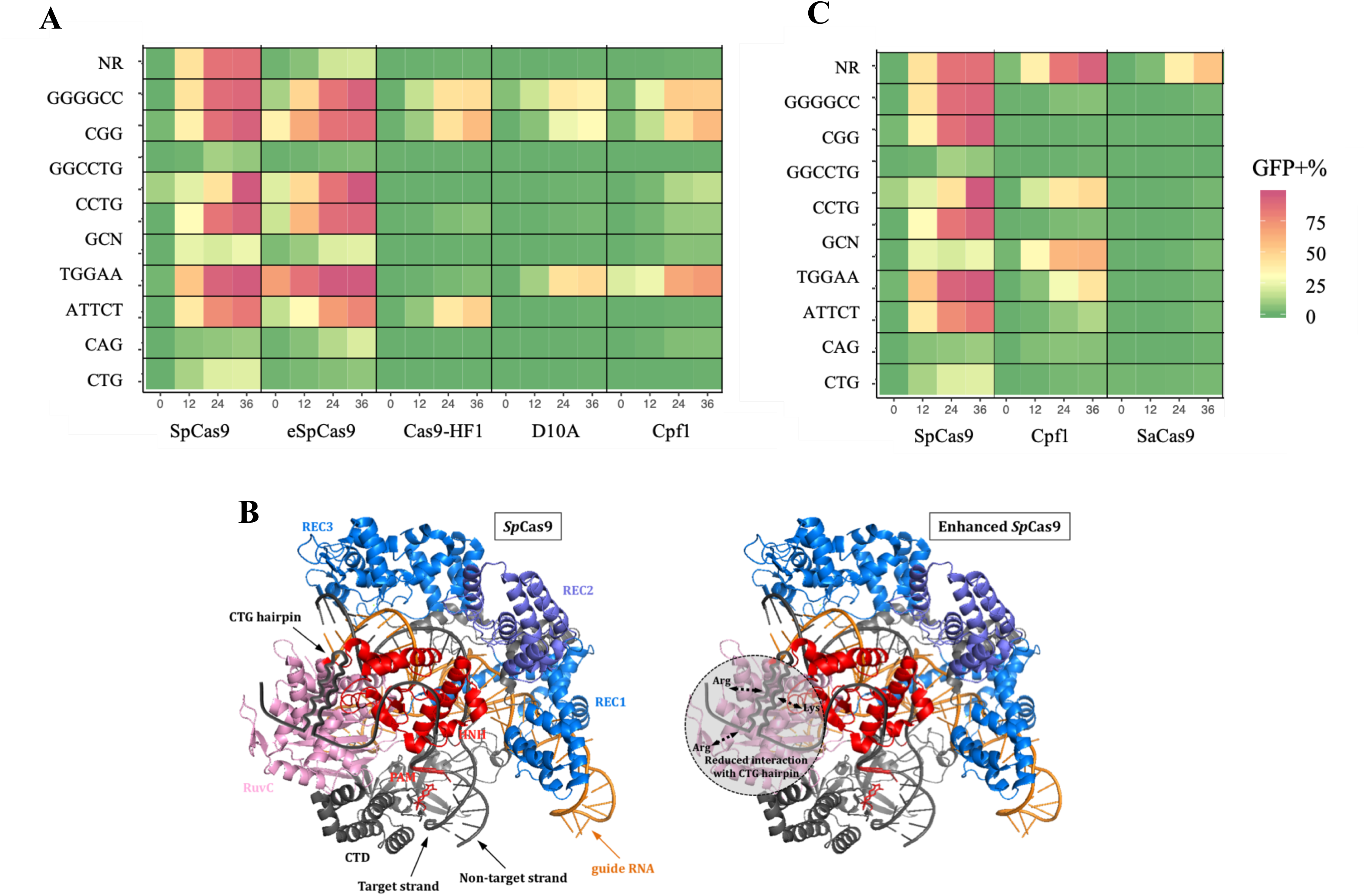
GFP-positive cells after DSB repair. **A:** SpCas9 and variants. NR: I-*Sce* I recognition site. Each microsatellite is shown on an horizontal line and called by its sequence motif. Recombination efficacies are indicated by the same color code as in Figure 2A. **B:** Reconstructed models of SpCas9 (left) and eSpCas9 (right) interacting with a structured CAG/CTG repeat, according to the SpCas9 crystal structure (PDB: 4UN3). In this model the CAG sequence is on the target strand whereas the CTG hairpin is on the non-target one. The three recognition domains are indicated in different shades of blue. The RuvC and the HNH nuclease domains are shown in pink and red, respectively. The three mutated amino acids in eSpCas9 (two arginine and one lysine residues) are also indicated. C: GFP-positive cells after SpCas9, SaCas9 or FnCpf1 inductions.

(Parkinson et al., 2002). This suggests that, despite possible secondary structures, this sequence is accessible to the nuclease *in vivo*. Alternatively, the presence of multiple PAMs at this locus may increase the chance that the nuclease would bind and make a DSB (Figure 1B). Two engineered variants of SpCas9 were then assayed. SpCas9 was more efficient that eSpCas9, itself consistently 2-10 times more efficient than Cas9-HF1 (Figure 3A). The NR sequence was also less efficiently cut, showing a general trend for these two variant nucleases. CTG repeats and CAG repeats were not cut the same way, eSpCas9 being more efficient on (CAG)_33_ than SpCas9, although the contrary was found for (CTG)_33_ (Figure 3A). It is known that CTG hairpins are more stable than CAG hairpins. Amrane *et al*., (2005) showed that the Tm of a (CTG)_25_ repeat was 58°C-61°C, depending on the method used for the measurement. In the same article, it was also shown that the Tm of a (CAG)_25_ repeat was only 54°C, proving that it was less stable. However, to the best of our knowledge, there is no evidence at the present time for formation of such secondary structures in living cells. However, given that we found opposite results with CAG and CTG repeat tracts, it is possible some kind of secondary structures may occur *in vivo*. Given that eSpCas9 shows a reduced interaction with the non-target strand, it may be inferred that a CTG hairpin on this strand should not affect eSpCas9 as much as its wild-type counterpart (Figure 3B). Therefore, CAG repeats on the target strand (CTG on the non-target strand) should be cut more efficiently by eSpCas9, as it was observed in the present experiments.

FnCpf1 was then tested on the same repeats (Figure 3C). (GAA)_33_ was the only one that was more efficiently cut by FnCpf1 than by SpCas9. This may be due to particular folding of the repeated sequence that makes it easier to cut by this nuclease. Alternatively, it may be due to the presence of several PAM on the complementary strand which may more easily attract this nuclease at this specific locus (Figure 1B).

None to very low level of recombinant cells were observed when SaCas9 was induced, although it efficiently cut the NR sequence (Figure 3C). This may be due to particular conformation issues of the DNA and/or of the guide or to their expression levels.

### gRNA and protein levels do not explain differences observed between nucleases

In order to determine whether DSB efficacies could be due to differences in protein levels or gRNA expression, we performed Western and Northern blots. For each guide, the signal corresponding to the expected RNA was quantified and compared to the signal of a control *SNR44* probe, corresponding to a snoRNA gene (Supplemental Figure S3A). For SpCas9, in one strain, (GCN)_33_, smaller species were detected, around 75 nt, that may correspond to degradation or abortive transcription. Using the classical phenol-glass beads protocol and despite numerous attempts, the FnCpf1 gRNA could not be detected. We hypothesized that it may be so tightly associated to its nuclease that phenol could not extract it, or that its amount was too low to be detected by Northern blot. Therefore, an alternative protocol used to extract very low levels of small RNAs was performed (see Materials & Methods), but did not allow to detect FnCpF1 gRNA. For SpCas9 and SaCas9, guide RNA levels were different among the ten strains. No correlation was found between gRNA quantification and GFP-positive cells, showing that gRNA steady state level was not the limiting factor in this reaction (Supplemental Figure S3B, left panel).

To assess the level of protein, total extracts were performed from yeast cells containing the NR sequence and the seven different nucleases. Note that different antibodies were used since proteins were not tagged. SpCas9 and its derivative mutant forms were detected with the same antibody, whereas SaCas9 and FnCpf1 were each detected with a specific monoclonal antibody (Supplemental Figure S3C). The same membranes were then stripped and rehybridized with an antibody directed against the product of *ZWF1*, encoding the ubiquitous glucose-6-phosphate dehydrogenase protein. Nuclease levels over control protein levels did not correlate with GFP-positive cells (Supplemental Figure S3B, right panel). Interestingly, the steady state level of eSpCas9 was found to be six times higher than SpCas9. This may be due to a higher stability of the protein, which could explain the high background of GFP-positive cells observed in repressed conditions (Supplemental Figure S2).

Overall, we concluded from these experiments that DSB efficacies were not obviously correlated to gRNA levels (at least for Sp- and SaCas9), nor to nuclease levels. This conclusion must be tempered by the fact that different antibodies with different affinities were used to detect nucleases. Therefore, we cannot totally rule out that SpCas9 was much more abundant than SaCas9 and/or FnCpf1 in our experiments.

### Secondary structure stability partly explains DSB efficacy

Trinucleotide repeats involved in human disorders are known to form stable secondary structures *in vitro*. This has been extensively studied and reviewed over the last 25 years (Gacy et al., 1995; Lenzmeier and Freudenreich, 2003; McMurray, 2010; Mirkin, 2006; Pearson et al., 2005; Richard et al., 2008; Usdin et al., 2015). Secondary structures are known to form both at DNA and at RNA levels (Kiliszek and Rypniewski, 2014; Kiliszek et al., 2010). It is however unclear if such structures actually exist in living cells, although genetic data strongly suggest that some kind of secondary DNA structures may be transiently encountered during replication and/or DNA repair. In our present experiments, secondary structures may possibly form on target DNA and on guide RNA. We therefore calculated theoretical Gibbs free energy for each target DNA and did not find any obvious correlation between structure stability and GFP-positive cells (Figure 4A, ANOVA test p-value=0.42) (see Materials & Methods). We subsequently performed the same calculation for the 20 nt guide RNA with or without their cognate scaffolds. Predicted structures of gRNA are shown in Figure 4B. When RNA scaffolds were taken into account, theoretical Gibbs energies were very low and comparable to each other, except for FnCpf1 guide RNA, which is much smaller than the others and for ATTCT gRNA that do not form secondary structure. This indicates that scaffold stability most frequently outweighs the 20 nt guide sequence stability. There was no correlation between scaffold stability and GFP-positive cells (Figure 4C, ANOVA test p-value=0.69). Finally, a statistically significant inverse correlation was found between the 20 nt guide RNA stability and GFP-positive cells (Figure 4D, ANOVA test p-value=0.015). We concluded that the 20 nt guide RNA stability was negatively correlated to GFP-positive cell formation, although it was not the sole determinant.

**Figure 4:**
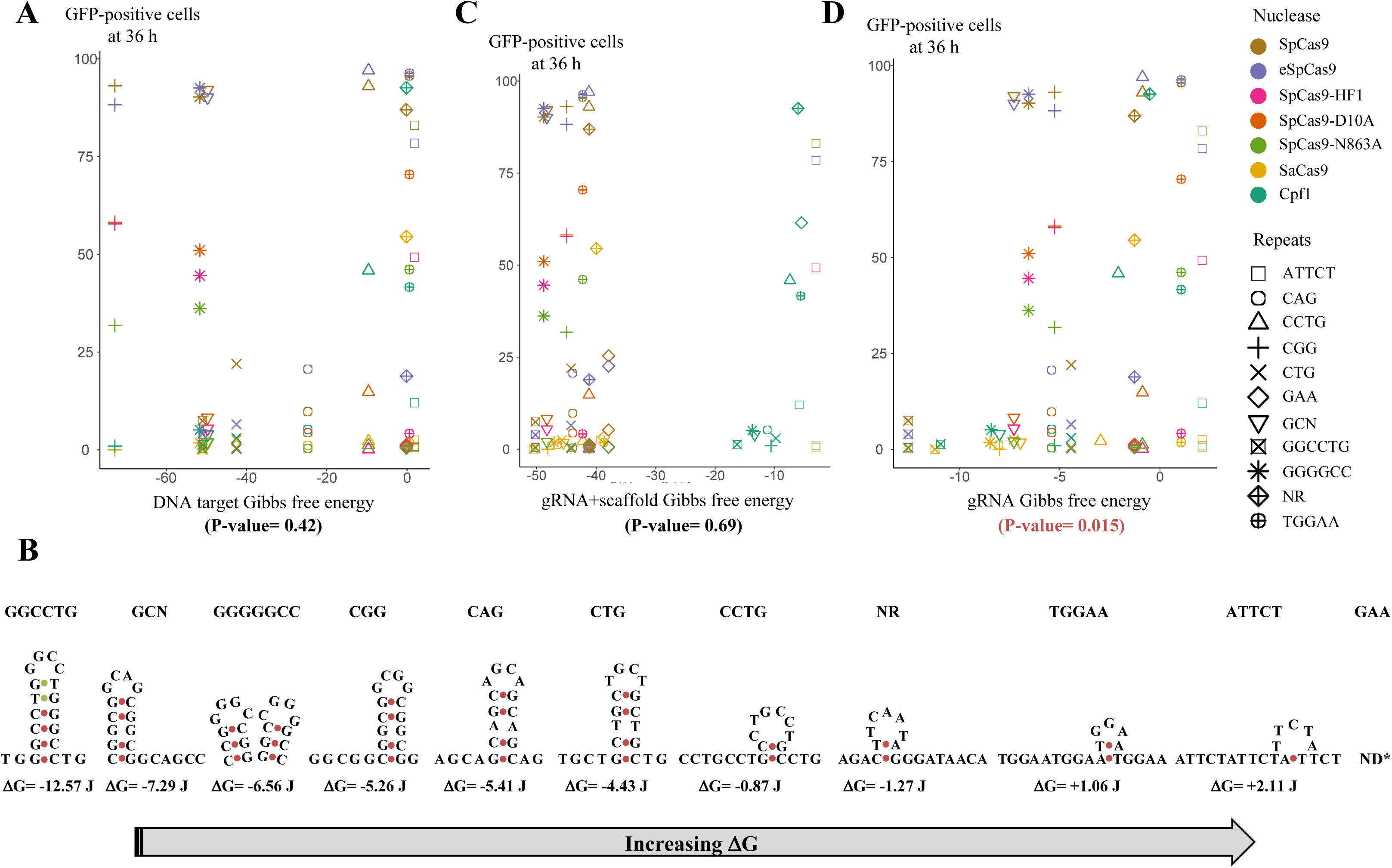
Secondary structures and Gibbs free energy. **A**: GFP-positive cells as a function of Gibbs energy calculated for each DNA sequence. Each nuclease is represented by a different color code and each microsatellite by a different shape. **B**: Predicted secondary structure of each SpCas9 gRNA. The *mfold* algorithm was used to model each structure. Only the most stable one is shown here. No structure could be calculated for GAA repeats. **C**: GFP-positive cells as a function of Gibbs energy calculated for each gRNA and its containing scaffold. **C**: GFP-positive cells as a function of Gibbs energy calculated for each gRNA alone. P-values are given below each graph.

### DSB-resection of microsatellites

Resection rate at *Sty* I restriction sites (Supplemental Figure S5) was measured by qPCR as previously described (Zierhut and Diffley, 2008) (Chen et al., 2013). Resected single-stranded DNA will not be digested by *Sty* I and will generate a PCR product whereas double-stranded DNA will be digested and will not be amplified. Resection ratios at 12h were calculated as resection at the repeat-containing end over resection at the non-repeated DSB end. They were normalized to the NR sequence whose ratio was set to 1. Resection values were only determined when the DSB was detected unambiguously at 12 hours. When SpCas9 was induced, resection rates were reduced at the repeated end as compared to the non-repeated end. When FnCpf1 was induced, resection rates were also lower on the repeated end for (GAA)_33_, (CTG)_33_ and (ATTCT)_20_, (CCTG)_25_ but not for (TGGAA)_20_. In conclusion, almost all repeats tested here inhibited resection to some level.

### Correlation between nuclease efficacy measured by flow cytometry and double strand break rate

To determine whether the flow cytometry assay recapitulates nuclease efficacy at molecular level, time courses were performed over 12-hour time periods for each nuclease-repeat couple. GFP-positive cell percentage at 12, 24 or 36 hours was plotted as a function of cumulative DSB over 12 hours (Figure 5). Given the number of different strains and nucleases tested, only one time course was performed in each condition (Supplemental Figure S5). However, data were very consistent between time points, showing that experimental variability was low. A linear correlation between the number of GFP-positive cells at 12 hours and the total signal of DSB accumulated during the same time period was found (linear regression test p-value=1.1×10^-9^, R^2^= 0.62) (Figure 5A). A good linear correlation was also found at later time points, 24 hours (p-value=2.6×10^-8^, R^2^=0.56) and 36 hours (p-value=4.8×10^-8^, R^2^=0.48) (Figures 5B, 5C). In conclusion, this GFP reporter assay is a good readout of double-strand break efficacy, and could be used in future experiments with other repeated sequences and different nucleases.

**Figure 5:**
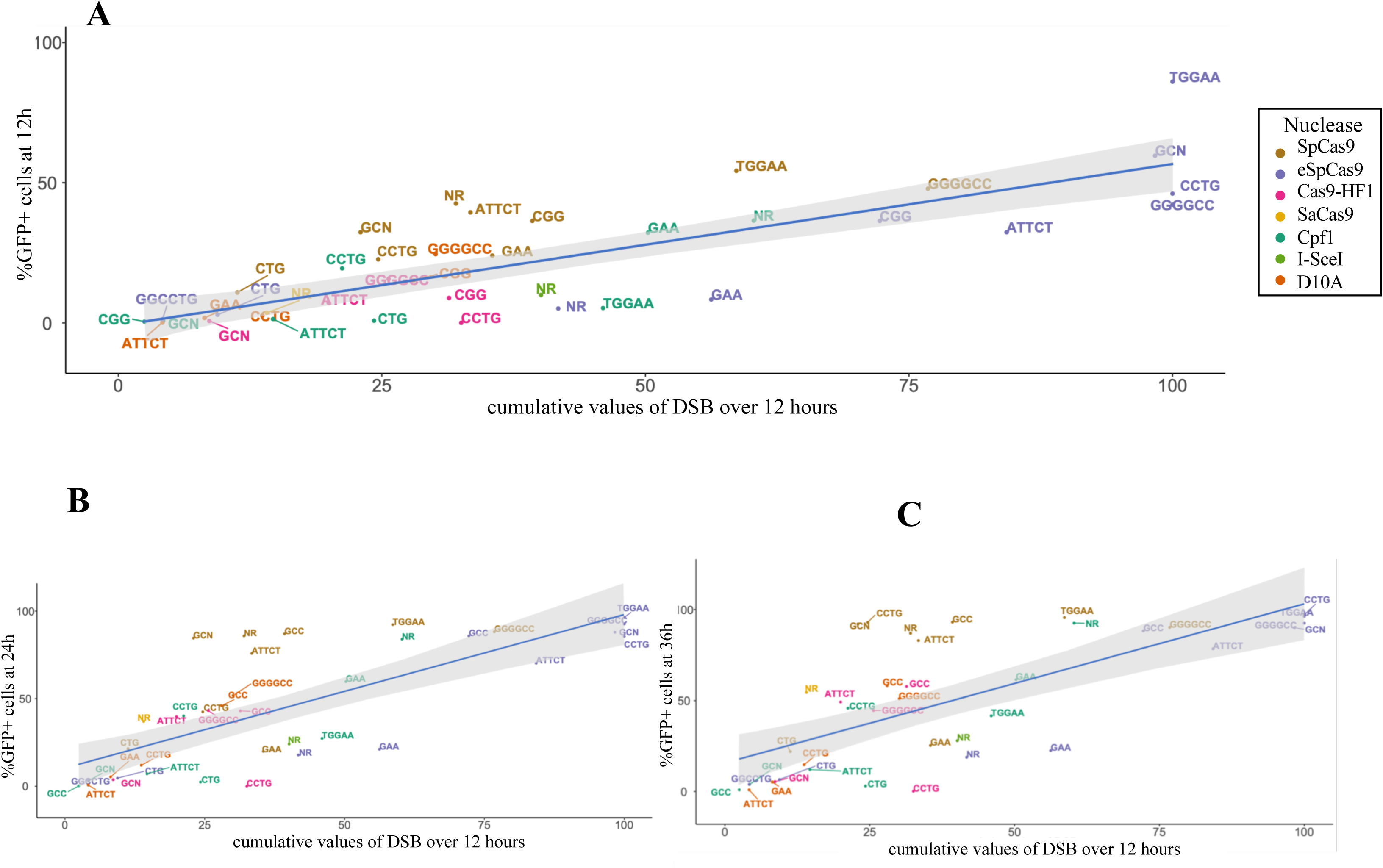
GFP-positive cell percentages as a function of DSB. Correlation between cumulative DSB level and GFP-positive cell percentage. Each color represents a nuclease. The blue line corresponds to the linear regression model. In grey: 95% confidence interval of the model. **A:** After 12 hours. **B**: After 24 hours. **C:** After 36 hours.

### Cas9-D10A nicks are converted to DSB *in vivo*

The Cas9 mutant D10A was more efficient than N863A on all repeats (Figure 3A). Surprisingly, both nickases were able to induce recombinogenic events on (GGGGCC)_15_, (GCC)_33_, (CCTG)_25_, (GCN)_33_, (GAA)_33_ and (TGGAA)_20_ repeats. SSBs are repaired by a specific machinery in yeast involving Base Excision Repair (BER) (Krokan and Bjørås, 2013). Nicks do not trigger homologous recombination unless they are converted to DSB. However, in our experiments, nicks trigger homologous recombination on some repeats. By Southern blot, DSBs were visible when Cas9-D10A was induced on those repeats (Supplemental Figure S5). These may be due to mechanical breakage during DNA preparation procedure, which converts SSB into DSB. Therefore, genomic DNA was prepared in agarose plugs to check this hypothesis. DNA extraction was carried out on Cas9-D10A time courses for (GAA)_33_ and (CGG)_33_ and NR sequences. DSBs were visible, suggesting that nicks were indeed converted into DSBs *in vivo* (Supplemental Figure S6).

### Genome-wide determination of off-target mutations

Microsatellites are very common elements of all eukaryotic genomes and the yeast genome contains 1,818 di-, tri- and tetranucleotide repeats (Malpertuy et al., 2003). In our present experiments, it was possible that other microsatellites of the yeast genome could also be mutated. We therefore decided to use an unbiased approach to determine all possible off-target sequences. The GUIDE-seq method was described in 2015 as a global approach to detect genome-wide DSB, and it was decided to adapt this method to budding yeast (see Materials & Methods). Shortly, cells were transformed with SpCas9 or FnCpf1, a gRNA and a modified double-stranded oligodeoxynucleotide (dsODN) to serve as a tag for targeted amplification. We chose the NR sequence as a control, as well as 6 out of 10 microsatellites cut by SpCas9 and the three most efficiently cut by FnCpf1. Colonies were collected, pooled and total genomic DNA extracted. Following random shearing and repair of DNA ends, two successive rounds of PCR were performed, using a primer complementary to the dsODN. DNA yield was unfortunately too low to be directly sequenced, and an additional round of PCR was performed (Figure 6A). The resulting libraries were loaded on an Illumina sequencer. Out of 7.8 millions reads, only 1.5 millions (1.9%) contained the dsODN. These reads were mapped to the yeast genome and found to be specially enriched at the rDNA locus and mitochondrial DNA (Supplemental Figure S7). In addition, from 7 – 2103 gene loci whose coverage was above twice the median coverage were identified in each library. These positions were compared to predicted off-targets using the CRISPOR web tool (Haeussler et al., 2016). Out of 68 genes in the CAG library, only one was predicted as a possible off-target and out of 2103 genes in the GAA library, only six were predicted as possible off-targets. In the libraries, there was an overlap between CRISPOR predictions and dsODN-containing gene loci. We therefore concluded that this approach was not efficient to identify real off-targets in the yeast genome.

**Figure 6:**
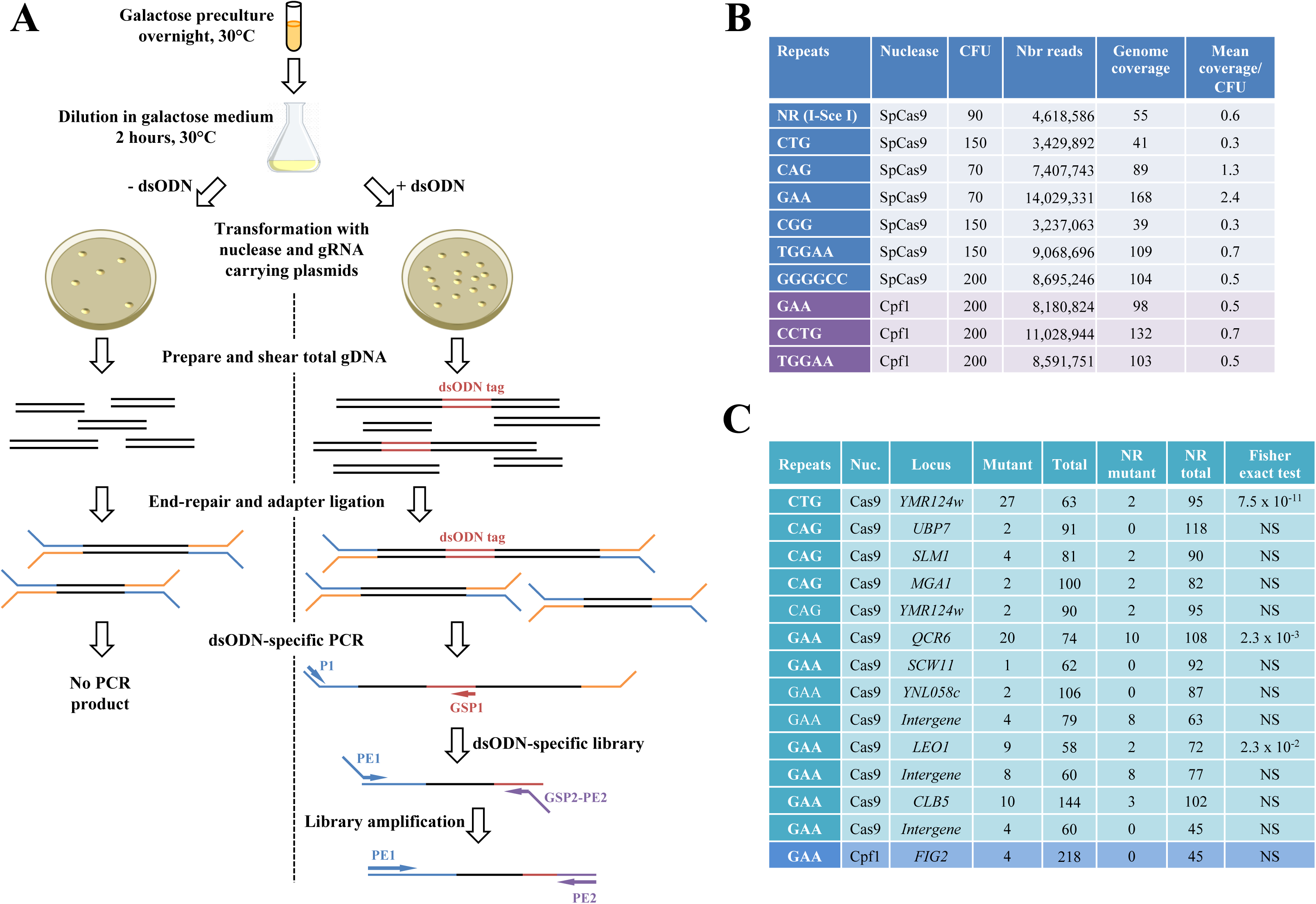
Off-target analysis. **A:** Cartoon depicting the experimental protocol (see text). **B**: Deep-sequencing results. For each library, the number of yeast colonies (CFU) after transformation and read numbers are given. Genome coverage was calculated by dividing (read number x 150 nucleotides) by 12.5 x 10^6^ nucleotides (haploid yeast genome). Mean coverage was found by dividing genome coverage by CFU. **C**: Statistical analysis. For each of the 14 putative off-targets, the Fisher exact test was used to compare mutants reads in each library to mutant reads in the NR library.

We decided to use a different approach to try to identify off-target sites, since that for each library, millions of reads homogeneously covered the whole genome. Classical SNP and indel calling algorithms aim at identifying frequent variants. However, off-targets are rare events, therefore the following pipeline of analysis was developed. In each library, variant reads were identified at each position predicted by CRISPOR. This ended up in 56 positions containing variant reads within microsatellites (Supplemental Table S5). Next, among these 56 positions, all positions containing only one mutant read were discarded. This left us with 14 genes containing at least one mutant read at a predicted off-target position. In order to determine whether these mutant reads were statistically significant, they were compared to the number of mutant reads at the same positions in the NR library used as a control. Given that colony number differed from one transformation to another one, the mean coverage per colony was used to normalize read number in each library (Mean coverage/CFU= genome coverage/CFU, Figure 6B). Once normalized, mutant reads in each library were compared to the NR control, using the Fisher exact test. Out of 14 possible off-target genes, only three exhibited read numbers significantly different from the NR control (Figure 6C). In the end, one gene (*YMR124w*) was identified to be a valid off-target for SpCas9 targeting CTG repeats, and two genes (*QCR6* and *LEO1*) were validated as off-targets for GAA repeats respectively targeted by SpCas9 and FnCpf1. Interestingly, all mutations in *YMR124w* were deletions of one or more triplets, but the two validated off-targets in the GAA library were all point mutations (Supplemental Figure S8). In conclusion, in the present experiments only nucleases targeted to CTG and GAA repeats exhibited some off-target effects.

## Discussion

Here, we successfully designed an assay for determining Cas9 variant efficacy on various microsatellites. Type II CRISPR-Cas nucleases were classified according to decreasing efficacies in the following order: SpCas9, eSpCas9, Cas9-HF1, SaCas9. FnCpf1, the only type V nuclease tested, was shown to exhibit substrate preferences different from type II nucleases. We also demonstrated that gRNA and protein levels did not generally correlate to nuclease activity and thus are not limiting factors in our experimental assay, ensuring that we are measuring nuclease activity and DSB repair *per se*.

### *In vivo* nuclease activities correlate to activities observed *in vitro*

Previous biophysical analyses showed that Cas9-HF1 and eSpCas9 bound to DNA similarly to SpCas9, but variants were trapped in an inactive state when bound to off-target sequences (Chen et al., 2017). Cas9-HF1 was more efficiently trapped in this inactive state than eSpCas9, showing more drastic impairment of cleavage. In our experiments, SpCas9 was more efficient than the two variants, confirming these biochemical data. Single molecule analyses enabled the precise determination of Cas9 binding and cleavage: first, the nuclease scrolls the genome for a PAM, then sequentially unwinds DNA starting from it (Sternberg et al., 2014). This explains why SpCas9 is not tolerant to mutations in the region proximal to the PAM. In our experiments, it may also explain why PAM-rich repeats were more easily cleaved, more protein could be recruited at the locus. However, Malina *et al* (2015) observed the opposite, decreased DSB repair when additional PAMs were present within the target sequence.

A very good correlation was generally observed between DSB efficacy and recombination (Figure 5). However, for some repeat/nuclease couples this was not the case (TGGAA/SpCas9 and GAA/eSpCas9 for example). We hypothesized that resection defects may lead to the observed phenotype, as we previously showed that a (CTG)_80_ repeat tract reduced resection efficacy in yeast in a *SAE2*-dependent manner (Mosbach et al., 2018). Comparison of resection values between repeated and non-repeated ends demonstrated that all repeats inhibit resection (Supplemental Figure S4). Therefore, differences in recombination are not due to resection defects alone. Note that in the present work, much shorter repeats (33 CTGs) were used as compared to our previous experiments with 80 CTGs.

### Nickases trigger homologous recombination on some repeat tracts

We confirm earlier findings that the RuvC Cas9-D10A mutant was more efficient than the HNH N863A variant (Gopalappa et al., 2018). Surprisingly, both nickases induced homologous recombination into GGGGCC, GCC, TGGAA repeat tracts and to a lower extent into CCTG, GCN and GAA repeat tracts (Supplemental Figure S2). Nicks are usually formed in the course of the BER pathway and trigger specific protein recruitment (Krokan and Bjørås, 2013). Nicks are therefore normally not processed by double-strand break repair machineries. However, there is some evidence supporting the hypothesis that nicks may be recombinogenic (Maizels and Davis, 2018; Strathern et al., 1991) which is in agreement with our data. For example, in *S. pombe,* mating type switching occurs by homologous recombination after the conversion of a nick into a DSB during replication (Arcangioli, 1998; Dalgaard and Klar, 2001). In our assay, replication may also convert a nick into a DSB, triggering homologous recombination in repeated sequences as suggested by the presence of a DSB observed throughout repair time course (Supplemental Figure S5). In a former work in human cells, Cas9-D10A was found to induce CTG/CAG repeat contractions, which may be due to the fact that many nicks were created into the target strand due to the repeated nature of the sequence, which then led to gap repair (Cinesi et al., 2016). In our experiment, gaps due to multiple nicks may arise in GGGGCC, CGG and TGGAA repeat tracts (Figure 1B), and GFP-positive cells were indeed observed in these three strains when Cas9-D10A was expressed. These multiple nicks may be either converted into DSB or form gaps that will then be converted into DSB.

### Correlation between secondary structure formation and nuclease efficacy

The sgRNA plays a crucial role in orchestrating conformational rearrangements of Cas9 (Wright et al., 2015). Stable secondary structure of the guide RNA as well as close state of the chromatin negatively affect Cas9 efficiency (Chari et al., 2015; Jensen et al., 2017). Possible secondary structures formed by the guide RNA are important to determine nuclease activity although there is no clear rule that can be sorted out and it is still challenging to know which hairpins will be detrimental (Thyme et al., 2016). We found that more stable gRNAs were correlated to less efficient DSBs (Figure 4D). This is consistent with former studies on non-repeated gRNAs showing that stable structured gRNAs (<-4 kcal/mol) were not efficient at inducing cleavage (Jensen et al., 2017). In addition, improperly folded inactive gRNAs could be competing with active and properly folded gRNAs within the same cell, to form inactive or poorly active complexes with Cas9, that will inefficiently induce a DSB (Thyme et al., 2016). Differential folding of CAG and CTG gRNA may explain the difference of efficacy observed with SpCas9 and eSpCas9. *In vitro* assays revealed that CTG hairpins were more stable than CAG hairpins because purines occupy more space than pyrimidines and are most likely to interfere with hairpin stacking forces (Amrane et al., 2005). This is probably also true *in vivo*, since CAG/CTG trinucleotide repeats are more unstable when the CTG triplets are located on the lagging strand template, supposedly more prone to form single-stranded secondary structures, than the leading strand template (Freudenreich et al., 1997; Viterbo et al., 2016). This difference in hairpin stability may impede Cas9/gRNA complex formation and/or impede recognition of target DNA by the complex. In our assay, SpCas9 cuts CTG more efficiently than CAG, whereas this is the other way around for eSpCas9. This may be due to reduced interaction between SpCas9 and the non-target strand, that is probably more prone to form secondary structures (Figure 3B).

Finally, a lower preference for T and a higher preference for G next to the PAM was previously reported (Chari et al., 2015). Other nucleotide preferences were found (Doench et al., 2014) but the preference for a G at position 20 of the guide is consistent across studies. This may explain why SpCas9 may be more efficient on CGG, CCTG and less on ATTCT repeats (Figure 3A).

### Defining the best nuclease to be used in gene therapy

It was previously shown that a DSB made into CTG repeat tracts by a TALEN was very efficient to trigger its shortening (Mosbach et al., 2018; Richard, 2015). Other approaches may be envisioned to specifically target toxic repeats in human using the CRISPR toolkit: i) Cas9-D10A induced CAG contractions (Cinesi et al., 2016), ii) dCas9 targeting microsatellites was able to partially block transcription, reversing partly phenotype in DM1, DM2 and ALS cell models (Pinto et al., 2017), iii) efficient elimination of microsatellite-containing toxic RNA using RNA-targeting Cas9 was also reported (Batra et al., 2017). Finally, if using CRISPR endonucleases to shorten toxic repeats involved in microsatellite disorders was envisioned, our study will help finding the best nuclease. For example, Fragile X syndrome CGG repeats could be efficiently targeted with SpCas9. It must be noted that all human microsatellites may not be targeted by all nucleases tested here, for some of them lacking a required PAM. However, our results allow to discard inefficient nucleases for further human studies.

However, specificity must also be taken into consideration. Previous analyses showed that the yeast genome contained 88 CAG/CTG, 133 GAA/CTT and no CGG/CCG trinucleotide repeats (Malpertuy et al., 2003). The dsODN tag was preferentially found at the rDNA locus and in mitochondrial DNA. This suggests that random breakage occurs frequently within these repeated sequences. This is compatible with the high recombination rate observed at the rDNA locus following replication stalling (Mirkin and Mirkin, 2007; Rothstein et al., 2000). However, using an alternative method to detect rare variants we were able to identify three real off-targets in the yeast genome. Off-target mutations were found in one CTG repeat out of 88 and two GAA repeats out of 133, for SpCas9 and FnCpf1. This shows that although very frequent sequences like microsatellites were predicted to be off-targets, few real mutations were indeed retrieved. By comparison, the human genome contains 900 or 1356 CAG/CTG repeats, depending on authors (Kozlowski et al., 2010)(Lander et al., 2001). Given our results, we can predict that ca. 1% of these would be real off-targets for a SpCas9 directed to a specific CTG microsatellite. However, in our experiments, the nuclease was continuously expressed, which is not envisioned in human genome editing approaches. Reducing the expression period of the nuclease should also help reducing off-target mutations, but this has now to be thoughtfully investigated.

The GUIDE-seq method was very successful at identifying off-target site sin the human genome, following Cas9 expression. In *S. cerevisiae*, we showed here that this approach was not efficient, most probably because NHEJ is not as active as in human cells, particularly in haploid yeast in which it is downregulated (Frank-Vaillant and Marcand, 2001; Valencia et al., 2001). Altogether, our results give a new insight into which nuclease could be efficiently used to induce a DSB into a microsatellite in other eukaryotes.

## Acknowlegments

L. P. was supported by a CIFRE PhD fellowship from Sanofi. Off-target studies were supported by the AFM-Telethon. We thank Heloïse Muller for sharing her unpublished protocol for yeast transformation by electroporation, and Carine Giovannangeli for the generous gift of CRISPR-Cas plasmids. This work was supported by Sanofi, the Institut Pasteur and the Centre National de la Recherche Scientifique (CNRS).

## Supplemental Figures

**Supplemental Figure S1: Flow cytometry at 36 hours for each repeat.** For each repeat (horizontal) the corresponding dot plot is shown for each nuclease (vertical). Gates are drawn to separate recombined from non-recombined populations.

**Supplemental Figure S2: Histogram of GFP-positive cell percentages for each repeat and each nuclease.** Galactose condition is highlighted in green and glucose condition in black for each of the four time points. Each experiment was performed 3-8 times, depending on the strain. Error bars are standard errors.

**Supplemental Figure S3: Southern blots and quantifications.** For each repeat, a Southern blot of an induction time course over 12 hours is shown above quantification graphs. Recombination and DSB percentages were calculated as fractions of the total signal in each lane.

**Supplemental Figure S4: gRNA and protein levels. A:** gRNA expression levels measured by Northern blot. Top: Northern blot was hybridized with a SpCas9 gRNA scaffold probe. Bottom: Same for SaCas9. The same Northern blots were rehybridized with a control *SNR44* probe, corresponding to a snoRNA gene. **B**: Signals of both gRNA and *SNR44* were quantified and their ratios compared to nuclease efficacy measured by the percentage of GFP-positive cells at 36 hours. **C:** Nuclease expression levels measured by Western blot. Blots were successively hybridized with Cas-specific antibodies and Zwf1p antibody. Ratios of Cas/Zwf1 signals are shown below the blots. Left: GFP+ cell percentage as a function of gRNA levels. Right: GFP+ cell percentage as a function of protein levels.

**Supplemental Figure S5: Evidence for *in vivo* DSBs generated by nickases.** Top: Southern blot of DNA prepared in agarose plugs for three strains (Non-repeated, CGG and GAA repeats) after 4, 8 and 12 hours of nuclease induction. Parental, recombinant and DSB bands are indicated. DSBs are visible for CGG and GAA strains, increasing with time. The asterisk indicates a migration artefact commonly observed when overloading DNA in a lane. Bottom: The same experiment was performed but DNA was prepared by the standard protocol. The DSB was faintly detected at similar time points. Note that for the control non-repeated strain, some DSB signal is detected at the T0 time point only, suggesting that it could correspond to mechanically broken DNA molecules.

**Supplemental Figure S6: Resection analysis of SpCas9 and FnCpf1 induced double-strand breaks.** Sty I restriction sites located on each side of the DSB site are indicated. Only yeast strains for which a DSB was detectable at 12 hours post-nuclease induction were analyzed. Resection at NR was set at 100% to normalize the data. Lower resection values indicate resection inhibition.

**Supplemental Figure S7: dsODN genome coverage.** Each horizontal line represents a whole genome, in which each chromosome is separated by a dashed line and identified by a roman number. Red arrows point to regions enriched for the dsODN tag (rDNA and mitochondrial DNA).

**Supplemental Figure S8:** Mutations identified in each of the three off-target sequences. Deletions are indicated by a red Δ, insertion in blue, and base substitutions in green. Triplets different from the microsatellite consensus are grey.

