## Supplementary figures and images for "Differential efficacies of Cas nucleases on microsatellites involved in human disorders and associated off-target mutations"

### Sup. Fig 1

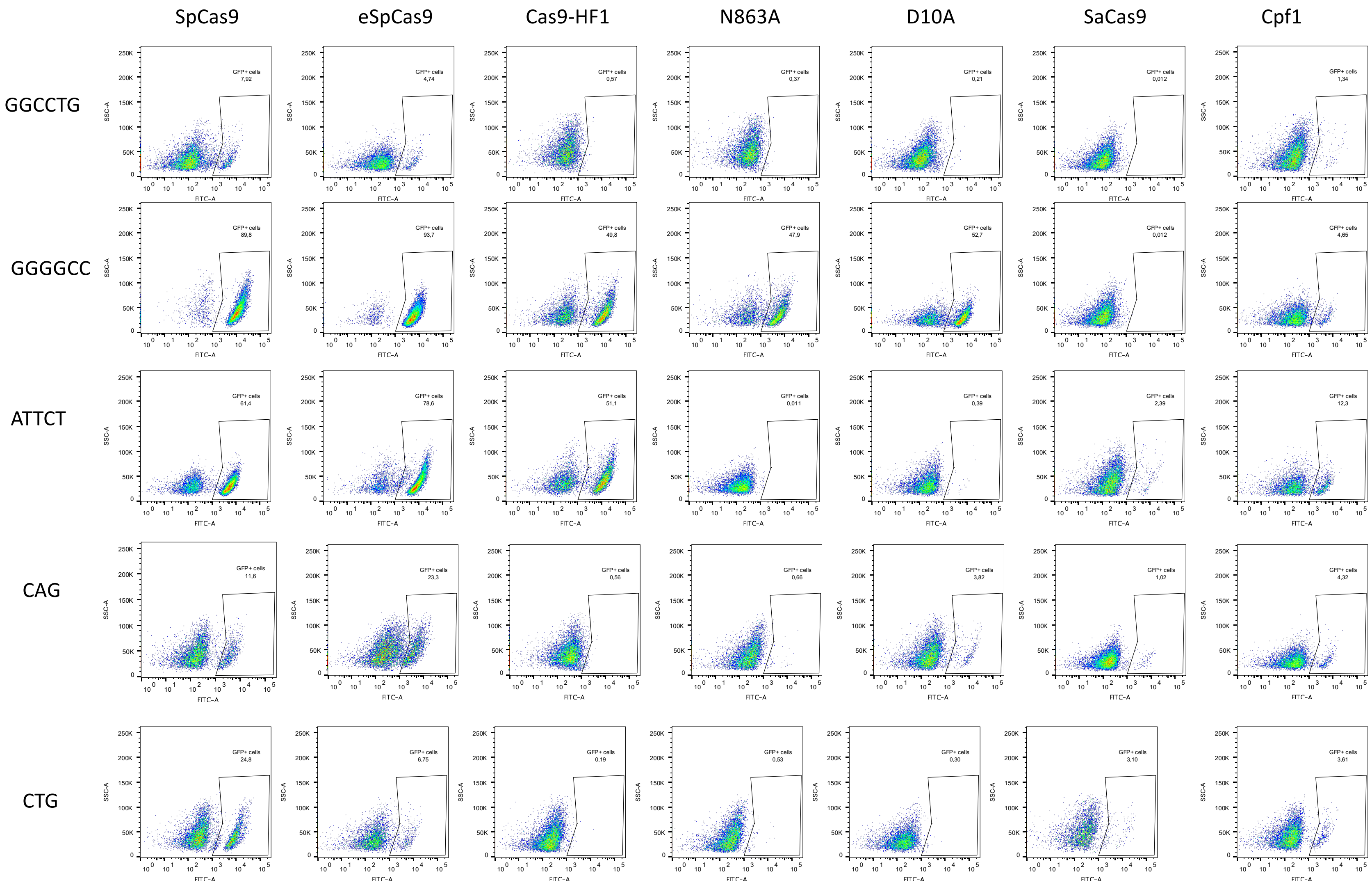


**Poggi *et al.***
**Figure S1**

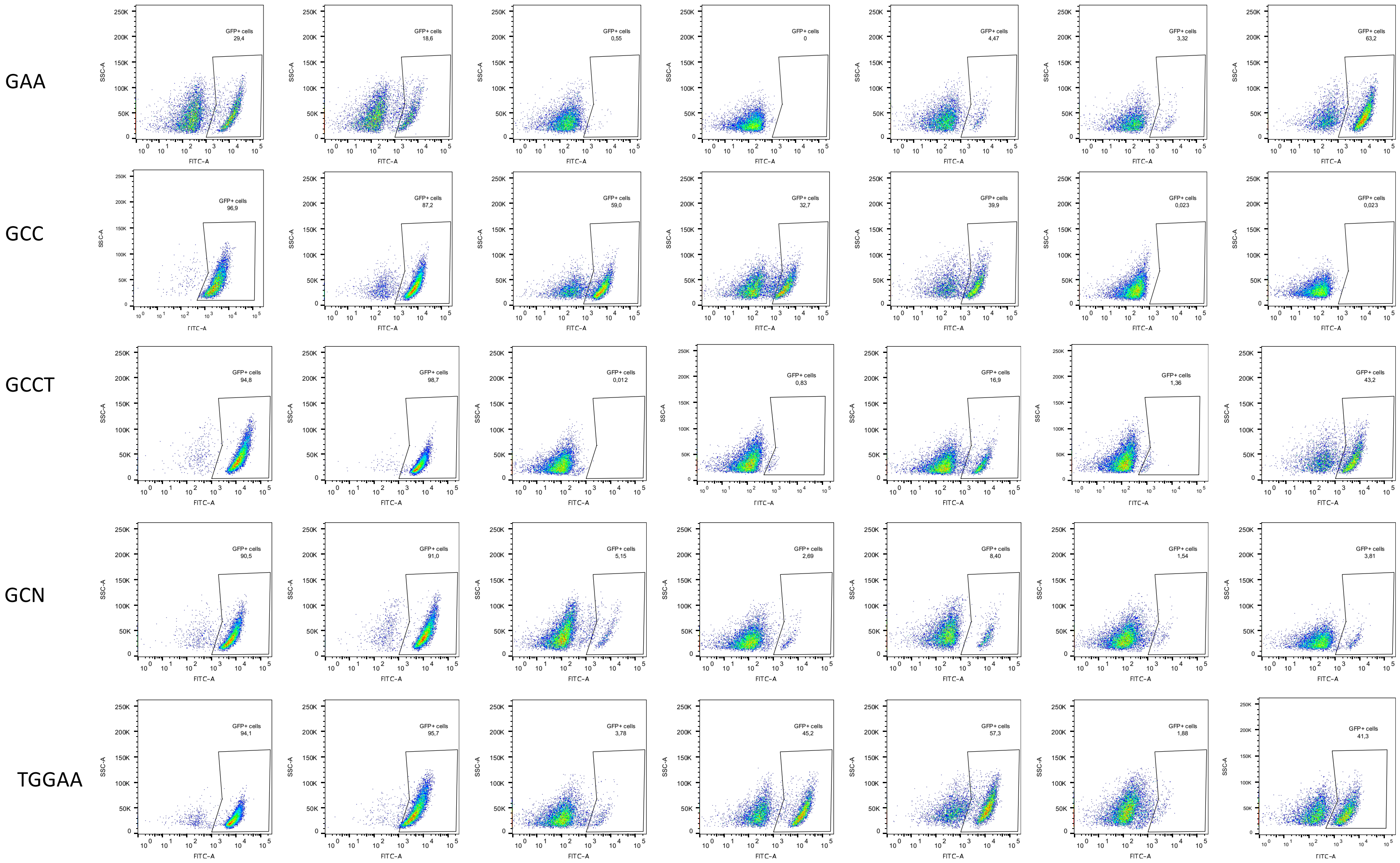


**Poggi *et al.***
**Figure S1**

Not repeated

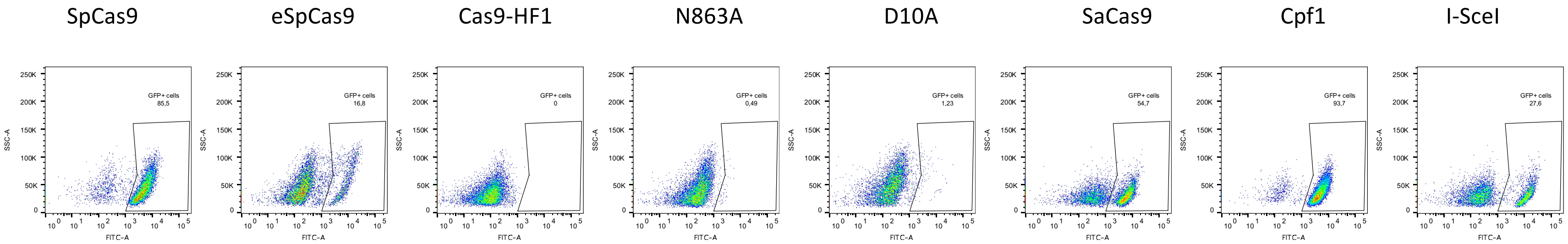


**Poggi *et al.***
**Figure S1**

### Sup. Fig. 2

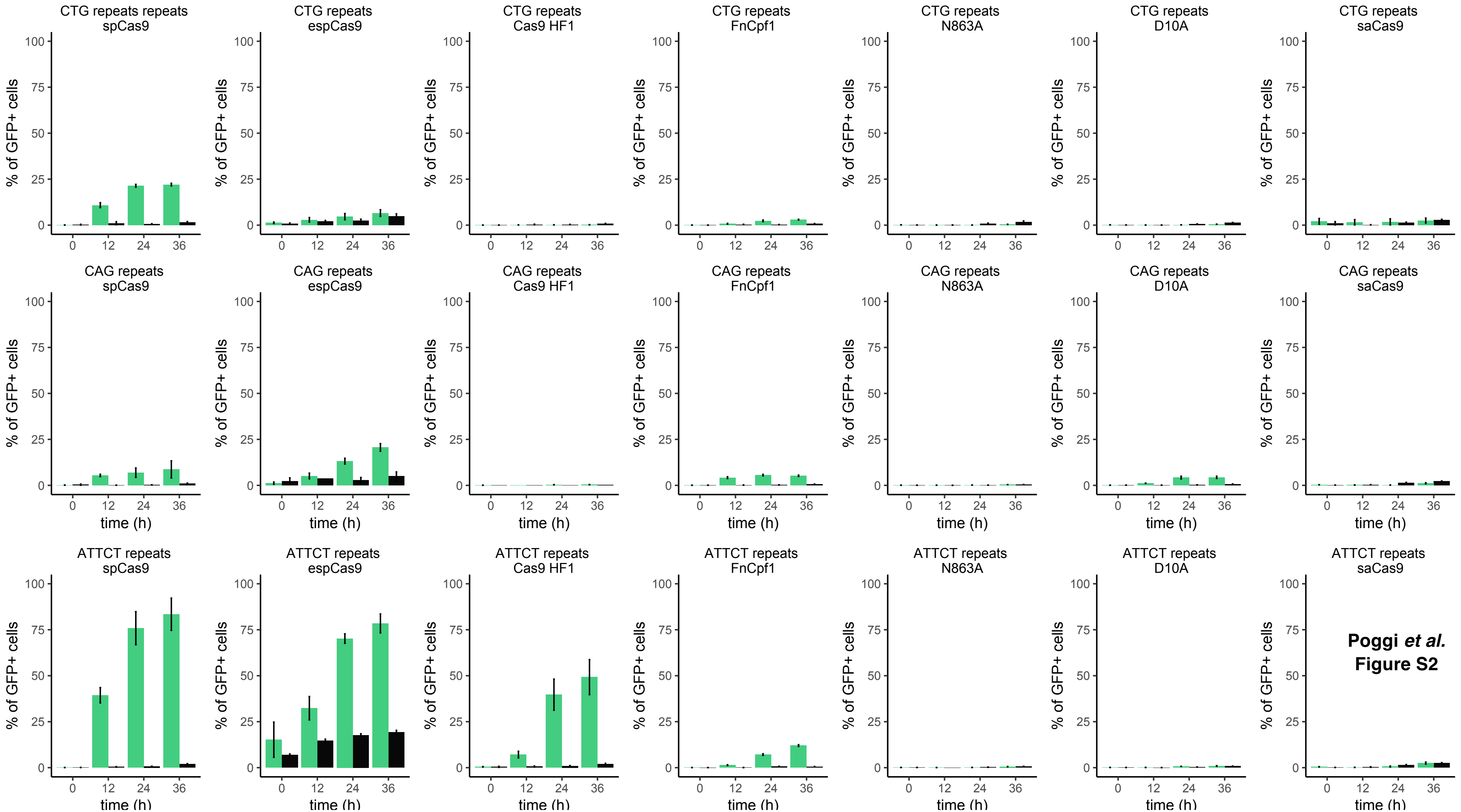


**Poggi *et al.***
**Figure S2**

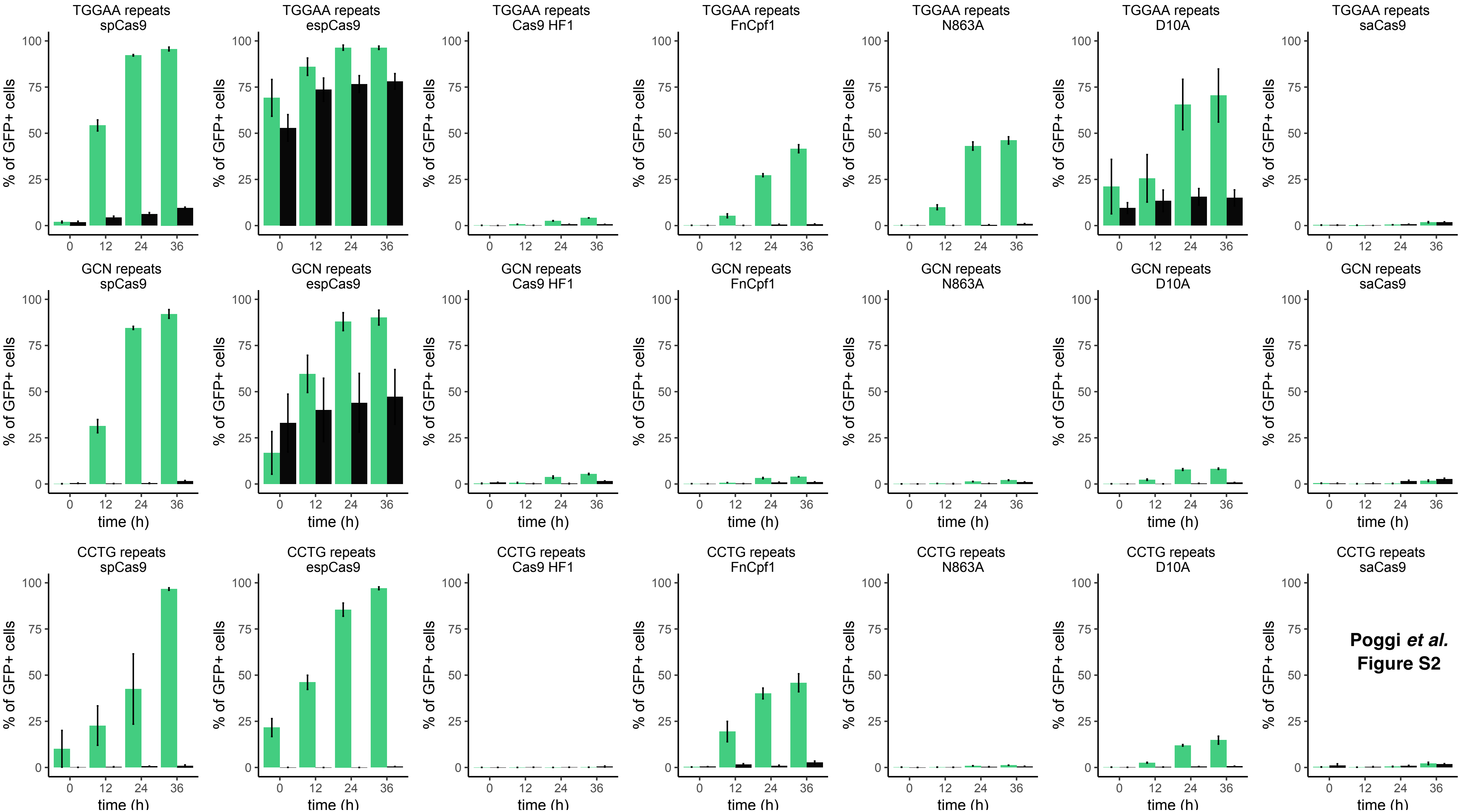

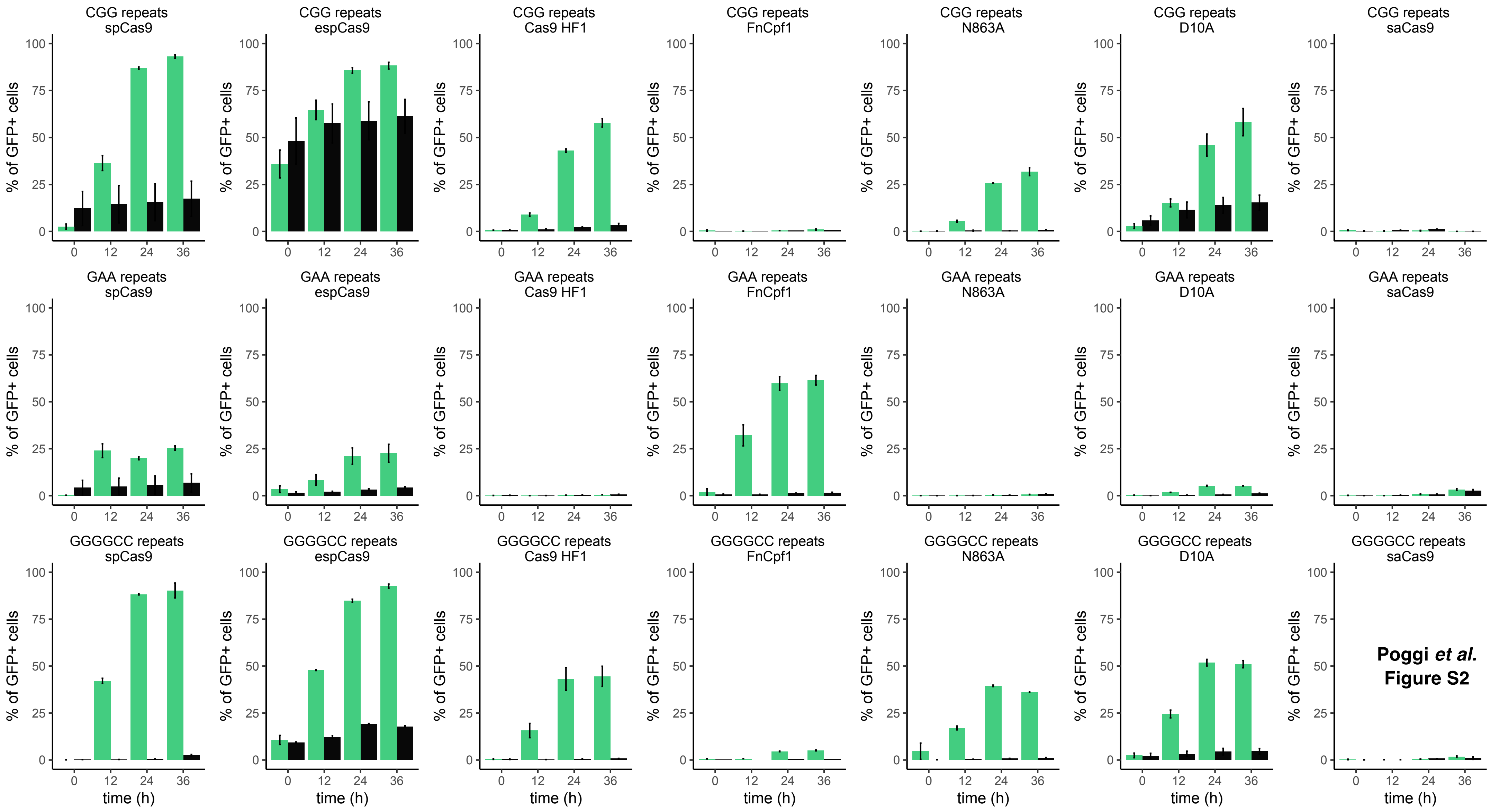


**Poggi *et al.***
**Figure S2**

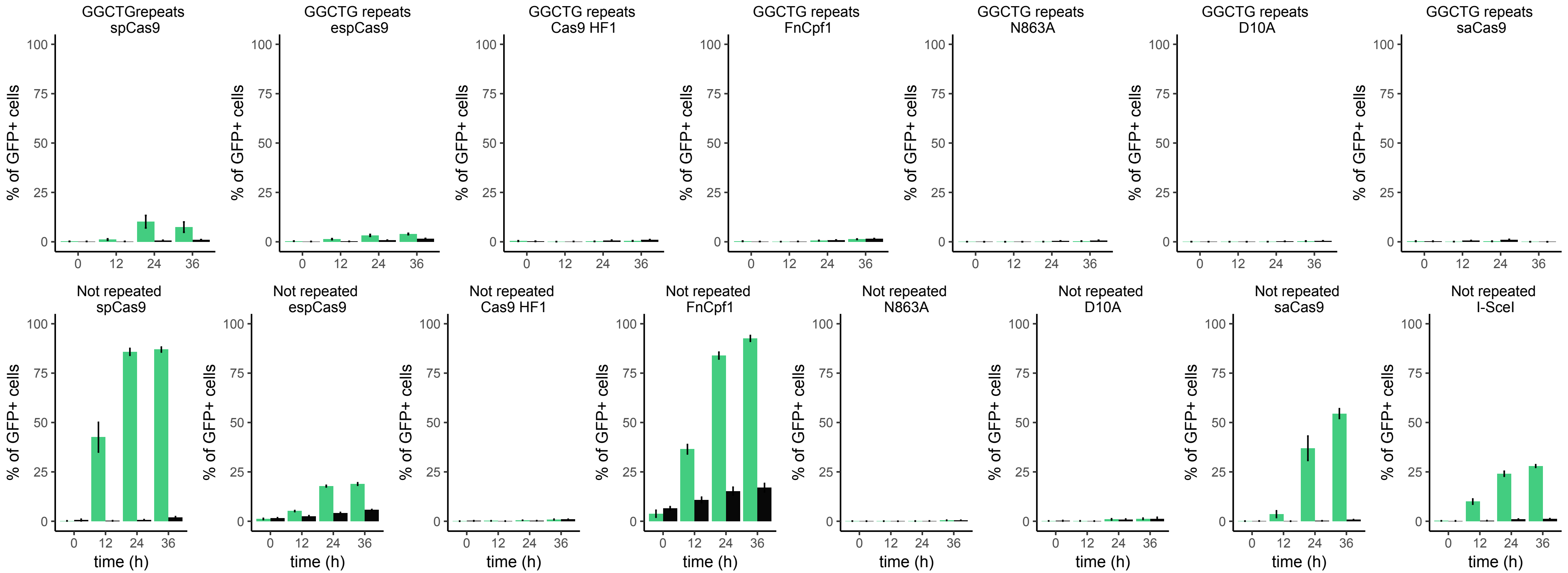


**Poggi *et al.***
**Figure S2**

### Sup. Fig. 3

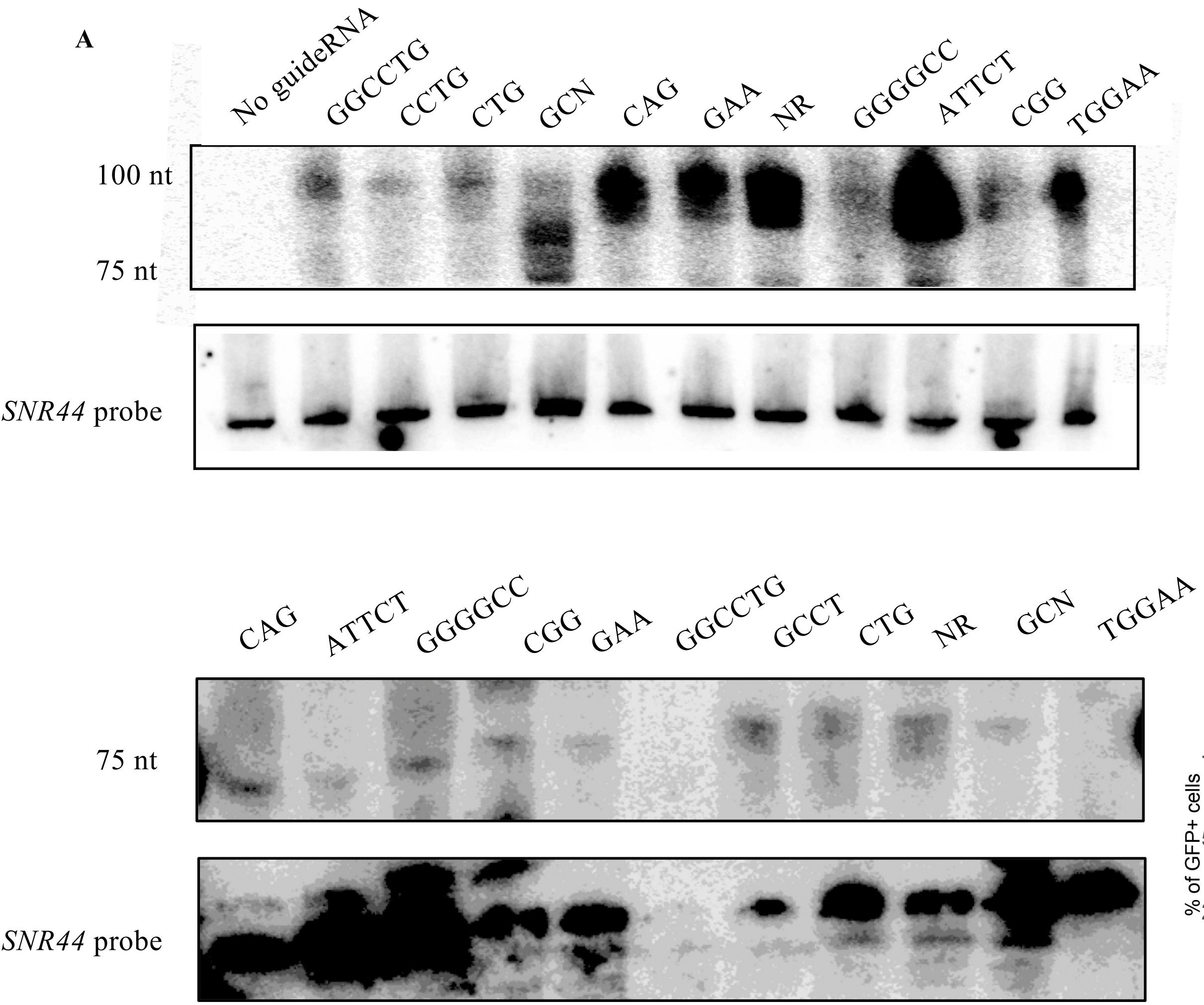


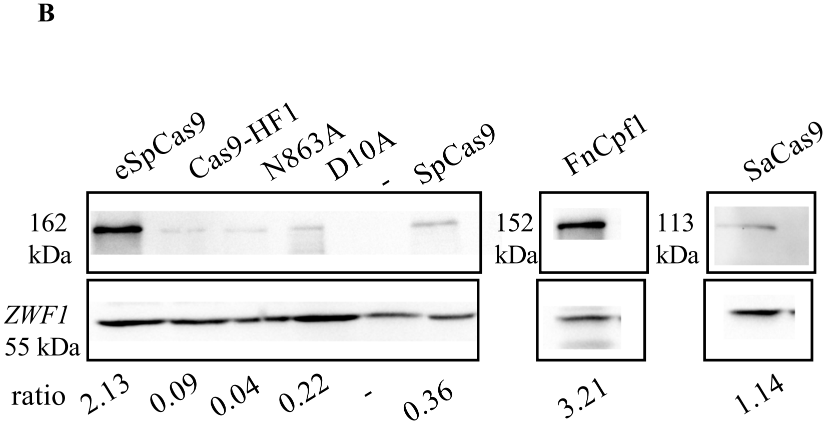


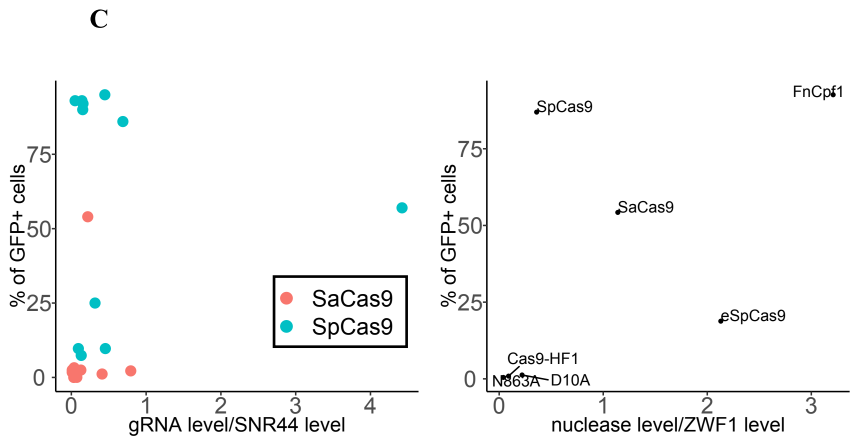


**Poggi *et al.* Supplemental Figure S3**

### Sup. Fig. 4

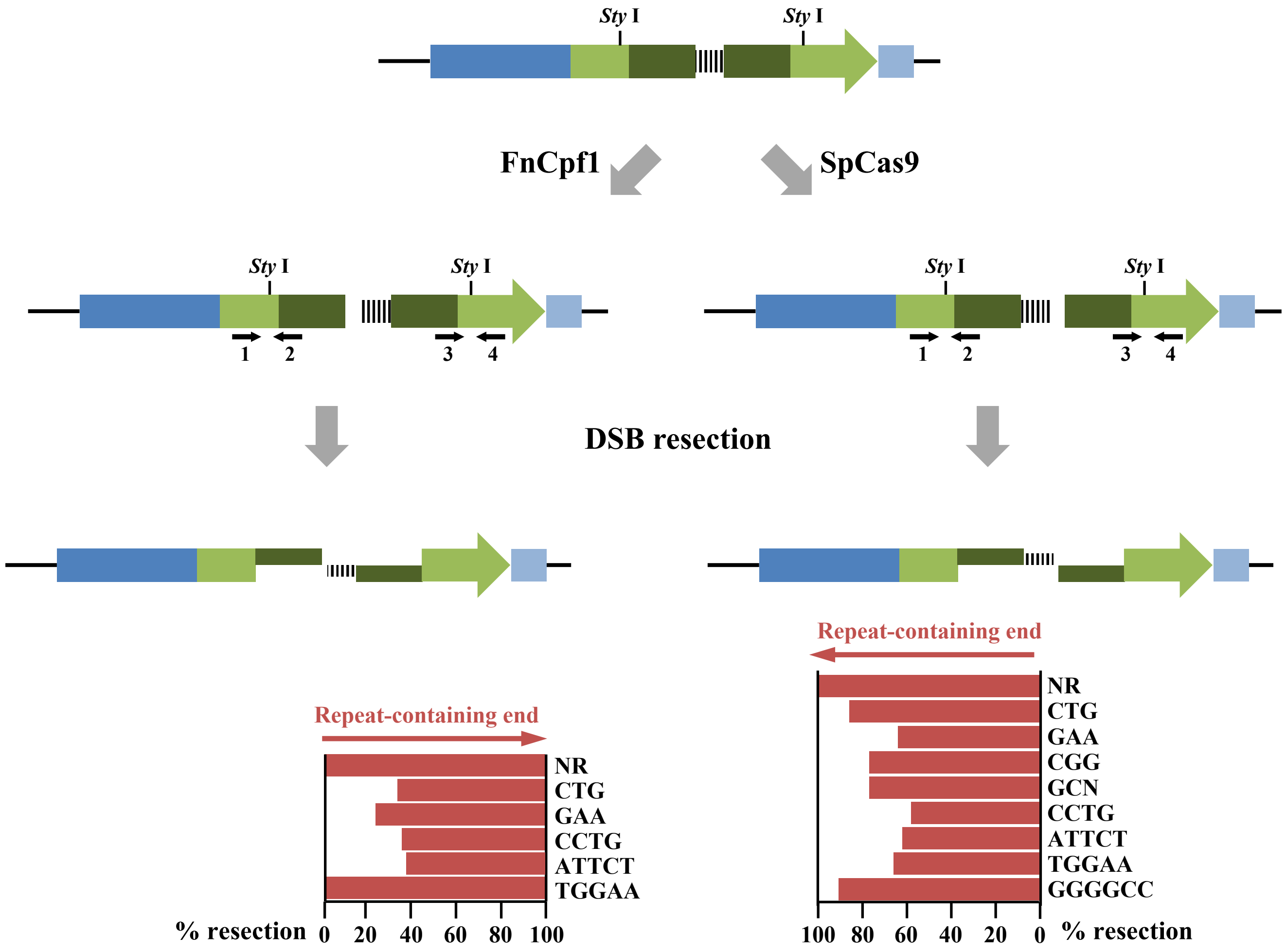


Poggi et al.
Supplemental Figure S4

### Sup. Fig. 6

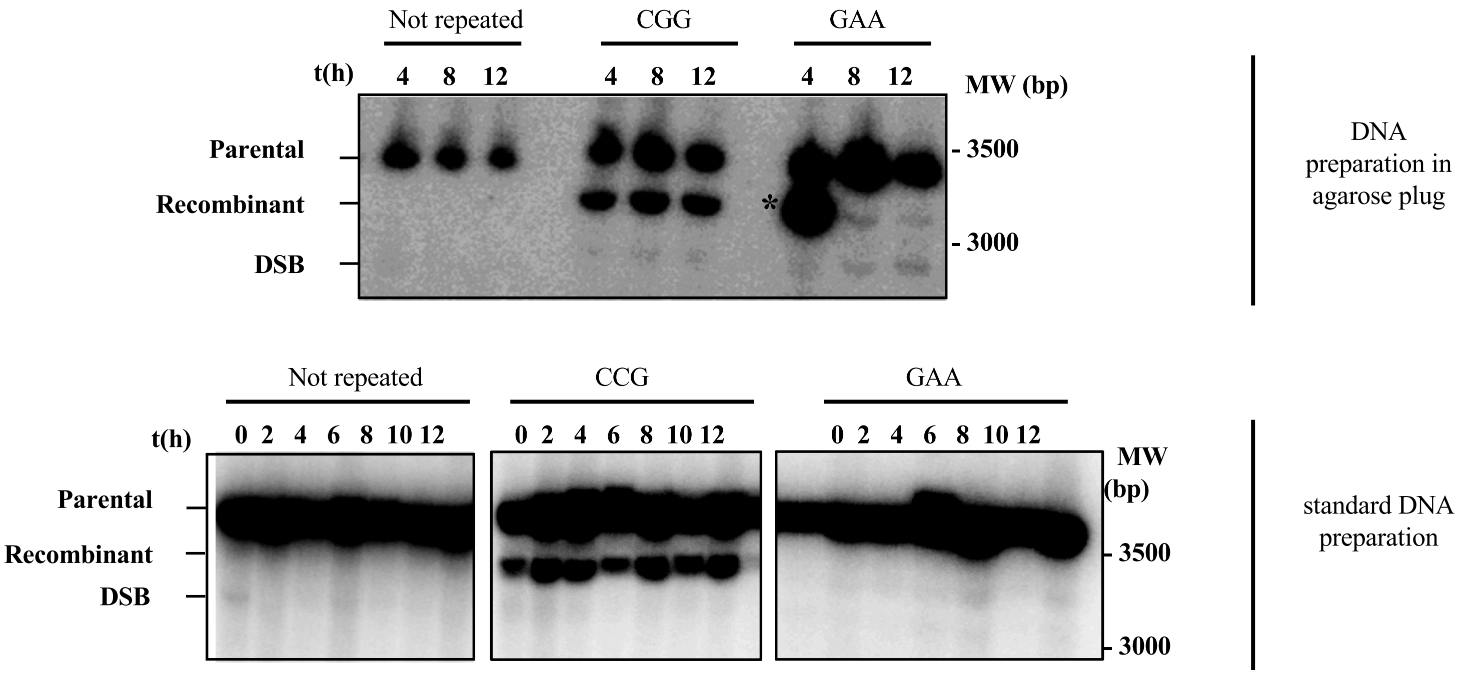


**Poggi *et al.***
**Supplemental Figure S6**

### Sup. Fig. 7

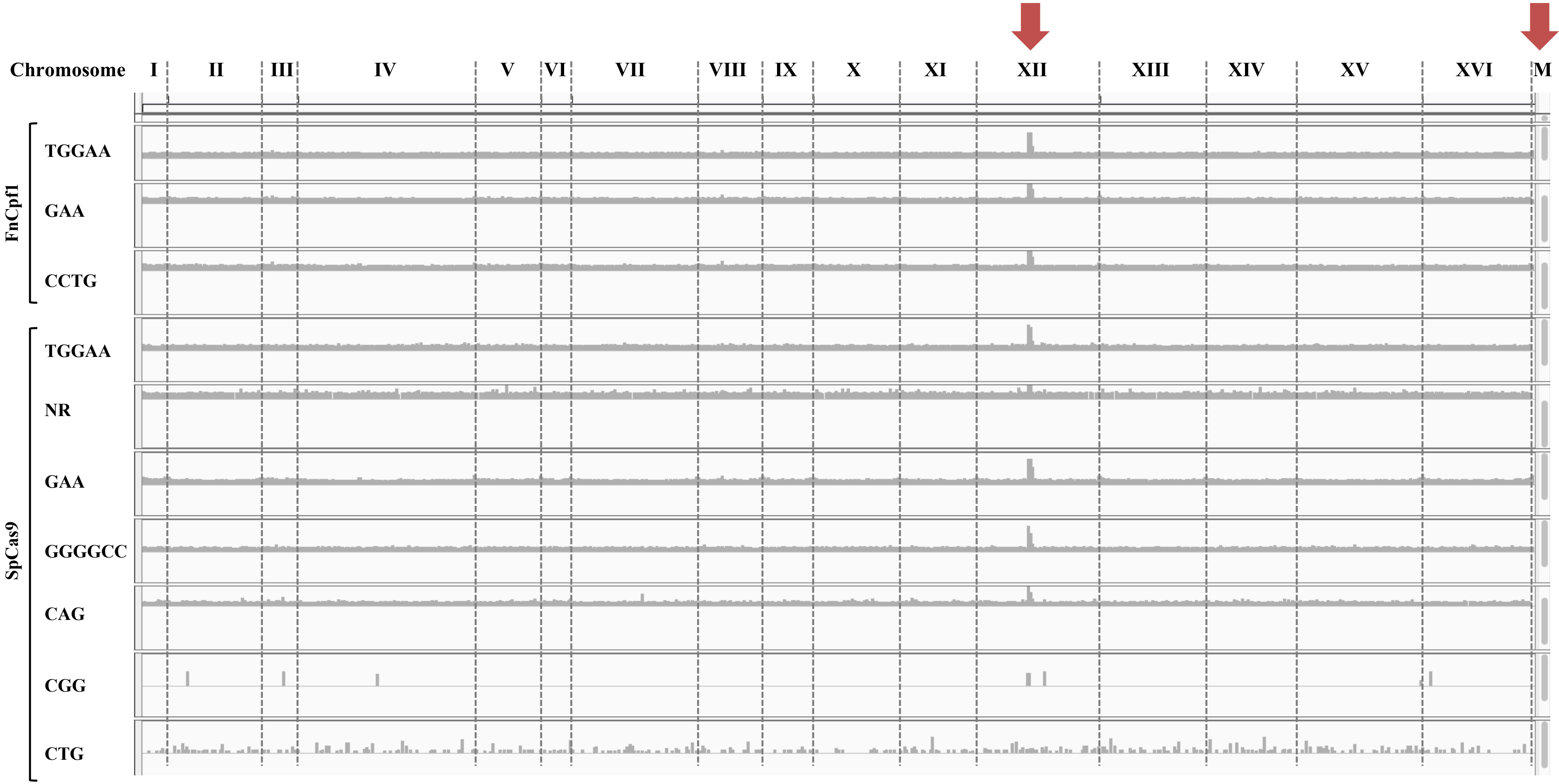


Poggi et al.
Supplemental Figure S7

### Supplemental Data 1

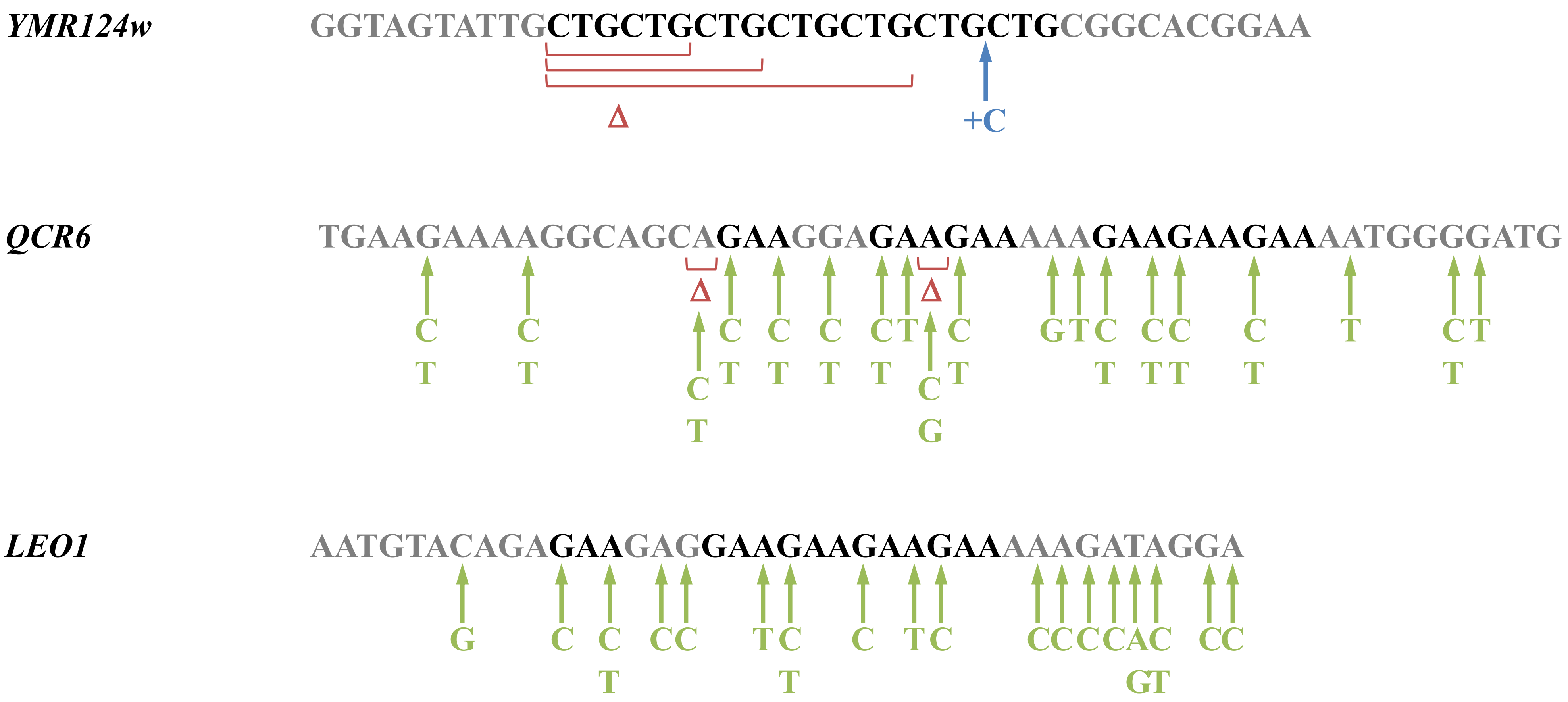


Poggi et al.
Supplemental Figure S8
