## Supplementary material for "Differential efficacies of Cas nucleases on microsatellites involved in human disorders and associated off-target mutations": Sup. Fig. 5

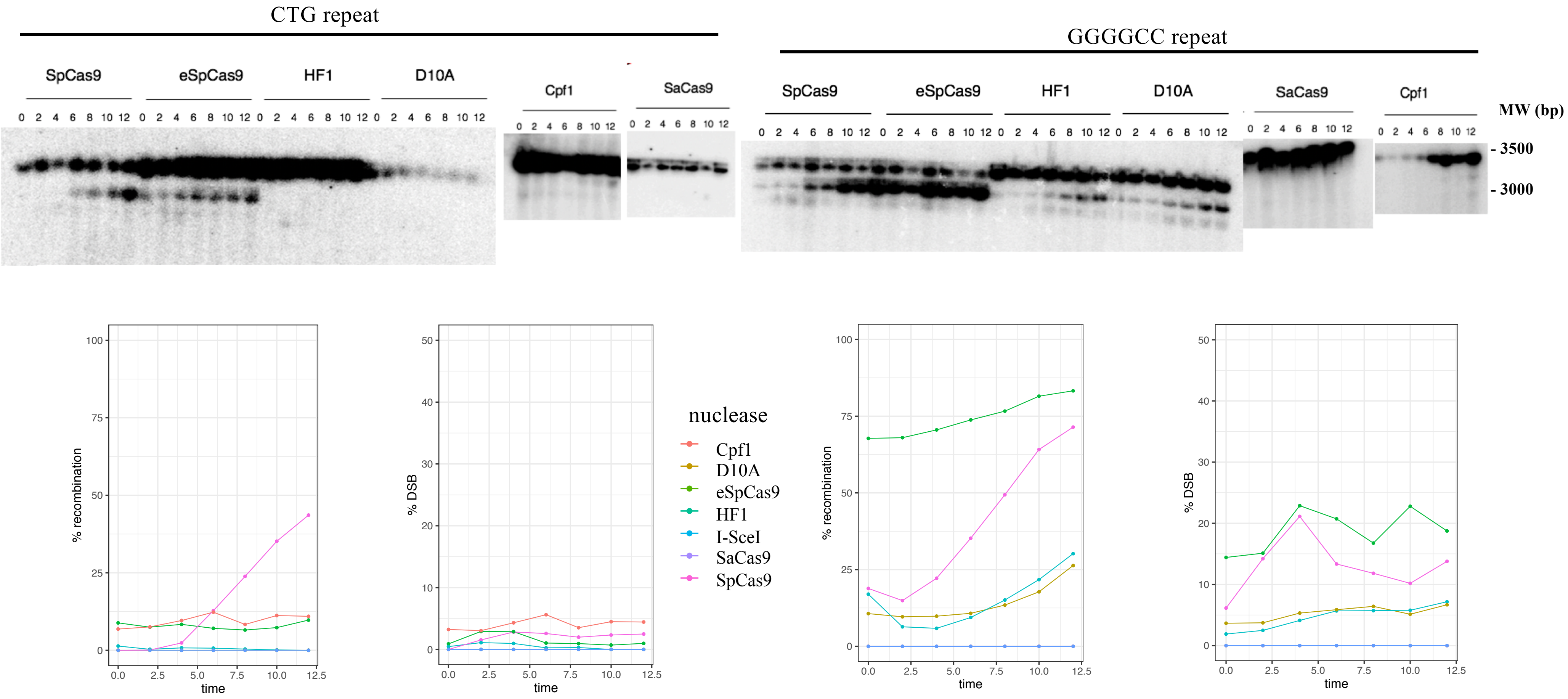


**Poggi *et al.* Supplemental Figure S5**

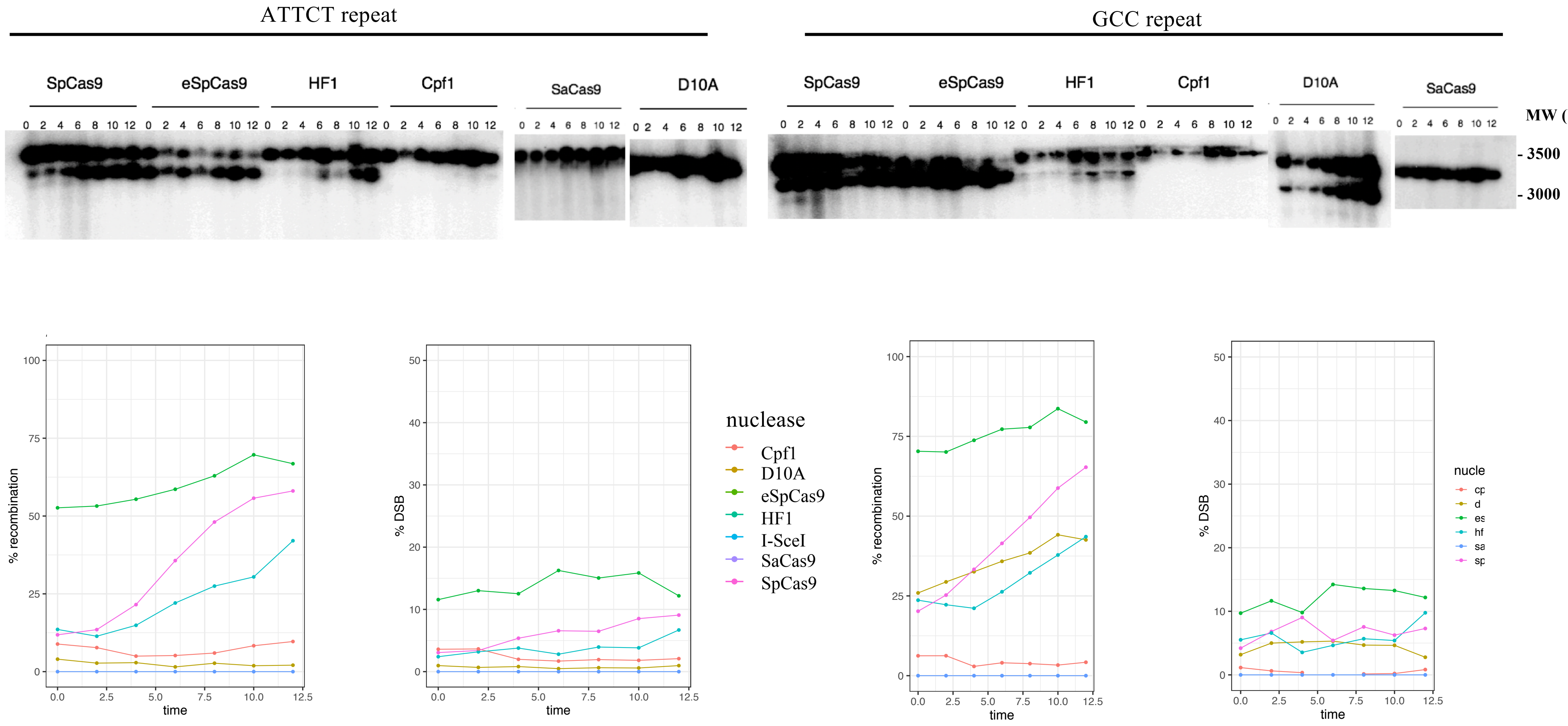


**Poggi *et al.* Supplemental Figure S5**

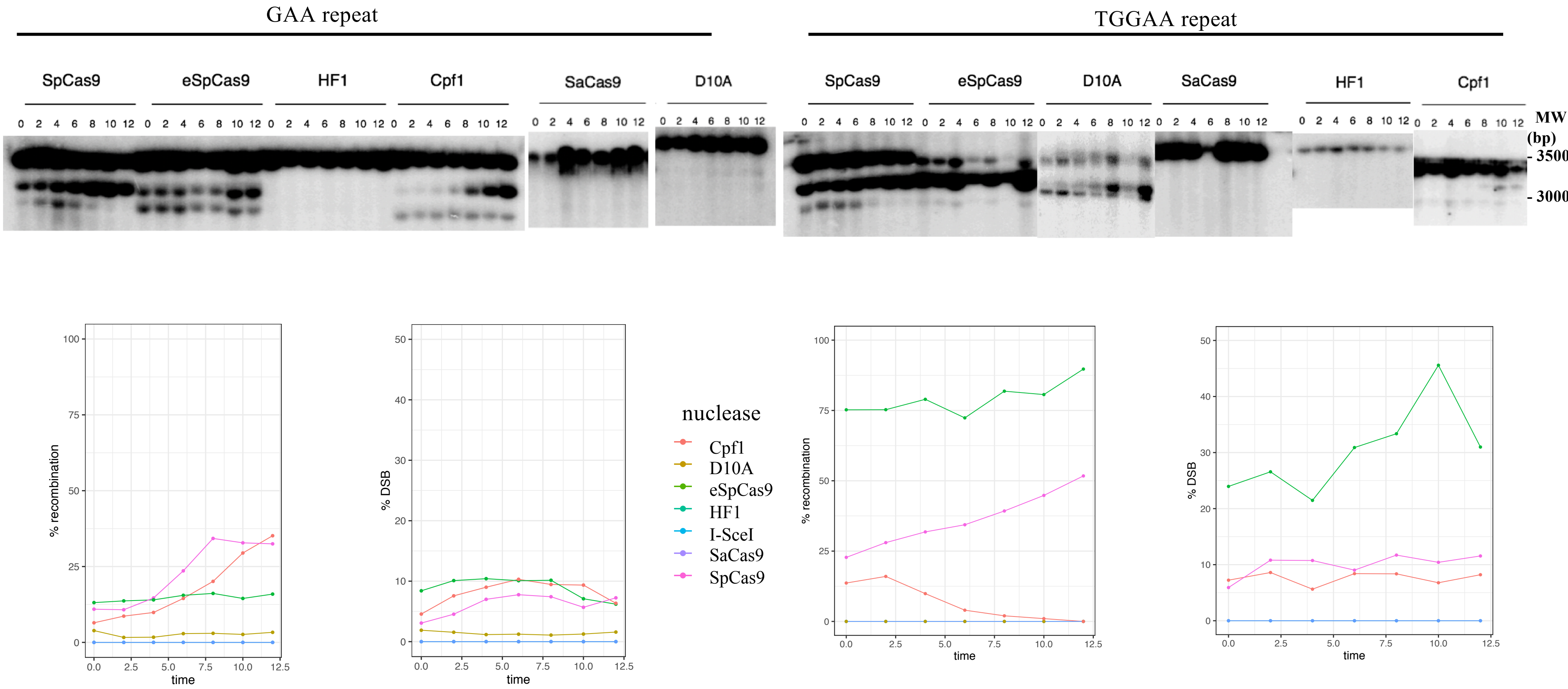


**Poggi *et al.***
**Supplemental Figure S5**

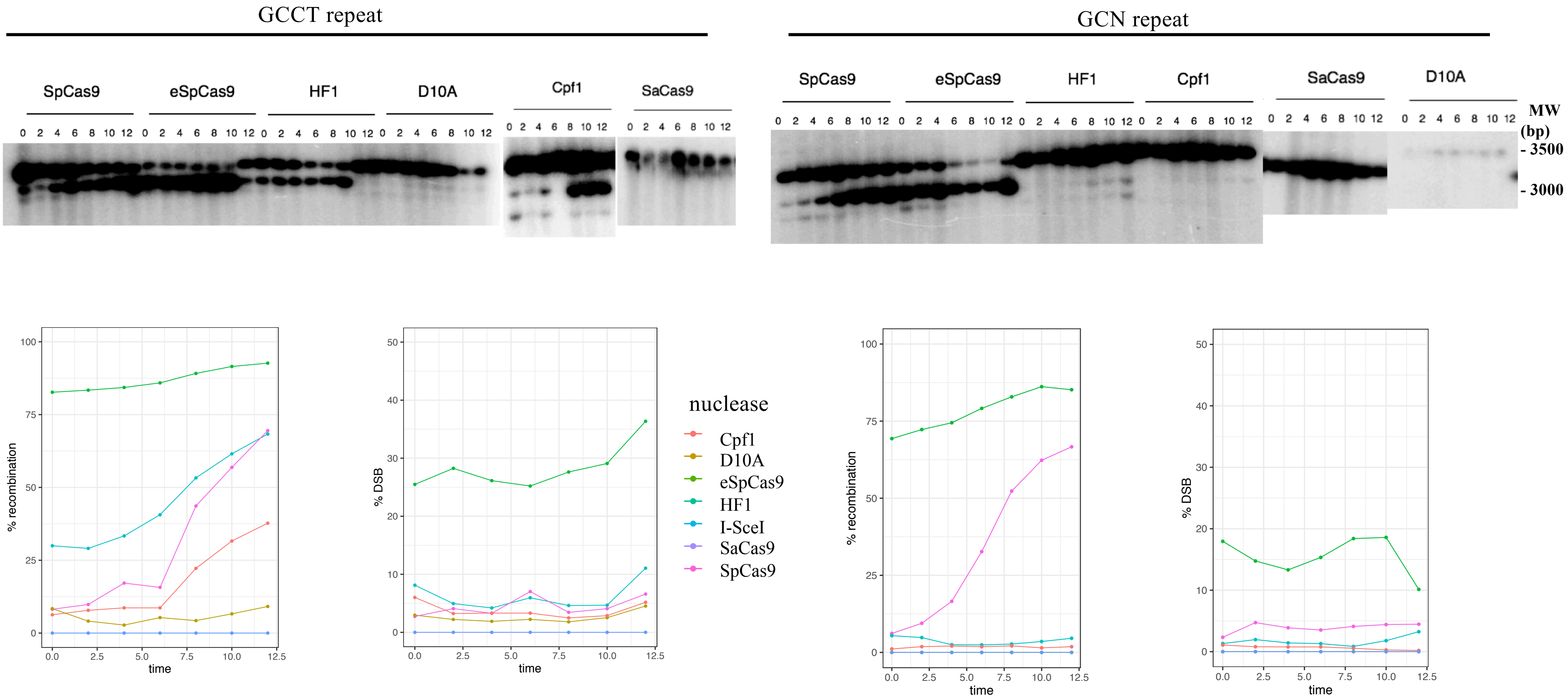


**Poggi *et al.***
**Supplemental Figure S5**

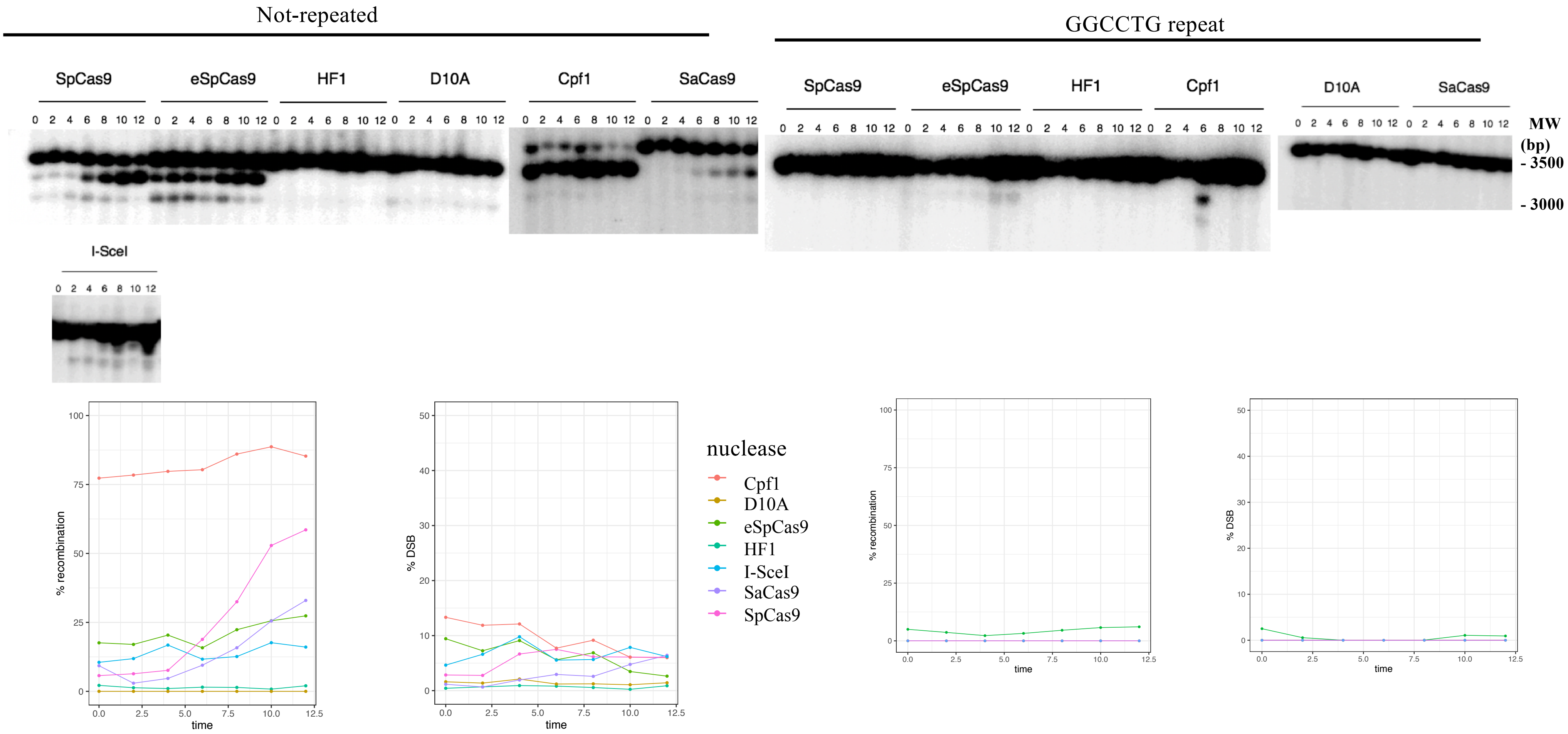


**Poggi *et al.* Supplemental Figure S5**
