## Supplementary material for "Differential efficacies of Cas nucleases on microsatellites involved in human disorders and associated off-target mutations": Sup. Table 1

**Supplemental Table S1: list of plasmids used in this study**

| **Plasmid no.** | **Description** | **Reference** |
| --- | --- | --- |
| pRS416 | URA3 selection marker | (Sikorski and Hieter, 1989) |
| pLPX210 | Sp-Cas9 guide CTG cloned at SpeI-XhoI in pRS416 | This study |
| pLPX209 | Sp-Cas9 guide CAG cloned at SpeI-XhoI in pRS416 | This study |
| pLPX206 | Sp-Cas9 guide TGGAA cloned at SpeI-XhoI in pRS416 | This study |
| pLPX203 | Sp-Cas9 guide GCN cloned at SpeI-XhoI in pRS416 | This study |
| pLPX208 | Sp-Cas9 guide CCTG cloned at SpeI-XhoI in pRS416 | This study |
| pLPX204 | Sp-Cas9 guide CGG cloned at SpeI-XhoI in pRS416 | This study |
| pLPX202 | Sp-Cas9 guide GAA cloned at SpeI-XhoI in pRS416 | This study |
| pLPX207 | Sp-Cas9 guide ATTCT cloned at SpeI-XhoI in pRS416 | This study |
| pLPX201 | Sp-Cas9 guide GGGGCC cloned at SpeI-XhoI in pRS416 | This study |
| pLPX205 | Sp-Cas9 guide GGCCTG cloned at SpeI-XhoI in pRS416 | This study |
| pLPX211 | Sp-Cas9 guide I-SceI cloned at SpeI-XhoI in pRS416 | This study |
| pLPX310 | Cpf1 guide CTG cloned at SpeI-XhoI in pRS416 | This study |
| pLPX309 | Cpf1 guide CAG cloned at SpeI-XhoI in pRS416 | This study |
| pLPX306 | Cpf1guide TGGAA cloned at SpeI-XhoI in pRS416 | This study |
| pLPX303 | Cpf1guide GCN cloned at SpeI-XhoI in pRS416 | This study |
| pLPX308 | Cpf1guide CCTG cloned at SpeI-XhoI in pRS416 | This study |
| pLPX304 | Cpf1guide CGG cloned at SpeI-XhoI in pRS416 | This study |
| pLPX302 | Cpf1guide GAA cloned at SpeI-XhoI in pRS416 | This study |
| pLPX307 | Cpf1guide ATTCT cloned at SpeI-XhoI in pRS416 | This study |
| pLPX301 | Cpf1guide GGGGCC cloned at SpeI-XhoI in pRS416 | This study |
| pLPX305 | Cpf1guide GGCCTG cloned at SpeI-XhoI in pRS416 | This study |
| pLPX311 | Cpf1guide I-SceI cloned at SpeI-XhoI in pRS416 | This study |
| pLPX410 | Sa-Cas9 guide CTG cloned at SpeI-XhoI in pRS416 | This study |
| pLPX409 | Sa-Cas9 guide CAG cloned at SpeI-XhoI in pRS416 | This study |
| pLPX406 | Sa-Cas9 guide TGGAA cloned at SpeI-XhoI in pRS416 | This study |
| pLPX403 | Sa-Cas9 guide GCN cloned at SpeI-XhoI in pRS416 | This study |
| pLPX408 | Sa-Cas9 guide CCTG cloned at SpeI-XhoI in pRS416 | This study |
| pLPX404 | Sa-Cas9 guide CGG cloned at SpeI-XhoI in pRS416 | This study |
| pLPX402 | Sa-Cas9 guide GAA cloned at SpeI-XhoI in pRS416 | This study |
| pLPX407 | Sa-Cas9 guide ATTCT cloned at SpeI-XhoI in pRS416 | This study |
| pLPX401 | Sa-Cas9 guide GGGGCC cloned at SpeI-XhoI in pRS416 | This study |
| pLPX405 | Sa-Cas9 guide GGCCTG cloned at SpeI-XhoI in pRS416 | This study |
| pLPX411 | Sa-Cas9 guide I-SceI cloned at SpeI-XhoI in pRS416 | This study |
| Addgene #43804 | p415-GalL-Cas9-CYC1t  LEU2 selection marker | (DiCarlo et al., 2013) |
| Addgene #72247 | Cas9-HF1 | (Kleinstiver et al., 2016) |
| Addgene #71814 | eSpCas9 | (Slaymaker et al., 2016) |
| Addgene #68706 | Cas9-N863A | (Nishimasu et al., 2014) |
| Addgene #48873 | Cas9-D10A | (Ran et al., 2013) |
| Addgene #84039 | SaCas9 | (Ran et al., 2015) |
| Addgene #69976 | FnCpf1 | (Zetsche et al., 2015) |
| pTRI103 | I-SceI - NLS | (Richard et al., 2003) |
| pLPX10 | pGAL-Cas9-HF1 | This study |
| pLPX11 | pGAL-eSpCas9 | This study |
| pLPX12 | pGAL-N863A | This study |
| pLPX13 | pGAL-D10A | This study |
| pLPX14 | pGAL-SaCas9 | This study |
| pLPX15 | pGAL-FnCpf1 | This study |
| pLPX16 | pGAL-I-SceI | This study |
| synYEGFP | pUC57 backone containing bipartite EGFP gene | This study |
| pLPX110 | bipartite EGFP gene + CTG repeat (100bp) | This study |
| pLPX109 | bipartite EGFP gene + CAG repeat (100bp) | This study |
| pLPX106 | bipartite EGFP gene + TGGGAA repeat (100bp) | This study |
| pLPX103 | bipartite EGFP gene + GCN repeat (100bp) | This study |
| pLPX108 | bipartite EGFP gene + CGG repeat (100bp) | This study |
| pLPX104 | bipartite EGFP gene + CCTG repeat (100bp) | This study |
| pLPX102 | bipartite EGFP gene + ATTCT repeat (100bp) | This study |
| pLPX107 | bipartite EGFP gene + GGGGCC repeat (100bp) | This study |
| pLPX101 | bipartite EGFP gene + GGCCTG repeat (100bp) | This study |
| pLPX105 | bipartite EGFP gene + GAA repeat (100bp) | This study |
