## Supplementary material for "Differential efficacies of Cas nucleases on microsatellites involved in human disorders and associated off-target mutations": Sup. Table 2

**Supplemental Table S2: list and sequence of primers used in this study**

| **Name** | **Oligonucleotide sequence (5’ – 3’)** | **Use** |
| --- | --- | --- |
| JEM1f | TGTGATTTGGCTGAGTTACAACG | qPCR assay |
| JEM1r | AACTGCCCAGCGATCCATT | qPCR assay |
| LP001 | AGTAGTGACTAAGGTTGGCC | qPCR assay |
| LP002 | GGTGAAGGTGATGCTACTTAC | qPCR assay |
| LP003 | GGGGCCTGTTTATTTGTACAAT | qPCR assay |
| LP004 | GATCCAAACGAAAAGAGAGACC | qPCR assay |
| LP400 | CCCCGGATTCTAGAACTAGTGGATCCCCCGGGaaaaaaATGGACTATAAGGACCACGACG | eSpCas9 cloning into pGAL plasmid |
| LP401 | TAAGCGTGACATAACTAATTACATGACTCGAGAAGAGATTACTTTTTCTTTTTTGCCTGGCC |  |
| LP402 | ACCCCGGATTCTAGAACTAGTGGATCCCCCGGGaaaaaaGCCGCCACCATGGATAAAAAG | Cas9-HF1 cloning into pGAL plasmid |
| LP403 | TGTAAGCGTGACATAACTAATTACATGACTCGAGAAGAGAGTCATCCTGCAGCCTTGTCA |  |
| LP408 | AACCCCGGATTCTAGAACTAGTGGATCCCCCGGGaaaaaaatgagcatctaccag | FnCpf1 cloning into pGAL plasmid |
| LP407 | TGTAAGCGTGACATAACTAATTACATGACTCGAGAAGAGAatgagcatctaccag |  |
| LP410 | AACCCCGGATTCTAGAACTAGTGGATCCCCCGGGaaaaaacaccatggccccaaagaagaa | N863A cloning into pGAL plasmid |
| LP411 | TGTAAGCGTGACATAACTAATTACATGACTCGAGAAGAGATTActtgtcatcgtcatcctttttcttttttgcctggcc |  |
| LP412 | AACCCCGGATTCTAGAACTAGTGGATCCCCCGGGaaaaaaCcatggactataaggaccacg | D10A cloning into pGAL plasmid |
| LP413 | TGTAAGCGTGACATAACTAATTACATGACTCGAGAAGAGAttcttactttttcttttttgcc |  |
| LP417 | AACCCCGGATTCTAGAACTAGTGGATCCCCCGGGaaaaaaGGCTGCAGATGCCTCCAAAAAAG | I-SceI cloning into pGAL plasmid |
| LP418 | TGTAAGCGTGACATAACTAATTACATGACTCGAGAAGAGACCCCTCGACTTATTATTTCA |  |
| LP419 | AACCCCGGATTCTAGAACTAGTGGATCCCCCGGGATGGCCCCAAAGAAGAAGCG | SaCas9 cloning into pGAL plasmid |
| LP422 | TGTAAGCGTGACATAACTAATTACATGACTCGAGAAGAGAttactttttcttttttgc |  |
| CAN133 | ACATTTCCACGCCATTTCGC | CAN1 probe |
| CAN135 | GGTTCTAGGTTCGGGTGACG |  |
| SNR44p | GATAACGGACTAGCCTTATTTT | SNR44 probe |
| Spgrnap | GATAACGGACTAGCCTTATTTT | SpCas9 gRNA probe |
| Sagrnap | Tgccttgttttagtagattctg | SaCas9 gRNA probe |
| LP30b | TTCCATGGCCAAC | Verification of cassette integration |
| LP33b | ATGGCTGACAAACAAAA |  |
| P1 | ACACTCTTTCCCTACACGA | Illumina libraries amplification |
| GSP1 | ATACCGTTATTAACATATGACA |  |
| PE1 | AATGATACGGCGACCACCGAGATCTACACTCTTTCCCTACACGACGCTCTTCCGATCT |  |
| GSP2-PE2 | CAAGCAGAAGACGGCATACTACCGTTATTAACATATGACAACTCAA |  |
| PE2 | CAAGCAGAAGACGGCATACGAGATCGGTCTCGGCATTCCTGCTGAACCGCTCTTCCGATCT |  |
