## Supplementary material for "Differential efficacies of Cas nucleases on microsatellites involved in human disorders and associated off-target mutations": Sup. Table 3

**Supplemental Table S3: list of strains used in this study**

| **Name** | **Genotype** | **Origin** |
| --- | --- | --- |
| FYBL1-4D | *MAT*a *ura3*Δ851 *trp*Δ63 *leu2*Δ1 *his3*Δ200 *lys2*Δ202 | FYBL1 spore |
| LPY110 | *MAT*a *ura3*Δ851 *trp*Δ63 *leu2*Δ1 *his3*Δ200 *lys2*Δ202  *can1*Δ:: yEGFP(CTG)_33_-*TRP1* | This study |
| LPY109 | *MAT*a *ura3*Δ851 *trp*Δ63 *leu2*Δ1 *his3*Δ200 *lys2*Δ202  *can1*Δ:: yEGFP(CAG)_33_-*TRP1* | This study |
| LPY106 | *MAT*a *ura3*Δ851 *trp*Δ63 *leu2*Δ1 *his3*Δ200 *lys2*Δ202  *can1*Δ:: yEGFP(TGGGAA)_20_-*TRP1* | This study |
| LPY111 | *MAT*a *ura3*Δ851 *trp*Δ63 *leu2*Δ1 *his3*Δ200 *lys2*Δ202  *can1*Δ:: yEGFP(I-SceI site)-*TRP1* | This study |
| LPY103 | *MAT*a *ura3*Δ851 *trp*Δ63 *leu2*Δ1 *his3*Δ200 *lys2*Δ202  *can1*Δ:: yEGFP(GCN)_33_-*TRP1* | This study |
| LPY108 | *MAT*a *ura3*Δ851 *trp*Δ63 *leu2*Δ1 *his3*Δ200 *lys2*Δ202  *can1*Δ:: yEGFP(CCTG)_25_-*TRP1* | This study |
| LPY104 | *MAT*a *ura3*Δ851 *trp*Δ63 *leu2*Δ1 *his3*Δ200 *lys2*Δ202  *can1*Δ:: yEGFP(CGG)_33_-*TRP1* | This study |
| LPY102 | *MAT*a *ura3*Δ851 *trp*Δ63 *leu2*Δ1 *his3*Δ200 *lys2*Δ202  *can1*Δ:: yEGFP(GAA)_33_-*TRP1* | This study |
| LPY107 | *MAT*a *ura3*Δ851 *trp*Δ63 *leu2*Δ1 *his3*Δ200 *lys2*Δ202  *can1*Δ:: yEGFP(ATTCT)_20_-*TRP1* | This study |
| LPY101 | *MAT*a *ura3*Δ851 *trp*Δ63 *leu2*Δ1 *his3*Δ200 *lys2*Δ202  *can1*Δ:: yEGFP(GGGGCC)_17_-*TRP1* | This study |
| LPY105 | *MAT*a *ura3*Δ851 *trp*Δ63 *leu2*Δ1 *his3*Δ200 *lys2*Δ202  *can1*Δ:: yEGFP(GGCCTG)_17_-*TRP1* | This study |
