## Supplementary material for "Differential efficacies of Cas nucleases on microsatellites involved in human disorders and associated off-target mutations": Sup. Table 4

| **Repeat** | **Nuclease** | **Number of replicas** | **Condition** | **time (h)** | **GFP+% cells** | **standard error** | **standard deviation** | **confidence interval at 95%** |
| --- | --- | --- | --- | --- | --- | --- | --- | --- |
| GGCCTG | Cpf1 | 3 | Gal | 0 | 0.29 | 0.19 | 0.11 | 0.47 |
| GGCCTG | D10A | 3 | Gal | 0 | 0.08 | 0.04 | 0.02 | 0.09 |
| GGCCTG | eSpCas9 | 3 | Gal | 0 | 0.33 | 0.13 | 0.08 | 0.32 |
| GGCCTG | HF1 | 3 | Gal | 0 | 0.38 | 0.28 | 0.16 | 0.70 |
| GGCCTG | N863A | 3 | Gal | 0 | 0.07 | 0.01 | 0.00 | 0.01 |
| GGCCTG | SaCas9 | 3 | Gal | 0 | 0.29 | 0.24 | 0.14 | 0.59 |
| GGCCTG | SpCas9 | 4 | Gal | 0 | 0.19 | 0.12 | 0.06 | 0.20 |
| GGCCTG | Cpf1 | 3 | Gal | 12 | 0.09 | 0.01 | 0.00 | 0.02 |
| GGCCTG | D10A | 3 | Gal | 12 | 0.05 | 0.04 | 0.02 | 0.10 |
| GGCCTG | eSpCas9 | 3 | Gal | 12 | 1.33 | 0.47 | 0.27 | 1.16 |
| GGCCTG | HF1 | 3 | Gal | 12 | 0.09 | 0.03 | 0.02 | 0.07 |
| GGCCTG | N863A | 3 | Gal | 12 | 0.03 | 0.02 | 0.01 | 0.04 |
| GGCCTG | SaCas9 | 3 | Gal | 12 | 0.10 | 0.04 | 0.02 | 0.10 |
| GGCCTG | SpCas9 | 4 | Gal | 12 | 1.13 | 0.73 | 0.37 | 1.17 |
| GGCCTG | Cpf1 | 3 | Gal | 24 | 0.60 | 0.17 | 0.10 | 0.42 |
| GGCCTG | D10A | 3 | Gal | 24 | 0.08 | 0.03 | 0.02 | 0.07 |
| GGCCTG | eSpCas9 | 3 | Gal | 24 | 3.25 | 0.84 | 0.48 | 2.08 |
| GGCCTG | HF1 | 3 | Gal | 24 | 0.20 | 0.08 | 0.05 | 0.21 |
| GGCCTG | N863A | 3 | Gal | 24 | 0.09 | 0.05 | 0.03 | 0.13 |
| GGCCTG | SaCas9 | 3 | Gal | 24 | 0.25 | 0.18 | 0.11 | 0.46 |
| GGCCTG | SpCas9 | 4 | Gal | 24 | 10.19 | 6.34 | 3.17 | 10.09 |
| GGCCTG | Cpf1 | 3 | Gal | 36 | 1.30 | 0.22 | 0.12 | 0.54 |
| GGCCTG | D10A | 3 | Gal | 36 | 0.33 | 0.18 | 0.10 | 0.44 |
| GGCCTG | eSpCas9 | 3 | Gal | 36 | 3.95 | 0.61 | 0.35 | 1.52 |
| GGCCTG | HF1 | 3 | Gal | 36 | 0.45 | 0.22 | 0.13 | 0.55 |
| GGCCTG | N863A | 3 | Gal | 36 | 0.34 | 0.12 | 0.07 | 0.30 |
| GGCCTG | SaCas9 | 3 | Gal | 36 | 0.04 | 0.06 | 0.04 | 0.16 |
| GGCCTG | SpCas9 | 4 | Gal | 36 | 7.44 | 5.23 | 2.61 | 8.32 |
| GGCCTG | Cpf1 | 3 | Glu | 0 | 0.12 | 0.03 | 0.02 | 0.07 |
| GGCCTG | D10A | 3 | Glu | 0 | 0.08 | 0.01 | 0.01 | 0.03 |
| GGCCTG | eSpCas9 | 3 | Glu | 0 | 0.14 | 0.05 | 0.03 | 0.12 |
| GGCCTG | HF1 | 3 | Glu | 0 | 0.23 | 0.15 | 0.09 | 0.37 |
| GGCCTG | N863A | 3 | Glu | 0 | 0.02 | 0.01 | 0.01 | 0.03 |
| GGCCTG | SaCas9 | 3 | Glu | 0 | 0.26 | 0.18 | 0.10 | 0.44 |
| GGCCTG | SpCas9 | 3 | Glu | 0 | 0.17 | 0.06 | 0.03 | 0.14 |
| GGCCTG | Cpf1 | 3 | Glu | 12 | 0.16 | 0.03 | 0.02 | 0.08 |
| GGCCTG | D10A | 3 | Glu | 12 | 0.05 | 0.05 | 0.03 | 0.13 |
| GGCCTG | eSpCas9 | 3 | Glu | 12 | 0.19 | 0.01 | 0.01 | 0.03 |
| GGCCTG | HF1 | 3 | Glu | 12 | 0.15 | 0.01 | 0.01 | 0.03 |
| GGCCTG | N863A | 3 | Glu | 12 | 0.04 | 0.03 | 0.02 | 0.06 |
| GGCCTG | SaCas9 | 3 | Glu | 12 | 0.56 | 0.19 | 0.11 | 0.47 |
| GGCCTG | SpCas9 | 3 | Glu | 12 | 0.17 | 0.05 | 0.03 | 0.12 |
| GGCCTG | Cpf1 | 3 | Glu | 24 | 0.88 | 0.06 | 0.03 | 0.14 |
| GGCCTG | D10A | 3 | Glu | 24 | 0.23 | 0.09 | 0.05 | 0.22 |
| GGCCTG | eSpCas9 | 3 | Glu | 24 | 0.76 | 0.13 | 0.07 | 0.31 |
| GGCCTG | HF1 | 3 | Glu | 24 | 0.69 | 0.30 | 0.17 | 0.74 |
| GGCCTG | N863A | 3 | Glu | 24 | 0.39 | 0.14 | 0.08 | 0.36 |
| GGCCTG | SaCas9 | 3 | Glu | 24 | 1.05 | 0.31 | 0.18 | 0.76 |
| GGCCTG | SpCas9 | 3 | Glu | 24 | 0.64 | 0.23 | 0.13 | 0.57 |
| GGCCTG | Cpf1 | 3 | Glu | 36 | 1.57 | 0.11 | 0.06 | 0.27 |
| GGCCTG | D10A | 3 | Glu | 36 | 0.46 | 0.09 | 0.05 | 0.21 |
| GGCCTG | eSpCas9 | 3 | Glu | 36 | 1.59 | 0.19 | 0.11 | 0.48 |
| GGCCTG | HF1 | 3 | Glu | 36 | 1.01 | 0.18 | 0.10 | 0.44 |
| GGCCTG | N863A | 3 | Glu | 36 | 0.69 | 0.18 | 0.10 | 0.43 |
| GGCCTG | SaCas9 | 3 | Glu | 36 | 0.00 | 0.00 | 0.00 | 0.00 |
| GGCCTG | SpCas9 | 3 | Glu | 36 | 1.04 | 0.12 | 0.07 | 0.30 |
| GGGGCC | Cpf1 | 2 | Gal | 0 | 0.70 | 0.18 | 0.13 | 1.65 |
| GGGGCC | D10A | 3 | Gal | 0 | 2.43 | 2.13 | 1.23 | 5.30 |
| GGGGCC | eSpCas9 | 3 | Gal | 0 | 10.65 | 4.18 | 2.41 | 10.38 |
| GGGGCC | HF1 | 4 | Gal | 0 | 0.47 | 0.38 | 0.19 | 0.61 |
| GGGGCC | N863A | 3 | Gal | 0 | 4.67 | 7.48 | 4.32 | 18.57 |
| GGGGCC | SaCas9 | 3 | Gal | 0 | 0.27 | 0.26 | 0.15 | 0.64 |
| GGGGCC | SpCas9 | 2 | Gal | 0 | 0.14 | 0.11 | 0.08 | 0.99 |
| GGGGCC | Cpf1 | 2 | Gal | 12 | 0.69 | 0.03 | 0.02 | 0.25 |
| GGGGCC | D10A | 3 | Gal | 12 | 24.50 | 3.56 | 2.05 | 8.83 |
| GGGGCC | eSpCas9 | 3 | Gal | 12 | 47.87 | 0.42 | 0.24 | 1.03 |
| GGGGCC | HF1 | 3 | Gal | 12 | 15.68 | 6.57 | 3.80 | 16.33 |
| GGGGCC | N863A | 3 | Gal | 12 | 17.07 | 1.75 | 1.01 | 4.34 |
| GGGGCC | SaCas9 | 3 | Gal | 12 | 0.14 | 0.03 | 0.02 | 0.08 |
| GGGGCC | SpCas9 | 2 | Gal | 12 | 42.15 | 1.91 | 1.35 | 17.15 |
| GGGGCC | Cpf1 | 2 | Gal | 24 | 4.57 | 0.25 | 0.18 | 2.22 |
| GGGGCC | D10A | 3 | Gal | 24 | 51.83 | 3.02 | 1.75 | 7.51 |
| GGGGCC | eSpCas9 | 3 | Gal | 24 | 84.80 | 1.41 | 0.81 | 3.50 |
| GGGGCC | HF1 | 4 | Gal | 24 | 43.18 | 12.16 | 6.08 | 19.35 |
| GGGGCC | N863A | 3 | Gal | 24 | 39.47 | 0.68 | 0.39 | 1.69 |
| GGGGCC | SaCas9 | 3 | Gal | 24 | 0.45 | 0.10 | 0.05 | 0.24 |
| GGGGCC | SpCas9 | 2 | Gal | 24 | 88.20 | 0.42 | 0.30 | 3.81 |
| GGGGCC | Cpf1 | 2 | Gal | 36 | 5.10 | 0.32 | 0.23 | 2.86 |
| GGGGCC | D10A | 3 | Gal | 36 | 51.03 | 3.35 | 1.93 | 8.32 |
| GGGGCC | eSpCas9 | 3 | Gal | 36 | 92.57 | 1.86 | 1.07 | 4.62 |
| GGGGCC | HF1 | 4 | Gal | 36 | 44.55 | 10.74 | 5.37 | 17.09 |
| GGGGCC | N863A | 3 | Gal | 36 | 36.17 | 0.23 | 0.13 | 0.57 |
| GGGGCC | SaCas9 | 3 | Gal | 36 | 1.78 | 0.66 | 0.38 | 1.63 |
| GGGGCC | SpCas9 | 2 | Gal | 36 | 90.25 | 5.59 | 3.95 | 50.19 |
| GGGGCC | Cpf1 | 1 | Glu | 0 | 0.21 | NA | NA | NA |
| GGGGCC | D10A | 3 | Glu | 0 | 2.19 | 2.37 | 1.37 | 5.88 |
| GGGGCC | eSpCas9 | 3 | Glu | 0 | 9.33 | 0.44 | 0.25 | 1.09 |
| GGGGCC | HF1 | 4 | Glu | 0 | 0.39 | 0.29 | 0.15 | 0.46 |
| GGGGCC | N863A | 3 | Glu | 0 | 0.15 | 0.07 | 0.04 | 0.16 |
| GGGGCC | SaCas9 | 3 | Glu | 0 | 0.13 | 0.14 | 0.08 | 0.35 |
| GGGGCC | SpCas9 | 2 | Glu | 0 | 0.24 | 0.05 | 0.04 | 0.44 |
| GGGGCC | Cpf1 | 1 | Glu | 12 | 0.01 | NA | NA | NA |
| GGGGCC | D10A | 3 | Glu | 12 | 3.28 | 2.42 | 1.40 | 6.01 |
| GGGGCC | eSpCas9 | 3 | Glu | 12 | 12.20 | 1.35 | 0.78 | 3.36 |
| GGGGCC | HF1 | 3 | Glu | 12 | 0.26 | 0.05 | 0.03 | 0.13 |
| GGGGCC | N863A | 3 | Glu | 12 | 0.38 | 0.04 | 0.02 | 0.09 |
| GGGGCC | SaCas9 | 3 | Glu | 12 | 0.15 | 0.07 | 0.04 | 0.17 |
| GGGGCC | SpCas9 | 2 | Glu | 12 | 0.30 | 0.04 | 0.03 | 0.32 |
| GGGGCC | Cpf1 | 1 | Glu | 24 | 0.50 | NA | NA | NA |
| GGGGCC | D10A | 3 | Glu | 24 | 4.45 | 3.05 | 1.76 | 7.58 |
| GGGGCC | eSpCas9 | 3 | Glu | 24 | 19.13 | 0.55 | 0.32 | 1.37 |
| GGGGCC | HF1 | 4 | Glu | 24 | 0.55 | 0.33 | 0.16 | 0.52 |
| GGGGCC | N863A | 3 | Glu | 24 | 0.76 | 0.23 | 0.13 | 0.57 |
| GGGGCC | SaCas9 | 3 | Glu | 24 | 0.77 | 0.07 | 0.04 | 0.16 |
| GGGGCC | SpCas9 | 2 | Glu | 24 | 0.55 | 0.05 | 0.04 | 0.44 |
| GGGGCC | Cpf1 | 1 | Glu | 36 | 0.71 | NA | NA | NA |
| GGGGCC | D10A | 3 | Glu | 36 | 4.63 | 2.63 | 1.52 | 6.53 |
| GGGGCC | eSpCas9 | 3 | Glu | 36 | 17.73 | 0.74 | 0.43 | 1.83 |
| GGGGCC | HF1 | 4 | Glu | 36 | 0.74 | 0.27 | 0.14 | 0.43 |
| GGGGCC | N863A | 3 | Glu | 36 | 1.21 | 0.29 | 0.17 | 0.72 |
| GGGGCC | SaCas9 | 3 | Glu | 36 | 0.97 | 1.16 | 0.67 | 2.87 |
| GGGGCC | SpCas9 | 2 | Glu | 36 | 2.43 | 0.70 | 0.50 | 6.29 |
| ATTCT | Cpf1 | 3 | Gal | 0 | 0.04 | 0.05 | 0.03 | 0.12 |
| ATTCT | D10A | 3 | Gal | 0 | 0.10 | 0.04 | 0.02 | 0.09 |
| ATTCT | eSpCas9 | 3 | Gal | 0 | 15.21 | 16.55 | 9.55 | 41.10 |
| ATTCT | HF1 | 4 | Gal | 0 | 0.46 | 0.26 | 0.13 | 0.42 |
| ATTCT | N863A | 3 | Gal | 0 | 0.11 | 0.08 | 0.04 | 0.19 |
| ATTCT | SaCas9 | 3 | Gal | 0 | 0.45 | 0.19 | 0.11 | 0.47 |
| ATTCT | SpCas9 | 3 | Gal | 0 | 0.05 | 0.03 | 0.02 | 0.07 |
| ATTCT | Cpf1 | 3 | Gal | 12 | 1.38 | 0.31 | 0.18 | 0.77 |
| ATTCT | D10A | 3 | Gal | 12 | 0.10 | 0.05 | 0.03 | 0.12 |
| ATTCT | eSpCas9 | 3 | Gal | 12 | 32.33 | 11.00 | 6.35 | 27.32 |
| ATTCT | HF1 | 4 | Gal | 12 | 7.11 | 3.59 | 1.79 | 5.70 |
| ATTCT | N863A | 3 | Gal | 12 | 0.05 | 0.05 | 0.03 | 0.11 |
| ATTCT | SaCas9 | 3 | Gal | 12 | 0.15 | 0.06 | 0.03 | 0.14 |
| ATTCT | SpCas9 | 3 | Gal | 12 | 39.40 | 7.20 | 4.16 | 17.89 |
| ATTCT | Cpf1 | 3 | Gal | 24 | 7.12 | 0.77 | 0.44 | 1.91 |
| ATTCT | D10A | 3 | Gal | 24 | 0.69 | 0.08 | 0.04 | 0.19 |
| ATTCT | eSpCas9 | 3 | Gal | 24 | 70.20 | 4.45 | 2.57 | 11.06 |
| ATTCT | HF1 | 4 | Gal | 24 | 39.70 | 16.90 | 8.45 | 26.90 |
| ATTCT | N863A | 3 | Gal | 24 | 0.12 | 0.09 | 0.05 | 0.21 |
| ATTCT | SaCas9 | 3 | Gal | 24 | 0.61 | 0.47 | 0.27 | 1.18 |
| ATTCT | SpCas9 | 3 | Gal | 24 | 75.83 | 15.55 | 8.98 | 38.62 |
| ATTCT | Cpf1 | 3 | Gal | 36 | 12.03 | 0.81 | 0.47 | 2.01 |
| ATTCT | D10A | 3 | Gal | 36 | 0.91 | 0.31 | 0.18 | 0.77 |
| ATTCT | eSpCas9 | 3 | Gal | 36 | 78.43 | 8.87 | 5.12 | 22.03 |
| ATTCT | HF1 | 4 | Gal | 36 | 49.25 | 19.05 | 9.52 | 30.31 |
| ATTCT | N863A | 3 | Gal | 36 | 0.56 | 0.15 | 0.08 | 0.36 |
| ATTCT | SaCas9 | 3 | Gal | 36 | 2.50 | 0.93 | 0.54 | 2.31 |
| ATTCT | SpCas9 | 3 | Gal | 36 | 83.43 | 15.17 | 8.76 | 37.68 |
| ATTCT | Cpf1 | 3 | Glu | 0 | 0.03 | 0.04 | 0.02 | 0.09 |
| ATTCT | D10A | 3 | Glu | 0 | 0.09 | 0.11 | 0.06 | 0.27 |
| ATTCT | eSpCas9 | 3 | Glu | 0 | 6.93 | 0.93 | 0.54 | 2.32 |
| ATTCT | HF1 | 4 | Glu | 0 | 0.44 | 0.56 | 0.28 | 0.89 |
| ATTCT | N863A | 3 | Glu | 0 | 0.10 | 0.05 | 0.03 | 0.12 |
| ATTCT | SaCas9 | 3 | Glu | 0 | 0.12 | 0.03 | 0.02 | 0.08 |
| ATTCT | SpCas9 | 7 | Glu | 0 | 0.10 | 0.04 | 0.02 | 0.04 |
| ATTCT | Cpf1 | 3 | Glu | 12 | 0.06 | 0.08 | 0.05 | 0.20 |
| ATTCT | D10A | 3 | Glu | 12 | 0.03 | 0.02 | 0.01 | 0.05 |
| ATTCT | eSpCas9 | 3 | Glu | 12 | 14.70 | 1.35 | 0.78 | 3.34 |
| ATTCT | HF1 | 4 | Glu | 12 | 0.61 | 0.56 | 0.28 | 0.89 |
| ATTCT | N863A | 3 | Glu | 12 | 0.02 | 0.04 | 0.02 | 0.10 |
| ATTCT | SaCas9 | 3 | Glu | 12 | 0.34 | 0.26 | 0.15 | 0.64 |
| ATTCT | SpCas9 | 7 | Glu | 12 | 0.43 | 0.38 | 0.15 | 0.36 |
| ATTCT | Cpf1 | 3 | Glu | 24 | 0.70 | 0.12 | 0.07 | 0.29 |
| ATTCT | D10A | 3 | Glu | 24 | 0.36 | 0.05 | 0.03 | 0.11 |
| ATTCT | eSpCas9 | 3 | Glu | 24 | 17.70 | 1.28 | 0.74 | 3.17 |
| ATTCT | HF1 | 4 | Glu | 24 | 0.93 | 0.44 | 0.22 | 0.70 |
| ATTCT | N863A | 3 | Glu | 24 | 0.26 | 0.12 | 0.07 | 0.29 |
| ATTCT | SaCas9 | 3 | Glu | 24 | 1.47 | 0.36 | 0.21 | 0.89 |
| ATTCT | SpCas9 | 7 | Glu | 24 | 0.75 | 0.11 | 0.04 | 0.10 |
| ATTCT | Cpf1 | 3 | Glu | 36 | 0.53 | 0.20 | 0.12 | 0.50 |
| ATTCT | D10A | 3 | Glu | 36 | 0.78 | 0.14 | 0.08 | 0.35 |
| ATTCT | eSpCas9 | 3 | Glu | 36 | 19.33 | 1.66 | 0.96 | 4.13 |
| ATTCT | HF1 | 4 | Glu | 36 | 1.93 | 0.84 | 0.42 | 1.34 |
| ATTCT | N863A | 3 | Glu | 36 | 0.65 | 0.13 | 0.07 | 0.32 |
| ATTCT | SaCas9 | 3 | Glu | 36 | 2.47 | 0.46 | 0.27 | 1.15 |
| ATTCT | SpCas9 | 7 | Glu | 36 | 1.91 | 0.75 | 0.28 | 0.69 |
| CAG | Cpf1 | 5 | Gal | 0 | 0.07 | 0.02 | 0.01 | 0.03 |
| CAG | D10A | 3 | Gal | 0 | 0.09 | 0.04 | 0.02 | 0.10 |
| CAG | eSpCas9 | 7 | Gal | 0 | 1.20 | 1.61 | 0.61 | 1.49 |
| CAG | HF1 | 3 | Gal | 0 | 0.07 | 0.04 | 0.03 | 0.11 |
| CAG | N863A | 3 | Gal | 0 | 0.09 | 0.04 | 0.02 | 0.09 |
| CAG | SaCas9 | 3 | Gal | 0 | 0.32 | 0.06 | 0.03 | 0.14 |
| CAG | SpCas9 | 4 | Gal | 0 | 0.13 | 0.13 | 0.07 | 0.21 |
| CAG | Cpf1 | 5 | Gal | 12 | 4.22 | 1.11 | 0.50 | 1.38 |
| CAG | D10A | 3 | Gal | 12 | 1.17 | 0.03 | 0.02 | 0.08 |
| CAG | eSpCas9 | 4 | Gal | 12 | 5.11 | 3.07 | 1.53 | 4.88 |
| CAG | HF1 | 1 | Gal | 12 | 0.06 | NA | NA | NA |
| CAG | N863A | 3 | Gal | 12 | 0.10 | 0.03 | 0.02 | 0.08 |
| CAG | SaCas9 | 3 | Gal | 12 | 0.20 | 0.08 | 0.05 | 0.20 |
| CAG | SpCas9 | 3 | Gal | 12 | 5.41 | 1.09 | 0.63 | 2.71 |
| CAG | Cpf1 | 5 | Gal | 24 | 5.61 | 0.85 | 0.38 | 1.05 |
| CAG | D10A | 3 | Gal | 24 | 4.25 | 1.28 | 0.74 | 3.19 |
| CAG | eSpCas9 | 7 | Gal | 24 | 13.17 | 4.15 | 1.57 | 3.83 |
| CAG | HF1 | 3 | Gal | 24 | 0.34 | 0.15 | 0.08 | 0.36 |
| CAG | N863A | 3 | Gal | 24 | 0.16 | 0.06 | 0.03 | 0.14 |
| CAG | SaCas9 | 3 | Gal | 24 | 0.17 | 0.03 | 0.02 | 0.08 |
| CAG | SpCas9 | 4 | Gal | 24 | 6.89 | 5.10 | 2.55 | 8.11 |
| CAG | Cpf1 | 5 | Gal | 36 | 5.23 | 0.75 | 0.34 | 0.93 |
| CAG | D10A | 3 | Gal | 36 | 4.37 | 1.03 | 0.60 | 2.57 |
| CAG | eSpCas9 | 7 | Gal | 36 | 20.64 | 5.40 | 2.04 | 4.99 |
| CAG | HF1 | 3 | Gal | 36 | 0.39 | 0.14 | 0.08 | 0.35 |
| CAG | N863A | 3 | Gal | 36 | 0.38 | 0.04 | 0.02 | 0.10 |
| CAG | SaCas9 | 3 | Gal | 36 | 1.15 | 0.49 | 0.28 | 1.22 |
| CAG | SpCas9 | 4 | Gal | 36 | 8.68 | 9.28 | 4.64 | 14.76 |
| CAG | Cpf1 | 5 | Glu | 0 | 0.10 | 0.06 | 0.03 | 0.08 |
| CAG | D10A | 3 | Glu | 0 | 0.14 | 0.07 | 0.04 | 0.18 |
| CAG | eSpCas9 | 2 | Glu | 0 | 2.33 | 2.47 | 1.75 | 22.24 |
| CAG | HF1 | 1 | Glu | 0 | 0.04 | NA | NA | NA |
| CAG | N863A | 2 | Glu | 0 | 0.06 | 0.06 | 0.04 | 0.51 |
| CAG | SaCas9 | 3 | Glu | 0 | 0.16 | 0.01 | 0.00 | 0.01 |
| CAG | SpCas9 | 4 | Glu | 0 | 0.37 | 0.61 | 0.31 | 0.97 |
| CAG | Cpf1 | 5 | Glu | 12 | 0.17 | 0.11 | 0.05 | 0.14 |
| CAG | D10A | 3 | Glu | 12 | 0.11 | 0.04 | 0.02 | 0.10 |
| CAG | eSpCas9 | 1 | Glu | 12 | 3.81 | NA | NA | NA |
| CAG | HF1 | 1 | Glu | 12 | 0.07 | NA | NA | NA |
| CAG | N863A | 2 | Glu | 12 | 0.00 | 0.00 | 0.00 | 0.00 |
| CAG | SaCas9 | 3 | Glu | 12 | 0.34 | 0.11 | 0.06 | 0.28 |
| CAG | SpCas9 | 3 | Glu | 12 | 0.11 | 0.03 | 0.01 | 0.06 |
| CAG | Cpf1 | 5 | Glu | 24 | 0.25 | 0.10 | 0.04 | 0.12 |
| CAG | D10A | 3 | Glu | 24 | 0.27 | 0.05 | 0.03 | 0.13 |
| CAG | eSpCas9 | 2 | Glu | 24 | 2.77 | 2.14 | 1.52 | 19.25 |
| CAG | HF1 | 1 | Glu | 24 | 0.15 | NA | NA | NA |
| CAG | N863A | 2 | Glu | 24 | 0.15 | 0.02 | 0.02 | 0.19 |
| CAG | SaCas9 | 3 | Glu | 24 | 1.47 | 0.29 | 0.16 | 0.71 |
| CAG | SpCas9 | 4 | Glu | 24 | 0.27 | 0.07 | 0.03 | 0.11 |
| CAG | Cpf1 | 5 | Glu | 36 | 0.59 | 0.23 | 0.10 | 0.29 |
| CAG | D10A | 3 | Glu | 36 | 0.65 | 0.23 | 0.13 | 0.57 |
| CAG | eSpCas9 | 2 | Glu | 36 | 4.99 | 3.25 | 2.30 | 29.16 |
| CAG | HF1 | 1 | Glu | 36 | 0.21 | NA | NA | NA |
| CAG | N863A | 2 | Glu | 36 | 0.41 | 0.01 | 0.00 | 0.06 |
| CAG | SaCas9 | 3 | Glu | 36 | 2.26 | 0.19 | 0.11 | 0.47 |
| CAG | SpCas9 | 4 | Glu | 36 | 0.96 | 0.55 | 0.28 | 0.88 |
| CTG | Cpf1 | 3 | Gal | 0 | 0.03 | 0.01 | 0.01 | 0.03 |
| CTG | D10A | 6 | Gal | 0 | 0.15 | 0.10 | 0.04 | 0.10 |
| CTG | eSpCas9 | 3 | Gal | 0 | 1.31 | 0.67 | 0.38 | 1.65 |
| CTG | HF1 | 3 | Gal | 0 | 0.03 | 0.03 | 0.02 | 0.07 |
| CTG | N863A | 6 | Gal | 0 | 0.15 | 0.07 | 0.03 | 0.07 |
| CTG | SaCas9 | 6 | Gal | 0 | 2.07 | 3.82 | 1.56 | 4.01 |
| CTG | SpCas9 | 8 | Gal | 0 | 0.07 | 0.06 | 0.02 | 0.05 |
| CTG | Cpf1 | 3 | Gal | 12 | 0.77 | 0.32 | 0.18 | 0.79 |
| CTG | D10A | 6 | Gal | 12 | 0.06 | 0.02 | 0.01 | 0.02 |
| CTG | eSpCas9 | 3 | Gal | 12 | 2.90 | 2.10 | 1.21 | 5.22 |
| CTG | HF1 | 3 | Gal | 12 | 0.12 | 0.07 | 0.04 | 0.17 |
| CTG | N863A | 6 | Gal | 12 | 0.05 | 0.02 | 0.01 | 0.02 |
| CTG | SaCas9 | 6 | Gal | 12 | 1.57 | 3.47 | 1.42 | 3.64 |
| CTG | SpCas9 | 8 | Gal | 12 | 10.86 | 3.73 | 1.32 | 3.12 |
| CTG | Cpf1 | 3 | Gal | 24 | 2.38 | 0.71 | 0.41 | 1.77 |
| CTG | D10A | 6 | Gal | 24 | 0.14 | 0.07 | 0.03 | 0.07 |
| CTG | eSpCas9 | 3 | Gal | 24 | 4.62 | 2.93 | 1.69 | 7.27 |
| CTG | HF1 | 3 | Gal | 24 | 0.15 | 0.07 | 0.04 | 0.19 |
| CTG | N863A | 6 | Gal | 24 | 0.06 | 0.03 | 0.01 | 0.03 |
| CTG | SaCas9 | 6 | Gal | 24 | 1.80 | 4.02 | 1.64 | 4.22 |
| CTG | SpCas9 | 8 | Gal | 24 | 21.41 | 2.11 | 0.75 | 1.77 |
| CTG | Cpf1 | 3 | Gal | 36 | 3.00 | 0.30 | 0.17 | 0.74 |
| CTG | D10A | 6 | Gal | 36 | 0.49 | 0.14 | 0.06 | 0.15 |
| CTG | eSpCas9 | 3 | Gal | 36 | 6.57 | 3.06 | 1.77 | 7.60 |
| CTG | HF1 | 3 | Gal | 36 | 0.22 | 0.11 | 0.07 | 0.28 |
| CTG | N863A | 6 | Gal | 36 | 0.41 | 0.20 | 0.08 | 0.21 |
| CTG | SaCas9 | 6 | Gal | 36 | 2.49 | 3.26 | 1.33 | 3.42 |
| CTG | SpCas9 | 8 | Gal | 36 | 22.06 | 1.97 | 0.70 | 1.64 |
| CTG | Cpf1 | 3 | Glu | 0 | 0.01 | 0.01 | 0.01 | 0.03 |
| CTG | D10A | 6 | Glu | 0 | 0.10 | 0.06 | 0.03 | 0.06 |
| CTG | eSpCas9 | 3 | Glu | 0 | 0.81 | 0.56 | 0.32 | 1.38 |
| CTG | HF1 | 3 | Glu | 0 | 0.03 | 0.02 | 0.01 | 0.05 |
| CTG | N863A | 6 | Glu | 0 | 0.10 | 0.02 | 0.01 | 0.02 |
| CTG | SaCas9 | 6 | Glu | 0 | 1.01 | 2.17 | 0.89 | 2.28 |
| CTG | SpCas9 | 8 | Glu | 0 | 0.26 | 0.53 | 0.19 | 0.44 |
| CTG | Cpf1 | 3 | Glu | 12 | 0.37 | 0.16 | 0.09 | 0.40 |
| CTG | D10A | 6 | Glu | 12 | 0.04 | 0.03 | 0.01 | 0.03 |
| CTG | eSpCas9 | 3 | Glu | 12 | 2.09 | 0.78 | 0.45 | 1.94 |
| CTG | HF1 | 3 | Glu | 12 | 0.37 | 0.29 | 0.17 | 0.73 |
| CTG | N863A | 6 | Glu | 12 | 0.06 | 0.06 | 0.02 | 0.06 |
| CTG | SaCas9 | 6 | Glu | 12 | 0.18 | 0.13 | 0.05 | 0.14 |
| CTG | SpCas9 | 8 | Glu | 12 | 1.04 | 2.28 | 0.80 | 1.90 |
| CTG | Cpf1 | 3 | Glu | 24 | 0.29 | 0.10 | 0.06 | 0.25 |
| CTG | D10A | 6 | Glu | 24 | 0.60 | 0.16 | 0.07 | 0.17 |
| CTG | eSpCas9 | 3 | Glu | 24 | 2.56 | 1.16 | 0.67 | 2.87 |
| CTG | HF1 | 3 | Glu | 24 | 0.34 | 0.16 | 0.09 | 0.40 |
| CTG | N863A | 6 | Glu | 24 | 0.85 | 0.64 | 0.26 | 0.68 |
| CTG | SaCas9 | 6 | Glu | 24 | 1.45 | 0.51 | 0.21 | 0.54 |
| CTG | SpCas9 | 8 | Glu | 24 | 0.61 | 0.45 | 0.16 | 0.38 |
| CTG | Cpf1 | 3 | Glu | 36 | 0.76 | 0.16 | 0.09 | 0.39 |
| CTG | D10A | 6 | Glu | 36 | 1.32 | 0.28 | 0.12 | 0.30 |
| CTG | eSpCas9 | 3 | Glu | 36 | 4.84 | 2.25 | 1.30 | 5.59 |
| CTG | HF1 | 3 | Glu | 36 | 0.81 | 0.26 | 0.15 | 0.64 |
| CTG | N863A | 6 | Glu | 36 | 1.83 | 1.13 | 0.46 | 1.18 |
| CTG | SaCas9 | 6 | Glu | 36 | 2.79 | 0.98 | 0.40 | 1.02 |
| CTG | SpCas9 | 8 | Glu | 36 | 1.57 | 1.14 | 0.40 | 0.96 |
| GAA | Cpf1 | 6 | Gal | 0 | 1.97 | 4.23 | 1.73 | 4.44 |
| GAA | D10A | 3 | Gal | 0 | 0.31 | 0.07 | 0.04 | 0.17 |
| GAA | eSpCas9 | 3 | Gal | 0 | 3.48 | 3.08 | 1.78 | 7.66 |
| GAA | HF1 | 3 | Gal | 0 | 0.12 | 0.09 | 0.05 | 0.22 |
| GAA | N863A | 3 | Gal | 0 | 0.08 | 0.01 | 0.01 | 0.03 |
| GAA | SaCas9 | 3 | Gal | 0 | 0.16 | 0.11 | 0.06 | 0.27 |
| GAA | SpCas9 | 6 | Gal | 0 | 0.21 | 0.09 | 0.04 | 0.10 |
| GAA | Cpf1 | 3 | Gal | 12 | 32.17 | 9.72 | 5.61 | 24.15 |
| GAA | D10A | 3 | Gal | 12 | 1.77 | 0.10 | 0.06 | 0.26 |
| GAA | eSpCas9 | 3 | Gal | 12 | 8.36 | 4.94 | 2.85 | 12.28 |
| GAA | HF1 | 3 | Gal | 12 | 0.06 | 0.03 | 0.02 | 0.09 |
| GAA | N863A | 3 | Gal | 12 | 0.09 | 0.06 | 0.04 | 0.15 |
| GAA | SaCas9 | 3 | Gal | 12 | 0.08 | 0.01 | 0.00 | 0.02 |
| GAA | SpCas9 | 6 | Gal | 12 | 24.00 | 9.00 | 3.67 | 9.44 |
| GAA | Cpf1 | 6 | Gal | 24 | 59.75 | 9.01 | 3.68 | 9.45 |
| GAA | D10A | 3 | Gal | 24 | 5.31 | 0.36 | 0.21 | 0.91 |
| GAA | eSpCas9 | 3 | Gal | 24 | 21.10 | 7.57 | 4.37 | 18.81 |
| GAA | HF1 | 3 | Gal | 24 | 0.25 | 0.03 | 0.02 | 0.07 |
| GAA | N863A | 3 | Gal | 24 | 0.37 | 0.23 | 0.13 | 0.56 |
| GAA | SaCas9 | 3 | Gal | 24 | 0.85 | 0.51 | 0.29 | 1.27 |
| GAA | SpCas9 | 6 | Gal | 24 | 19.92 | 1.89 | 0.77 | 1.98 |
| GAA | Cpf1 | 6 | Gal | 36 | 61.52 | 6.25 | 2.55 | 6.56 |
| GAA | D10A | 3 | Gal | 36 | 5.24 | 0.06 | 0.04 | 0.16 |
| GAA | eSpCas9 | 3 | Gal | 36 | 22.57 | 8.36 | 4.83 | 20.76 |
| GAA | HF1 | 3 | Gal | 36 | 0.53 | 0.07 | 0.04 | 0.18 |
| GAA | N863A | 3 | Gal | 36 | 0.62 | 0.19 | 0.11 | 0.46 |
| GAA | SaCas9 | 3 | Gal | 36 | 3.26 | 0.85 | 0.49 | 2.10 |
| GAA | SpCas9 | 6 | Gal | 36 | 25.38 | 2.80 | 1.14 | 2.93 |
| GAA | Cpf1 | 4 | Glu | 0 | 0.68 | 0.56 | 0.28 | 0.89 |
| GAA | D10A | 3 | Glu | 0 | 0.09 | 0.04 | 0.02 | 0.09 |
| GAA | eSpCas9 | 3 | Glu | 0 | 1.63 | 0.77 | 0.44 | 1.90 |
| GAA | HF1 | 3 | Glu | 0 | 0.20 | 0.13 | 0.07 | 0.32 |
| GAA | N863A | 3 | Glu | 0 | 0.03 | 0.02 | 0.01 | 0.05 |
| GAA | SaCas9 | 3 | Glu | 0 | 0.09 | 0.04 | 0.02 | 0.09 |
| GAA | SpCas9 | 6 | Glu | 0 | 4.24 | 9.68 | 3.95 | 10.16 |
| GAA | Cpf1 | 3 | Glu | 12 | 0.66 | 0.20 | 0.11 | 0.48 |
| GAA | D10A | 3 | Glu | 12 | 0.33 | 0.07 | 0.04 | 0.17 |
| GAA | eSpCas9 | 3 | Glu | 12 | 2.14 | 0.41 | 0.23 | 1.01 |
| GAA | HF1 | 3 | Glu | 12 | 0.11 | 0.03 | 0.02 | 0.08 |
| GAA | N863A | 3 | Glu | 12 | 0.11 | 0.08 | 0.04 | 0.19 |
| GAA | SaCas9 | 3 | Glu | 12 | 0.28 | 0.19 | 0.11 | 0.47 |
| GAA | SpCas9 | 6 | Glu | 12 | 4.92 | 10.82 | 4.42 | 11.35 |
| GAA | Cpf1 | 3 | Glu | 24 | 1.32 | 0.09 | 0.05 | 0.22 |
| GAA | D10A | 3 | Glu | 24 | 0.69 | 0.07 | 0.04 | 0.17 |
| GAA | eSpCas9 | 3 | Glu | 24 | 3.20 | 0.81 | 0.47 | 2.02 |
| GAA | HF1 | 3 | Glu | 24 | 0.42 | 0.07 | 0.04 | 0.18 |
| GAA | N863A | 3 | Glu | 24 | 0.30 | 0.09 | 0.05 | 0.23 |
| GAA | SaCas9 | 3 | Glu | 24 | 0.74 | 0.29 | 0.17 | 0.72 |
| GAA | SpCas9 | 6 | Glu | 24 | 5.73 | 11.80 | 4.82 | 12.38 |
| GAA | Cpf1 | 6 | Glu | 36 | 1.49 | 0.86 | 0.35 | 0.90 |
| GAA | D10A | 3 | Glu | 36 | 1.26 | 0.16 | 0.09 | 0.39 |
| GAA | eSpCas9 | 3 | Glu | 36 | 4.36 | 0.84 | 0.49 | 2.09 |
| GAA | HF1 | 3 | Glu | 36 | 0.68 | 0.18 | 0.10 | 0.45 |
| GAA | N863A | 3 | Glu | 36 | 0.92 | 0.18 | 0.10 | 0.44 |
| GAA | SaCas9 | 3 | Glu | 36 | 2.59 | 1.24 | 0.72 | 3.08 |
| GAA | SpCas9 | 6 | Glu | 36 | 6.84 | 11.84 | 4.83 | 12.43 |
| CGG | Cpf1 | 2 | Gal | 0 | 0.43 | 0.55 | 0.39 | 4.93 |
| CGG | D10A | 4 | Gal | 0 | 2.88 | 2.61 | 1.30 | 4.15 |
| CGG | eSpCas9 | 6 | Gal | 0 | 35.90 | 18.16 | 7.41 | 19.05 |
| CGG | HF1 | 3 | Gal | 0 | 0.65 | 0.11 | 0.06 | 0.27 |
| CGG | N863A | 2 | Gal | 0 | 0.08 | 0.02 | 0.02 | 0.22 |
| CGG | SaCas9 | 3 | Gal | 0 | 0.59 | 0.34 | 0.20 | 0.85 |
| CGG | SpCas9 | 5 | Gal | 0 | 2.55 | 3.15 | 1.41 | 3.91 |
| CGG | Cpf1 | 2 | Gal | 12 | 0.17 | 0.05 | 0.04 | 0.44 |
| CGG | D10A | 4 | Gal | 12 | 15.18 | 4.19 | 2.10 | 6.67 |
| CGG | eSpCas9 | 6 | Gal | 12 | 64.68 | 12.66 | 5.17 | 13.28 |
| CGG | HF1 | 3 | Gal | 12 | 8.96 | 1.50 | 0.86 | 3.71 |
| CGG | N863A | 2 | Gal | 12 | 5.44 | 0.69 | 0.49 | 6.23 |
| CGG | SaCas9 | 3 | Gal | 12 | 0.23 | 0.15 | 0.08 | 0.36 |
| CGG | SpCas9 | 5 | Gal | 12 | 36.38 | 8.90 | 3.98 | 11.05 |
| CGG | Cpf1 | 2 | Gal | 24 | 0.45 | 0.07 | 0.05 | 0.64 |
| CGG | D10A | 4 | Gal | 24 | 45.95 | 11.78 | 5.89 | 18.75 |
| CGG | eSpCas9 | 6 | Gal | 24 | 85.73 | 3.66 | 1.50 | 3.84 |
| CGG | HF1 | 3 | Gal | 24 | 43.00 | 1.59 | 0.92 | 3.94 |
| CGG | N863A | 2 | Gal | 24 | 25.70 | 0.00 | 0.00 | 0.00 |
| CGG | SaCas9 | 3 | Gal | 24 | 0.40 | 0.31 | 0.18 | 0.78 |
| CGG | SpCas9 | 5 | Gal | 24 | 87.00 | 1.30 | 0.58 | 1.62 |
| CGG | Cpf1 | 2 | Gal | 36 | 0.93 | 0.42 | 0.30 | 3.81 |
| CGG | D10A | 4 | Gal | 36 | 58.20 | 14.50 | 7.25 | 23.07 |
| CGG | eSpCas9 | 6 | Gal | 36 | 88.25 | 4.45 | 1.82 | 4.67 |
| CGG | HF1 | 3 | Gal | 36 | 57.80 | 3.85 | 2.22 | 9.57 |
| CGG | N863A | 2 | Gal | 36 | 31.80 | 2.97 | 2.10 | 26.68 |
| CGG | SaCas9 | 3 | Gal | 36 | 0.00 | 0.00 | 0.00 | 0.00 |
| CGG | SpCas9 | 5 | Gal | 36 | 93.10 | 2.13 | 0.95 | 2.64 |
| CGG | Cpf1 | 1 | Glu | 0 | 0.01 | NA | NA | NA |
| CGG | D10A | 3 | Glu | 0 | 5.78 | 4.31 | 2.49 | 10.70 |
| CGG | eSpCas9 | 6 | Glu | 0 | 48.17 | 30.02 | 12.25 | 31.50 |
| CGG | HF1 | 3 | Glu | 0 | 0.76 | 0.48 | 0.28 | 1.19 |
| CGG | N863A | 2 | Glu | 0 | 0.23 | 0.07 | 0.05 | 0.64 |
| CGG | SaCas9 | 3 | Glu | 0 | 0.26 | 0.38 | 0.22 | 0.94 |
| CGG | SpCas9 | 5 | Glu | 0 | 12.20 | 20.29 | 9.07 | 25.19 |
| CGG | Cpf1 | 1 | Glu | 12 | 0.03 | NA | NA | NA |
| CGG | D10A | 3 | Glu | 12 | 11.43 | 7.21 | 4.16 | 17.91 |
| CGG | eSpCas9 | 6 | Glu | 12 | 57.48 | 25.43 | 10.38 | 26.69 |
| CGG | HF1 | 3 | Glu | 12 | 1.10 | 0.33 | 0.19 | 0.82 |
| CGG | N863A | 2 | Glu | 12 | 0.43 | 0.32 | 0.23 | 2.86 |
| CGG | SaCas9 | 3 | Glu | 12 | 0.60 | 0.19 | 0.11 | 0.48 |
| CGG | SpCas9 | 5 | Glu | 12 | 14.46 | 22.30 | 9.97 | 27.69 |
| CGG | Cpf1 | 1 | Glu | 24 | 0.38 | NA | NA | NA |
| CGG | D10A | 3 | Glu | 24 | 13.91 | 7.19 | 4.15 | 17.87 |
| CGG | eSpCas9 | 6 | Glu | 24 | 58.93 | 24.69 | 10.08 | 25.91 |
| CGG | HF1 | 3 | Glu | 24 | 2.03 | 0.64 | 0.37 | 1.58 |
| CGG | N863A | 2 | Glu | 24 | 0.39 | 0.06 | 0.05 | 0.57 |
| CGG | SaCas9 | 3 | Glu | 24 | 1.15 | 0.11 | 0.06 | 0.27 |
| CGG | SpCas9 | 5 | Glu | 24 | 15.59 | 22.11 | 9.89 | 27.46 |
| CGG | Cpf1 | 1 | Glu | 36 | 0.69 | NA | NA | NA |
| CGG | D10A | 3 | Glu | 36 | 15.29 | 6.99 | 4.04 | 17.37 |
| CGG | eSpCas9 | 6 | Glu | 36 | 61.30 | 22.13 | 9.03 | 23.22 |
| CGG | HF1 | 3 | Glu | 36 | 3.45 | 1.38 | 0.80 | 3.44 |
| CGG | N863A | 2 | Glu | 36 | 0.88 | 0.02 | 0.02 | 0.19 |
| CGG | SaCas9 | 3 | Glu | 36 | 0.00 | 0.00 | 0.00 | 0.00 |
| CGG | SpCas9 | 5 | Glu | 36 | 17.38 | 20.97 | 9.38 | 26.04 |
| CCTG | Cpf1 | 6 | Gal | 0 | 0.24 | 0.13 | 0.05 | 0.13 |
| CCTG | D10A | 3 | Gal | 0 | 0.17 | 0.08 | 0.05 | 0.20 |
| CCTG | eSpCas9 | 4 | Gal | 0 | 21.60 | 9.70 | 4.85 | 15.44 |
| CCTG | HF1 | 3 | Gal | 0 | 0.03 | 0.01 | 0.00 | 0.02 |
| CCTG | N863A | 3 | Gal | 0 | 0.16 | 0.11 | 0.06 | 0.26 |
| CCTG | SaCas9 | 3 | Gal | 0 | 0.26 | 0.16 | 0.09 | 0.39 |
| CCTG | SpCas9 | 3 | Gal | 0 | 10.11 | 17.23 | 9.95 | 42.79 |
| CCTG | Cpf1 | 3 | Gal | 12 | 19.44 | 9.55 | 5.52 | 23.73 |
| CCTG | D10A | 3 | Gal | 12 | 2.60 | 0.37 | 0.22 | 0.93 |
| CCTG | eSpCas9 | 4 | Gal | 12 | 46.10 | 7.68 | 3.84 | 12.23 |
| CCTG | HF1 | 3 | Gal | 12 | 0.02 | 0.03 | 0.02 | 0.06 |
| CCTG | N863A | 3 | Gal | 12 | 0.17 | 0.10 | 0.06 | 0.26 |
| CCTG | SaCas9 | 3 | Gal | 12 | 0.16 | 0.03 | 0.02 | 0.08 |
| CCTG | SpCas9 | 6 | Gal | 12 | 22.69 | 26.14 | 10.67 | 27.43 |
| CCTG | Cpf1 | 6 | Gal | 24 | 40.08 | 7.16 | 2.92 | 7.51 |
| CCTG | D10A | 3 | Gal | 24 | 12.00 | 0.72 | 0.42 | 1.79 |
| CCTG | eSpCas9 | 4 | Gal | 24 | 85.48 | 7.15 | 3.57 | 11.37 |
| CCTG | HF1 | 3 | Gal | 24 | 0.02 | 0.02 | 0.01 | 0.05 |
| CCTG | N863A | 3 | Gal | 24 | 0.90 | 0.13 | 0.07 | 0.31 |
| CCTG | SaCas9 | 3 | Gal | 24 | 0.49 | 0.27 | 0.15 | 0.66 |
| CCTG | SpCas9 | 6 | Gal | 24 | 42.49 | 46.59 | 19.02 | 48.89 |
| CCTG | Cpf1 | 6 | Gal | 36 | 45.87 | 11.92 | 4.86 | 12.50 |
| CCTG | D10A | 3 | Gal | 36 | 14.80 | 3.70 | 2.14 | 9.19 |
| CCTG | eSpCas9 | 4 | Gal | 36 | 97.05 | 1.53 | 0.76 | 2.43 |
| CCTG | HF1 | 3 | Gal | 36 | 0.12 | 0.05 | 0.03 | 0.14 |
| CCTG | N863A | 3 | Gal | 36 | 1.20 | 0.04 | 0.02 | 0.10 |
| CCTG | SaCas9 | 3 | Gal | 36 | 2.19 | 1.10 | 0.63 | 2.73 |
| CCTG | SpCas9 | 3 | Gal | 36 | 96.63 | 1.16 | 0.67 | 2.88 |
| CCTG | Cpf1 | 6 | Glu | 0 | 0.42 | 0.14 | 0.06 | 0.15 |
| CCTG | D10A | 3 | Glu | 0 | 0.09 | 0.03 | 0.01 | 0.06 |
| CCTG | eSpCas9 | 3 | Glu | 0 | 0.04 | 0.05 | 0.03 | 0.12 |
| CCTG | HF1 | 3 | Glu | 0 | 0.03 | 0.01 | 0.00 | 0.02 |
| CCTG | N863A | 3 | Glu | 0 | 0.02 | 0.01 | 0.01 | 0.03 |
| CCTG | SaCas9 | 3 | Glu | 0 | 1.15 | 1.45 | 0.84 | 3.61 |
| CCTG | SpCas9 | 5 | Glu | 0 | 0.06 | 0.06 | 0.03 | 0.08 |
| CCTG | Cpf1 | 3 | Glu | 12 | 1.68 | 0.67 | 0.38 | 1.65 |
| CCTG | D10A | 3 | Glu | 12 | 0.24 | 0.08 | 0.05 | 0.21 |
| CCTG | eSpCas9 | 3 | Glu | 12 | 0.00 | 0.01 | 0.00 | 0.02 |
| CCTG | HF1 | 3 | Glu | 12 | 0.11 | 0.03 | 0.02 | 0.07 |
| CCTG | N863A | 3 | Glu | 12 | 0.10 | 0.09 | 0.05 | 0.24 |
| CCTG | SaCas9 | 3 | Glu | 12 | 0.36 | 0.26 | 0.15 | 0.64 |
| CCTG | SpCas9 | 2 | Glu | 12 | 0.35 | 0.16 | 0.11 | 1.40 |
| CCTG | Cpf1 | 6 | Glu | 24 | 0.89 | 0.93 | 0.38 | 0.98 |
| CCTG | D10A | 3 | Glu | 24 | 0.46 | 0.14 | 0.08 | 0.34 |
| CCTG | eSpCas9 | 3 | Glu | 24 | 0.01 | 0.02 | 0.01 | 0.04 |
| CCTG | HF1 | 3 | Glu | 24 | 0.11 | 0.02 | 0.01 | 0.05 |
| CCTG | N863A | 3 | Glu | 24 | 0.36 | 0.17 | 0.10 | 0.42 |
| CCTG | SaCas9 | 3 | Glu | 24 | 0.91 | 0.41 | 0.24 | 1.02 |
| CCTG | SpCas9 | 2 | Glu | 24 | 0.70 | 0.03 | 0.02 | 0.25 |
| CCTG | Cpf1 | 6 | Glu | 36 | 2.63 | 2.05 | 0.84 | 2.16 |
| CCTG | D10A | 3 | Glu | 36 | 0.65 | 0.11 | 0.07 | 0.28 |
| CCTG | eSpCas9 | 3 | Glu | 36 | 0.47 | 0.15 | 0.09 | 0.38 |
| CCTG | HF1 | 3 | Glu | 36 | 0.50 | 0.50 | 0.29 | 1.24 |
| CCTG | N863A | 3 | Glu | 36 | 0.55 | 0.24 | 0.14 | 0.60 |
| CCTG | SaCas9 | 3 | Glu | 36 | 1.79 | 0.50 | 0.29 | 1.25 |
| CCTG | SpCas9 | 5 | Glu | 36 | 0.95 | 1.01 | 0.45 | 1.26 |
| GCN | Cpf1 | 3 | Gal | 0 | 0.16 | 0.02 | 0.01 | 0.04 |
| GCN | D10A | 3 | Gal | 0 | 0.12 | 0.08 | 0.05 | 0.20 |
| GCN | eSpCas9 | 5 | Gal | 0 | 16.85 | 25.83 | 11.55 | 32.07 |
| GCN | HF1 | 4 | Gal | 0 | 0.30 | 0.33 | 0.16 | 0.52 |
| GCN | N863A | 3 | Gal | 0 | 0.07 | 0.03 | 0.02 | 0.08 |
| GCN | SaCas9 | 3 | Gal | 0 | 0.38 | 0.22 | 0.13 | 0.54 |
| GCN | SpCas9 | 3 | Gal | 0 | 0.13 | 0.12 | 0.07 | 0.29 |
| GCN | Cpf1 | 3 | Gal | 12 | 0.74 | 0.08 | 0.04 | 0.19 |
| GCN | D10A | 3 | Gal | 12 | 2.34 | 0.62 | 0.36 | 1.55 |
| GCN | eSpCas9 | 4 | Gal | 12 | 59.63 | 20.16 | 10.08 | 32.08 |
| GCN | HF1 | 4 | Gal | 12 | 0.68 | 0.44 | 0.22 | 0.69 |
| GCN | N863A | 3 | Gal | 12 | 0.27 | 0.09 | 0.05 | 0.23 |
| GCN | SaCas9 | 3 | Gal | 12 | 0.15 | 0.16 | 0.09 | 0.39 |
| GCN | SpCas9 | 3 | Gal | 12 | 31.33 | 6.21 | 3.59 | 15.43 |
| GCN | Cpf1 | 3 | Gal | 24 | 3.20 | 0.58 | 0.33 | 1.43 |
| GCN | D10A | 3 | Gal | 24 | 7.84 | 0.83 | 0.48 | 2.07 |
| GCN | eSpCas9 | 5 | Gal | 24 | 87.92 | 10.86 | 4.86 | 13.49 |
| GCN | HF1 | 4 | Gal | 24 | 3.71 | 1.27 | 0.63 | 2.02 |
| GCN | N863A | 3 | Gal | 24 | 1.32 | 0.17 | 0.10 | 0.43 |
| GCN | SaCas9 | 3 | Gal | 24 | 0.29 | 0.15 | 0.09 | 0.37 |
| GCN | SpCas9 | 3 | Gal | 24 | 84.53 | 1.46 | 0.85 | 3.64 |
| GCN | Cpf1 | 3 | Gal | 36 | 3.96 | 0.05 | 0.03 | 0.12 |
| GCN | D10A | 3 | Gal | 36 | 8.26 | 0.62 | 0.36 | 1.54 |
| GCN | eSpCas9 | 5 | Gal | 36 | 90.12 | 9.01 | 4.03 | 11.19 |
| GCN | HF1 | 4 | Gal | 36 | 5.42 | 0.68 | 0.34 | 1.08 |
| GCN | N863A | 3 | Gal | 36 | 2.06 | 0.29 | 0.16 | 0.71 |
| GCN | SaCas9 | 3 | Gal | 36 | 1.77 | 0.71 | 0.41 | 1.77 |
| GCN | SpCas9 | 3 | Gal | 36 | 92.10 | 4.09 | 2.36 | 10.15 |
| GCN | Cpf1 | 3 | Glu | 0 | 0.15 | 0.08 | 0.04 | 0.19 |
| GCN | D10A | 3 | Glu | 0 | 0.11 | 0.03 | 0.02 | 0.07 |
| GCN | eSpCas9 | 4 | Glu | 0 | 33.00 | 31.34 | 15.67 | 49.87 |
| GCN | HF1 | 4 | Glu | 0 | 0.74 | 0.33 | 0.17 | 0.53 |
| GCN | N863A | 3 | Glu | 0 | 0.03 | 0.03 | 0.02 | 0.08 |
| GCN | SaCas9 | 3 | Glu | 0 | 0.36 | 0.23 | 0.13 | 0.56 |
| GCN | SpCas9 | 3 | Glu | 0 | 0.39 | 0.23 | 0.13 | 0.56 |
| GCN | Cpf1 | 3 | Glu | 12 | 0.22 | 0.04 | 0.02 | 0.10 |
| GCN | D10A | 3 | Glu | 12 | 0.09 | 0.05 | 0.03 | 0.12 |
| GCN | eSpCas9 | 4 | Glu | 12 | 40.14 | 34.29 | 17.14 | 54.56 |
| GCN | HF1 | 4 | Glu | 12 | 0.23 | 0.16 | 0.08 | 0.25 |
| GCN | N863A | 3 | Glu | 12 | 0.16 | 0.21 | 0.12 | 0.51 |
| GCN | SaCas9 | 3 | Glu | 12 | 0.36 | 0.23 | 0.13 | 0.56 |
| GCN | SpCas9 | 3 | Glu | 12 | 0.23 | 0.09 | 0.05 | 0.23 |
| GCN | Cpf1 | 3 | Glu | 24 | 0.91 | 0.16 | 0.09 | 0.39 |
| GCN | D10A | 3 | Glu | 24 | 0.33 | 0.13 | 0.07 | 0.32 |
| GCN | eSpCas9 | 4 | Glu | 24 | 44.00 | 31.77 | 15.89 | 50.56 |
| GCN | HF1 | 4 | Glu | 24 | 0.21 | 0.33 | 0.16 | 0.52 |
| GCN | N863A | 3 | Glu | 24 | 0.31 | 0.08 | 0.05 | 0.20 |
| GCN | SaCas9 | 3 | Glu | 24 | 1.57 | 0.81 | 0.47 | 2.01 |
| GCN | SpCas9 | 3 | Glu | 24 | 0.40 | 0.30 | 0.17 | 0.73 |
| GCN | Cpf1 | 3 | Glu | 36 | 1.08 | 0.13 | 0.08 | 0.32 |
| GCN | D10A | 3 | Glu | 36 | 0.78 | 0.08 | 0.05 | 0.20 |
| GCN | eSpCas9 | 4 | Glu | 36 | 47.18 | 29.63 | 14.81 | 47.15 |
| GCN | HF1 | 4 | Glu | 36 | 1.54 | 0.28 | 0.14 | 0.44 |
| GCN | N863A | 3 | Glu | 36 | 0.96 | 0.24 | 0.14 | 0.59 |
| GCN | SaCas9 | 3 | Glu | 36 | 2.62 | 0.90 | 0.52 | 2.24 |
| GCN | SpCas9 | 3 | Glu | 36 | 1.51 | 0.66 | 0.38 | 1.63 |
| NR | Cpf1 | 4 | Gal | 0 | 3.81 | 3.52 | 1.76 | 5.60 |
| NR | D10A | 3 | Gal | 0 | 0.16 | 0.04 | 0.03 | 0.11 |
| NR | eSpCas9 | 3 | Gal | 0 | 1.17 | 0.51 | 0.29 | 1.26 |
| NR | HF1 | 3 | Gal | 0 | 0.06 | 0.01 | 0.00 | 0.02 |
| NR | ISCEI | 3 | Gal | 0 | 0.21 | 0.01 | 0.01 | 0.03 |
| NR | N863A | 3 | Gal | 0 | 0.10 | 0.01 | 0.01 | 0.03 |
| NR | SaCas9 | 4 | Gal | 0 | 0.09 | 0.03 | 0.02 | 0.05 |
| NR | SpCas9 | 6 | Gal | 0 | 0.18 | 0.15 | 0.06 | 0.15 |
| NR | Cpf1 | 4 | Gal | 12 | 36.48 | 4.76 | 2.38 | 7.58 |
| NR | D10A | 3 | Gal | 12 | 0.17 | 0.15 | 0.09 | 0.37 |
| NR | eSpCas9 | 3 | Gal | 12 | 5.16 | 0.45 | 0.26 | 1.12 |
| NR | HF1 | 3 | Gal | 12 | 0.22 | 0.12 | 0.07 | 0.31 |
| NR | ISCEI | 3 | Gal | 12 | 9.96 | 2.28 | 1.32 | 5.67 |
| NR | N863A | 3 | Gal | 12 | 0.06 | 0.02 | 0.01 | 0.04 |
| NR | SaCas9 | 4 | Gal | 12 | 3.60 | 3.39 | 1.70 | 5.40 |
| NR | SpCas9 | 6 | Gal | 12 | 42.57 | 18.43 | 7.52 | 19.34 |
| NR | Cpf1 | 4 | Gal | 24 | 83.90 | 3.48 | 1.74 | 5.54 |
| NR | D10A | 3 | Gal | 24 | 0.95 | 0.54 | 0.31 | 1.34 |
| NR | eSpCas9 | 3 | Gal | 24 | 17.80 | 0.72 | 0.42 | 1.79 |
| NR | HF1 | 3 | Gal | 24 | 0.50 | 0.02 | 0.01 | 0.04 |
| NR | ISCEI | 3 | Gal | 24 | 24.07 | 2.20 | 1.27 | 5.47 |
| NR | N863A | 3 | Gal | 24 | 0.10 | 0.08 | 0.05 | 0.20 |
| NR | SaCas9 | 4 | Gal | 24 | 36.98 | 12.37 | 6.18 | 19.68 |
| NR | SpCas9 | 6 | Gal | 24 | 85.78 | 4.26 | 1.74 | 4.47 |
| NR | Cpf1 | 4 | Gal | 36 | 92.60 | 2.94 | 1.47 | 4.68 |
| NR | D10A | 3 | Gal | 36 | 1.27 | 0.55 | 0.32 | 1.36 |
| NR | eSpCas9 | 3 | Gal | 36 | 18.87 | 1.10 | 0.64 | 2.74 |
| NR | HF1 | 3 | Gal | 36 | 0.90 | 0.28 | 0.16 | 0.69 |
| NR | ISCEI | 3 | Gal | 36 | 27.97 | 1.16 | 0.67 | 2.88 |
| NR | N863A | 3 | Gal | 36 | 0.46 | 0.18 | 0.10 | 0.45 |
| NR | SaCas9 | 4 | Gal | 36 | 54.53 | 4.94 | 2.47 | 7.86 |
| NR | SpCas9 | 6 | Gal | 36 | 86.95 | 3.04 | 1.24 | 3.19 |
| NR | Cpf1 | 4 | Glu | 0 | 6.54 | 1.71 | 0.85 | 2.72 |
| NR | D10A | 3 | Glu | 0 | 0.24 | 0.28 | 0.16 | 0.69 |
| NR | eSpCas9 | 3 | Glu | 0 | 1.58 | 0.38 | 0.22 | 0.93 |
| NR | HF1 | 3 | Glu | 0 | 0.20 | 0.19 | 0.11 | 0.48 |
| NR | ISCEI | 3 | Glu | 0 | 0.17 | 0.13 | 0.08 | 0.33 |
| NR | N863A | 3 | Glu | 0 | 0.05 | 0.04 | 0.02 | 0.10 |
| NR | SaCas9 | 4 | Glu | 0 | 0.14 | 0.06 | 0.03 | 0.09 |
| NR | SpCas9 | 6 | Glu | 0 | 0.62 | 1.06 | 0.43 | 1.11 |
| NR | Cpf1 | 4 | Glu | 12 | 10.79 | 2.81 | 1.40 | 4.47 |
| NR | D10A | 3 | Glu | 12 | 0.10 | 0.07 | 0.04 | 0.17 |
| NR | eSpCas9 | 3 | Glu | 12 | 2.50 | 0.59 | 0.34 | 1.47 |
| NR | HF1 | 3 | Glu | 12 | 0.09 | 0.09 | 0.05 | 0.21 |
| NR | ISCEI | 3 | Glu | 12 | 0.30 | 0.12 | 0.07 | 0.30 |
| NR | N863A | 3 | Glu | 12 | 0.01 | 0.01 | 0.01 | 0.02 |
| NR | SaCas9 | 4 | Glu | 12 | 0.03 | 0.02 | 0.01 | 0.03 |
| NR | SpCas9 | 6 | Glu | 12 | 0.23 | 0.23 | 0.10 | 0.25 |
| NR | Cpf1 | 4 | Glu | 24 | 15.25 | 4.00 | 2.00 | 6.36 |
| NR | D10A | 3 | Glu | 24 | 0.77 | 0.66 | 0.38 | 1.63 |
| NR | eSpCas9 | 3 | Glu | 24 | 4.16 | 0.55 | 0.32 | 1.36 |
| NR | HF1 | 3 | Glu | 24 | 0.28 | 0.07 | 0.04 | 0.18 |
| NR | ISCEI | 3 | Glu | 24 | 1.09 | 0.06 | 0.03 | 0.14 |
| NR | N863A | 3 | Glu | 24 | 0.15 | 0.03 | 0.02 | 0.07 |
| NR | SaCas9 | 4 | Glu | 24 | 0.22 | 0.08 | 0.04 | 0.13 |
| NR | SpCas9 | 6 | Glu | 24 | 0.70 | 0.38 | 0.16 | 0.40 |
| NR | Cpf1 | 4 | Glu | 36 | 17.00 | 4.29 | 2.14 | 6.82 |
| NR | D10A | 3 | Glu | 36 | 1.22 | 1.39 | 0.80 | 3.44 |
| NR | eSpCas9 | 3 | Glu | 36 | 5.79 | 0.30 | 0.17 | 0.75 |
| NR | HF1 | 3 | Glu | 36 | 0.93 | 0.17 | 0.10 | 0.43 |
| NR | ISCEI | 3 | Glu | 36 | 1.26 | 0.11 | 0.06 | 0.27 |
| NR | N863A | 3 | Glu | 36 | 0.47 | 0.10 | 0.06 | 0.26 |
| NR | SaCas9 | 4 | Glu | 36 | 0.80 | 0.13 | 0.06 | 0.20 |
| NR | SpCas9 | 6 | Glu | 36 | 1.89 | 1.15 | 0.47 | 1.21 |
| TGGAA | Cpf1 | 3 | Gal | 0 | 0.15 | 0.06 | 0.03 | 0.15 |
| TGGAA | D10A | 6 | Gal | 0 | 21.12 | 36.12 | 14.74 | 37.90 |
| TGGAA | eSpCas9 | 3 | Gal | 0 | 69.17 | 17.24 | 9.95 | 42.83 |
| TGGAA | HF1 | 3 | Gal | 0 | 0.12 | 0.11 | 0.07 | 0.28 |
| TGGAA | N863A | 3 | Gal | 0 | 0.18 | 0.04 | 0.02 | 0.10 |
| TGGAA | SaCas9 | 3 | Gal | 0 | 0.26 | 0.17 | 0.10 | 0.41 |
| TGGAA | SpCas9 | 3 | Gal | 0 | 1.93 | 0.81 | 0.47 | 2.02 |
| TGGAA | Cpf1 | 3 | Gal | 12 | 5.30 | 1.82 | 1.05 | 4.53 |
| TGGAA | D10A | 3 | Gal | 12 | 25.59 | 22.27 | 12.86 | 55.33 |
| TGGAA | eSpCas9 | 3 | Gal | 12 | 86.03 | 8.13 | 4.69 | 20.19 |
| TGGAA | HF1 | 3 | Gal | 12 | 0.69 | 0.13 | 0.07 | 0.31 |
| TGGAA | N863A | 3 | Gal | 12 | 9.94 | 2.29 | 1.32 | 5.68 |
| TGGAA | SaCas9 | 3 | Gal | 12 | 0.21 | 0.18 | 0.10 | 0.45 |
| TGGAA | SpCas9 | 3 | Gal | 12 | 54.23 | 5.15 | 2.97 | 12.79 |
| TGGAA | Cpf1 | 3 | Gal | 24 | 27.30 | 1.39 | 0.80 | 3.45 |
| TGGAA | D10A | 6 | Gal | 24 | 65.58 | 33.45 | 13.66 | 35.11 |
| TGGAA | eSpCas9 | 3 | Gal | 24 | 96.37 | 2.40 | 1.39 | 5.96 |
| TGGAA | HF1 | 3 | Gal | 24 | 2.58 | 0.19 | 0.11 | 0.48 |
| TGGAA | N863A | 3 | Gal | 24 | 43.10 | 3.87 | 2.24 | 9.62 |
| TGGAA | SaCas9 | 3 | Gal | 24 | 0.43 | 0.07 | 0.04 | 0.17 |
| TGGAA | SpCas9 | 3 | Gal | 24 | 92.23 | 0.68 | 0.39 | 1.69 |
| TGGAA | Cpf1 | 3 | Gal | 36 | 41.63 | 3.76 | 2.17 | 9.34 |
| TGGAA | D10A | 6 | Gal | 36 | 70.44 | 35.17 | 14.36 | 36.91 |
| TGGAA | eSpCas9 | 3 | Gal | 36 | 96.30 | 1.56 | 0.90 | 3.88 |
| TGGAA | HF1 | 3 | Gal | 36 | 4.15 | 0.05 | 0.03 | 0.13 |
| TGGAA | N863A | 3 | Gal | 36 | 46.13 | 3.39 | 1.96 | 8.43 |
| TGGAA | SaCas9 | 3 | Gal | 36 | 1.84 | 0.57 | 0.33 | 1.42 |
| TGGAA | SpCas9 | 3 | Gal | 36 | 95.63 | 1.78 | 1.03 | 4.42 |
| TGGAA | Cpf1 | 3 | Glu | 0 | 0.13 | 0.03 | 0.02 | 0.08 |
| TGGAA | D10A | 6 | Glu | 0 | 9.48 | 7.16 | 2.92 | 7.52 |
| TGGAA | eSpCas9 | 2 | Glu | 0 | 52.90 | 10.18 | 7.20 | 91.48 |
| TGGAA | HF1 | 3 | Glu | 0 | 0.05 | 0.02 | 0.01 | 0.04 |
| TGGAA | N863A | 3 | Glu | 0 | 0.17 | 0.09 | 0.05 | 0.22 |
| TGGAA | SaCas9 | 3 | Glu | 0 | 0.26 | 0.19 | 0.11 | 0.46 |
| TGGAA | SpCas9 | 3 | Glu | 0 | 1.88 | 0.83 | 0.48 | 2.06 |
| TGGAA | Cpf1 | 3 | Glu | 12 | 0.10 | 0.02 | 0.01 | 0.05 |
| TGGAA | D10A | 3 | Glu | 12 | 13.42 | 10.12 | 5.84 | 25.14 |
| TGGAA | eSpCas9 | 2 | Glu | 12 | 73.60 | 8.91 | 6.30 | 80.05 |
| TGGAA | HF1 | 3 | Glu | 12 | 0.12 | 0.03 | 0.02 | 0.08 |
| TGGAA | N863A | 3 | Glu | 12 | 0.13 | 0.03 | 0.02 | 0.07 |
| TGGAA | SaCas9 | 3 | Glu | 12 | 0.07 | 0.02 | 0.01 | 0.06 |
| TGGAA | SpCas9 | 3 | Glu | 12 | 4.32 | 1.37 | 0.79 | 3.40 |
| TGGAA | Cpf1 | 3 | Glu | 24 | 0.61 | 0.23 | 0.14 | 0.58 |
| TGGAA | D10A | 6 | Glu | 24 | 15.54 | 11.06 | 4.52 | 11.61 |
| TGGAA | eSpCas9 | 2 | Glu | 24 | 76.65 | 6.43 | 4.55 | 57.81 |
| TGGAA | HF1 | 3 | Glu | 24 | 0.68 | 0.08 | 0.04 | 0.19 |
| TGGAA | N863A | 3 | Glu | 24 | 0.40 | 0.17 | 0.10 | 0.42 |
| TGGAA | SaCas9 | 3 | Glu | 24 | 0.67 | 0.16 | 0.09 | 0.39 |
| TGGAA | SpCas9 | 3 | Glu | 24 | 6.22 | 1.44 | 0.83 | 3.59 |
| TGGAA | Cpf1 | 3 | Glu | 36 | 0.78 | 0.20 | 0.12 | 0.49 |
| TGGAA | D10A | 6 | Glu | 36 | 15.00 | 10.58 | 4.32 | 11.10 |
| TGGAA | eSpCas9 | 2 | Glu | 36 | 78.10 | 5.94 | 4.20 | 53.37 |
| TGGAA | HF1 | 3 | Glu | 36 | 0.64 | 0.06 | 0.03 | 0.15 |
| TGGAA | N863A | 3 | Glu | 36 | 0.91 | 0.29 | 0.17 | 0.73 |
| TGGAA | SaCas9 | 3 | Glu | 36 | 1.80 | 0.41 | 0.24 | 1.02 |
| TGGAA | SpCas9 | 3 | Glu | 36 | 9.50 | 0.89 | 0.51 | 2.20 |

**Supplemental Table S4. Summary of flow cytometry experiments quantifications.**
